# Rooting the tree of eukaryotes using ancient gene duplications

**DOI:** 10.64898/2026.11.20.768066

**Authors:** Chenxue Lin, Christopher Kay, Oliver Schick, Anja Spang, Gergely J. Szöllősi, Davide Pisani, Philip C. J. Donoghue, Tom A. Williams

## Abstract

Identifying the root of eukaryotic phylogeny is integral to attempts to infer the nature of the last eukaryotic common ancestor and elucidate the prokaryote-to-eukaryote transition. However, the position of the eukaryotic root has proven difficult to resolve because of a high level of sequence divergence separating eukaryotes from their closest prokaryotic relatives, which can induce phylogenetic error in root placement. To overcome these issues, we exploit a complementary source of information: ancient gene duplications that occurred prior to the last eukaryotic common ancestor (LECA). For duplicated genes, each paralogue can be used to root its sibling, providing information independent of prokaryotic outgroups that can be pooled across duplicate gene families using phylogenetic reconciliation. We identified 219 pre-LECA duplicate gene families, and reconciliation analyses based on these families rejected most candidate root positions on the eukaryote tree. Most analyses favoured a root on, or within, the Metamonada, although roots on Discoba, or a deep split between Opimoda and Diphoda, were also not rejected under some conditions. The signal for an early-diverging Metamonada comes from their basal position within each paralogue in ∼23-25% of duplicate gene trees, more than for other groups examined. We discuss the intriguing agreement between concatenation and reconciliation methods on the eukaryotic root, potential biases that might affect reconciliation analyses, and the implications of this credible set of roots for the nature of LECA and models of early eukaryotic evolution.

## Introduction

Eukaryotes constitute one of life’s major domains, encompassing animals, plants, fungi, algae and a wide diversity of microbial lineages. They share a characteristic cellular architecture including a nucleus, a dynamic cytoskeleton, an extensive endomembrane system and a mitochondrion (or mitochondrion-related organelle) that underpin the structural and functional complexity of eukaryotic cells. Understanding how this complexity arose requires determining the position of the eukaryotic root, but published analyses have proposed a range of root positions, leading to contrasting inferences about the physiology and metabolic capabilities of the last eukaryotic common ancestor (LECA) (Figure 1A; Baldauf, 2003; Cavalier-Smith, 2010; Cerón-Romero et al., 2022; Derelle et al., 2015; He et al., 2014; Lamża, 2026; Rogozin et al., 2009; Stechmann & Cavalier-Smith, 2002; Al Jewari and Baldauf 2023; Torruella et al., 2025; Williamson et al., 2025). The major challenge in rooting the eukaryotic tree is that phylogenetic methods typically require an outgroup to resolve the root position, yet the closest available outgroups—Asgard archaea for archaeal-derived genes and alphaproteobacteria for mitochondrial genes—are evolutionarily distant from the eukaryotic crown group (Figure 1B). Their inclusion results in a long branch separating the ingroup and the outgroup that can distort within-eukaryote relationships and promote long-branch attraction (LBA), potentially pulling fast-evolving lineages such as metamonads toward the outgroup (Al Jewari & Baldauf, 2023; Derelle & Lang, 2012; Philippe et al., 2011; Williams et al., 2020). Although available better-fitting, site-heterogeneous substitution models and the use of slowly-evolving outgroups can partially mitigate long-branch attraction (LBA), these approaches do not guarantee complete elimination of this systematic bias (Philippe et al., 2005; Philippe & Roure, 2011).

**Figure 1.**
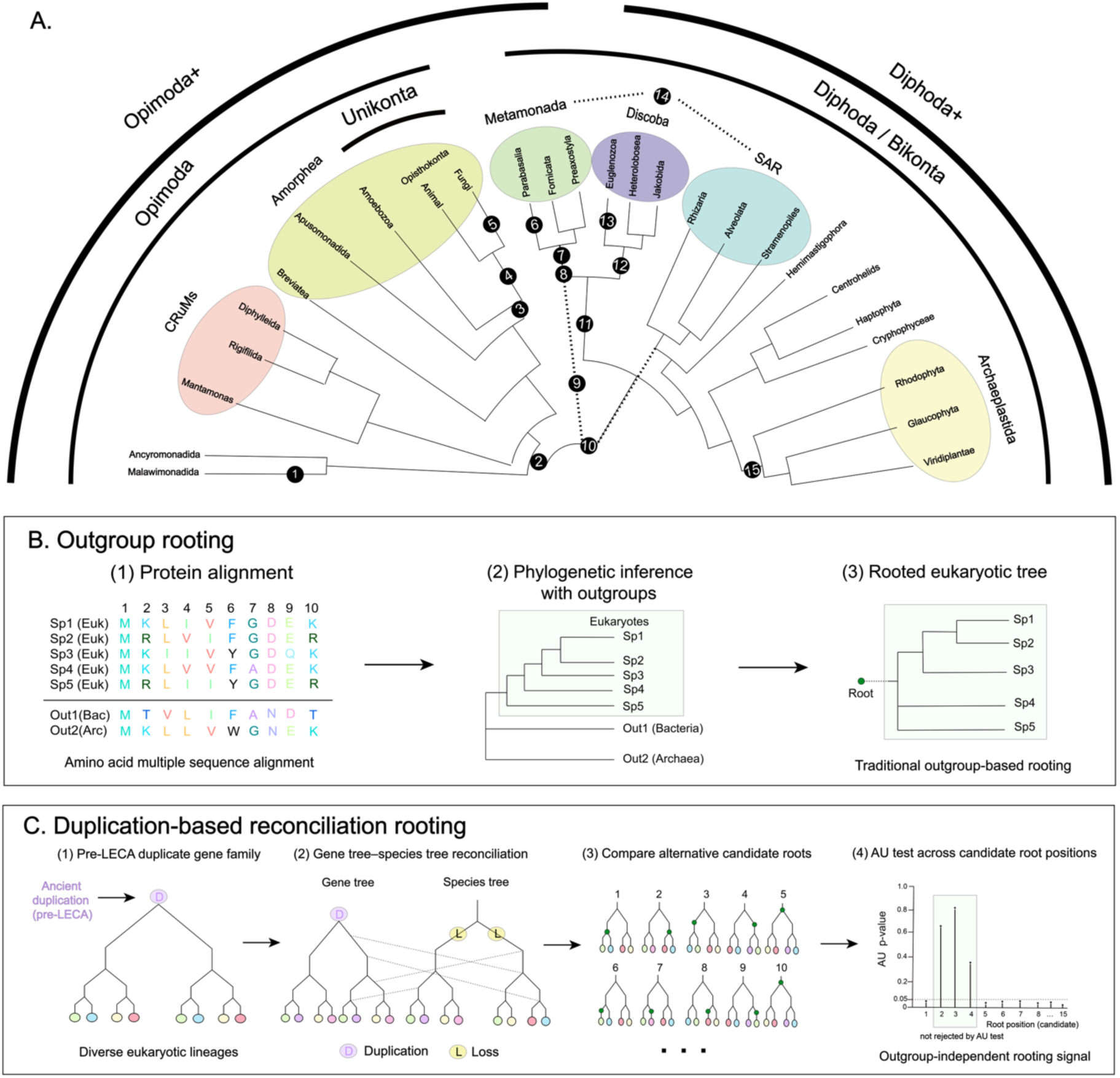
Overview of candidate eukaryotic rooting hypotheses and rooting strategies compared in this study. **(A)** Schematic overview of major eukaryotic lineages showing the 15 alternative rooting hypotheses evaluated in this study. Black numbered circles indicate the candidate rooting hypotheses considered in the duplication-based rooting analyses. Coloured regions mark major eukaryotic clades, including Amorphea, Metamonada, Discoba, SAR, Archaeplastida, and CRuMs (Collodictyonids, Rigifilida, and Mantamonas). The tested hypotheses comprise Malawimonadida (1; Cavalier-Smith, 2022), Opimoda–Diphoda (2; Derelle et al., 2015), Unikonta–Bikonta (3; Derelle & Lang, 2012; Stechmann & Cavalier-Smith, 2002), Opisthokonta (4; Cerón-Romero et al., 2022; Katz, 2012), Fungi (5; Cerón-Romero et al., 2022), Parabasalia (6; Al Jewari & Baldauf, 2023), Metamonada (7; Cerón-Romero et al., 2022), Metamonada-OpiDiph (8), Opimoda+/Diphoda+ (OpiDip+) (9; Williamson et al., 2025), Metamonada + Discoba + SAR (10), Metamonada + Discoba (11), Discoba (12; He et al., 2014), Euglenozoa (13; Cavalier-Smith, 2010), Metamonada + SAR (14), and Plantae (15; Rogozin et al., 2009). Hypotheses 8, 10, 11, and 14 represent additional configurations tested here to explore alternative placements of Metamonada relative to other major eukaryotic lineages, motivated by recent phylogenomic analyses (Torruella et al., 2025; Williamson et al., 2025). Auxiliary dotted connectors are used for hypotheses involving alternative combinations or placements of major lineages that cannot be represented by a single edge on the schematic backbone. This schematic is intended only as a guide to the hypotheses tested and does not imply a preferred root, basal lineage, or deep bipartition among eukaryotes. Some hypotheses were represented by more than one rooted species tree in downstream analyses because alternative species-tree backbones were also evaluated. **(B)** Schematic illustration of outgroup rooting, in which an archaeal or bacterial outgroup is used to root the eukaryotic tree. **(C)** Schematic illustration of duplication-based reconciliation rooting, in which pre-LECA duplicate gene families are reconciled across alternative candidate rooting hypotheses. Green dots indicate candidate root placements, and approximately unbiased (AU) tests are used to identify rooting hypotheses that cannot be rejected.

An alternative strategy is to infer the root without using external outgroups, by exploiting other sources of information (Katz 2012; Cerón-Romero et al. 2022). One such source can be found in eukaryotic gene families inferred to have undergone duplication before LECA, that is, families in which at least two paralogues were already present in LECA. Because these duplicated paralogous regions allow internal paralogous comparisons, they can help avoid reliance on the long branch separating eukaryotes from prokaryotic outgroups and provide rooting information that is difficult to obtain from outgroups alone. In principle, each of these families can inform the position of the eukaryotic root because each paralogue can serve as an internal outgroup for the other, without the need to include long-branching archaeal or bacterial outgroups (Dayhoff et al. 1972; Figure 1C). A similar approach has been used to root the entire tree of life based on gene families that duplicated prior to the divergence between Archaea and Bacteria (Brown & Doolittle, 1995; Gogarten et al., 1989; Iwabe et al., 1989; Zhaxybayeva & Gogarten, 2004). However, individual gene families often contain limited information about ancient divergences (Rokas & Carroll, 2005; Shen et al., 2017; Williams et al., 2020), such that information needs to be combined from many gene families in order to draw robust conclusions about the eukaryotic root. One approach is to concatenate duplicate gene families to boost signal (Moody et al., 2024), but since there is no natural matching of paralogues resulting from independent duplications, the number of possible equivalent ways to concatenate families rises exponentially with the number of families analyzed. This problem is likely to be particularly acute for the pre-LECA duplicate families considered here, because many eukaryotic gene families have more complex histories of duplication and retention, making it difficult to identify congruent sets of paralogues that could be concatenated consistently across families. An appealing alternative for combining root information from many pre-LECA duplicate families is gene tree-species tree reconciliation. Reconciliation methods map gene trees onto a species tree by invoking gene duplication, transfer and loss events, with the parameters of the reconciliation model (the rates of each type of event) inferred from the data, providing a natural framework to combine information on the root of the eukaryotic tree from many families. In particular, the joint likelihood in reconciliation can be evaluated across all pre-LECA duplicate families, allowing the individually weak rooting signal contained in each family to be combined into an overall statistical comparison of the support for alternative root positions. Work on the archaeal and bacterial domains has shown that gene- and species-tree reconciliation provides such information when outgroups are too distant (Williams et al. 2017; Coleman et al. 2021; Huang et al. 2025; Emms and Kelly 2017). However, previous reconciliation-based attempts to root the eukaryotic tree (Katz 2012; Cerón-Romero et al. 2022) have generally relied on relatively simple reconciliation approaches, such as gene tree parsimony or branch-homogeneous DTL models, which may not fully capture the substantial heterogeneity of eukaryotic genome evolution, potentially affecting root inference as seen for the archaeal tree of life (Huang et al., 2025).

Here, we perform a reconciliation-based analysis of the eukaryotic root, using a new reconciliation algorithm implemented in AleRax v1.4.0 (Morel et al., 2024) that allows rates of gene duplication, transfer and loss to vary across the branches of the species tree. To ameliorate issues that might arise from variation in genome and gene family completeness, we perform our reconciliation analyses on that subset of widely conserved eukaryotic gene families that we predict will contain direct information about the root: i.e. those families that underwent gene duplications prior to the last eukaryotic common ancestor (LECA). We identified 219 such pre-LECA duplicate gene families, and used these for gene tree-species tree reconciliations to investigate the position of the eukaryotic root.

## Results and Discussion

### Identification of pre-LECA duplicate families

Building on a recently developed pipeline (Kay et al., 2025), we identified 219 gene families containing pre-LECA duplication nodes from an analysis of 19,630 cross-domain gene family trees that include bacterial, archaeal, and eukaryotic sequences (Supplementary Table S1). Following previous studies (Tria et al., 2021; Vosseberg & Snel, 2017), a duplication node was classified as pre-LECA when the duplication was inferred to have occurred on the eukaryotic stem, after the divergence of eukaryotic sequences from their archaeal and bacterial homologues but before the diversification of crown eukaryotes, and both of its descendant branches contained representatives from multiple eukaryotic supergroups. This requirement distinguishes the pre-LECA duplications analysed here from older duplications predating the separation of eukaryotic, archaeal, or bacterial homologues. As shown in our workflow (Figure 2), we reanalysed these candidate gene families using a curated dataset of 131 representative eukaryotic species (Supplementary Table S2). To maximize phylogenetic breadth across major eukaryotic supergroups, representative species were selected based on recent large-scale phylogenomic analyses (e.g. Williamson et al., 2025; Torruella et al., 2025). Sampling emphasized balanced representation across major clades while minimizing taxonomic redundancy within densely sampled lineages to reduce potential biases arising from uneven representation. Predicted proteomes for selected taxa were retrieved from publicly available genome assemblies deposited in NCBI (O’Leary et al., 2016) or the EukProt v3 database (Richter et al., 2020). When multiple assembly versions were available, we prioritized the most complete and well-annotated datasets.

**Figure 2.**
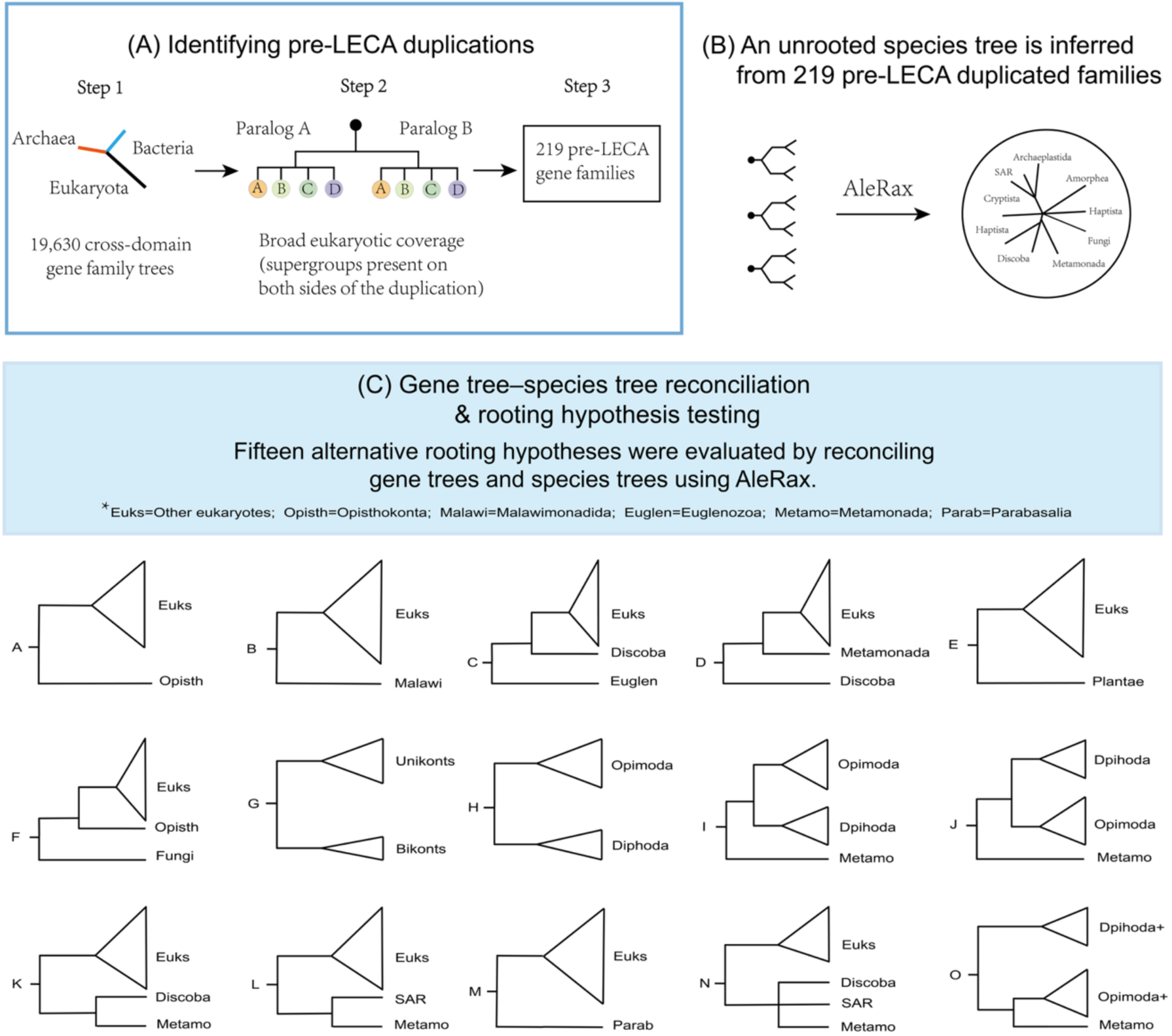
Workflow for identifying pre-LECA duplications, reconstructing an unrooted eukaryotic species tree, and testing alternative root hypotheses using gene tree–species tree reconciliation. **(A)** Candidate pre-LECA duplications were identified from 19,630 cross-domain gene family trees by requiring broad eukaryotic representation on both sides of a duplication node. **(B)** An unrooted species tree was inferred from 219 retained pre-LECA duplicate gene families. **(C)** Fifteen alternative rooting hypotheses were evaluated using gene tree–species tree reconciliation in AleRax. Some hypotheses were tested on alternative species-tree backbones, resulting in 18 rooted configurations for the full 129-taxon dataset and 12 for the reduced 117-taxon dataset.

The 219 identified gene families are distributed across all major eukaryotic supergroups, including Amorphea, Archaeplastida, SAR, Discoba, and Metamonada. Functional annotation of these gene families (Supplementary Table S3) shows that they are primarily associated with core cellular functions, such as information processing, cytoskeleton organization, membrane trafficking, mitochondrial and metabolic processes, signal transduction and regulation, and protein quality control (Supplementary Fig. S1). Among these pre-LECA duplicate gene families, those related to information processing and membrane trafficking account for the largest proportion.

These 219 pre-LECA duplicate gene families formed the basis for subsequent phylogenetic analyses, including species-tree inference and evaluation of alternative eukaryotic rooting hypotheses based on gene tree–species tree reconciliation. To explore the impact of substitution model fit on downstream analyses, we inferred gene-tree sets for these families under the single-matrix, C20–PMSF and C60–PMSF frameworks.

### Species tree inference

We inferred a species tree of the 131 representative eukaryotic taxa from the 219 high-confidence pre-LECA duplicate gene families using AleRax v1.4.0 (Morel et al., 2024) on the single-matrix ML gene trees. Gene families were analysed as separate multi-copy gene trees under the AleRax gene tree–species tree reconciliation framework, rather than being split into paralogue subsets or concatenated. AleRax searched for the rooted species tree that maximised the combined reconciliation likelihood across all gene families under the DTL model, using hill-climbing subtree-prune-and-regraft (SPR) moves. Because tree search under branch-heterogeneous models is computationally intractable, this analysis was performed under a branch-homogeneous model in which the same duplication, transfer and loss rates apply to all branches of the tree. The complete list of sampled taxa, dataset identifiers, and data sources is provided in Supplementary Table S2. Most internal branches of the original AleRax-inferred topology received strong support (Local Posterior Probability, LPP ≥ 0.95), indicating strong support for many of the inferred relationships under this reconciliation model. The recovered topology is broadly consistent with established eukaryotic relationships, with major clades such as Discoba, SAR, and Opisthokonta forming well-supported monophyletic groups. However, Metamonada was not recovered as monophyletic, and Archaeplastida was also split in the single-matrix tree. A small number of deep divergences received only moderate support (0.5 ≤ LPP < 0.9), likely reflecting limited phylogenetic signal or gene-tree discordance at these ancient nodes. Repeating the species-tree search with the C20–PMSF and C60–PMSF ML gene trees recovered broadly similar major clades, although the placements of several deep-branching lineages varied among frameworks. Metamonada remained non-monophyletic in both analyses, whereas Archaeplastida was non-monophyletic under C20–PMSF but monophyletic under C60–PMSF. Overall, the inferred species tree provides a robust framework for subsequent comparative and evolutionary analyses. The original single-matrix-based AleRax species tree and its LPP support values are provided in Supplementary Fig. S2, and the corresponding species-tree files, including the single-matrix, C20–PMSF and C60–PMSF AleRax trees and the final modified tree used for downstream reconciliation analyses, are provided in Supplementary Data 1.

In the unconstrained single-matrix-based species tree, the apusomonad taxa *Thecamonas trahens* and *Manchomonas bermudensis* were placed together with Centrohelida. Although this topology received strong statistical support, it is inconsistent with previous phylogenomic studies placing apusomonads and breviates within or close to Obazoa/Apusozoa rather than with Centrohelida (Brown et al. 2013; Cavalier-Smith et al. 2014; Adl et al. 2019; Torruella et al. 2023). We therefore manually grouped *Thecamonas*, *Manchomonas*, and *Pygsuia biforma* together as a clade in the species tree to maintain consistency with previous phylogenomic analyses. For the downstream rooting analyses, Metamonada and Archaeplastida were also treated as monophyletic in the modified reference topology. Because the topology was modified, the original AleRax LPP values were not transferred to the modified topology. Instead, quartet concordance for the final displayed topology (Figure 3) was assessed using ASTRAL-Pro3 quartet proportions (q1), by scoring the fixed modified topology against the same 219 multi-copy gene trees. This modified species tree served as the baseline topology for downstream analyses, with additional species-tree configurations evaluated for rooting hypotheses involving alternative relationships among major eukaryotic lineages. Under the default global DTL model, AleRax returned a maximum-likelihood (ML) root on the branch separating Malawimonadida + Ancyromonadida + Hemimastigophora from all other eukaryotes. This default root provides a useful reference point, but the final assessment of rooting hypotheses was carried out using UFBoot gene trees under both branch-homogeneous and branch-heterogeneous DTL models, with the latter including unrestricted and rate-bounded analyses and yielding higher total reconciliation log-likelihoods than the corresponding branch-homogeneous models on the same rooted species trees (Supplementary Table S4).

**Figure 3.**
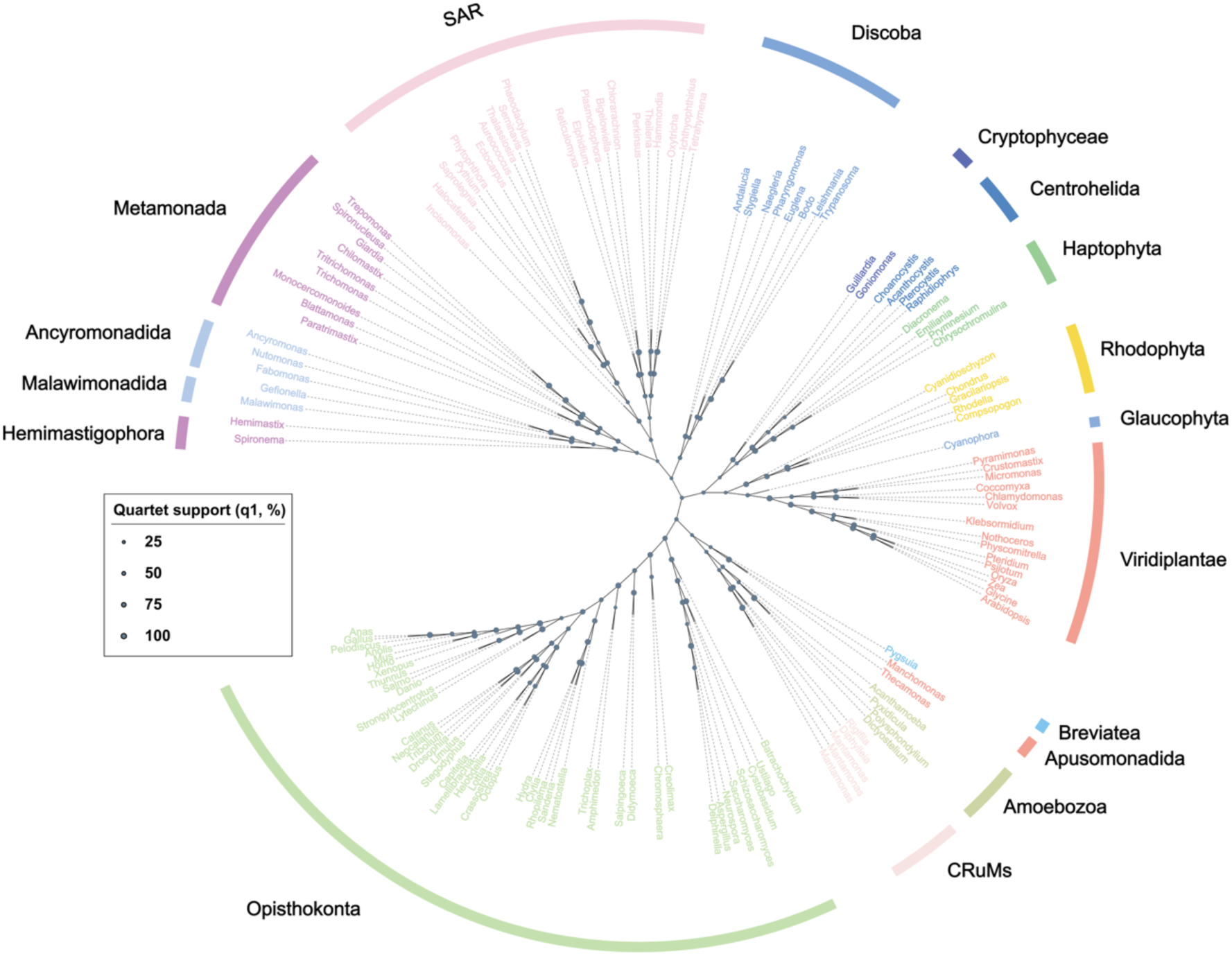
Unrooted species tree of 131 representative eukaryotic taxa inferred from 219 pre-LECA duplicate gene families. The initial species tree was inferred using AleRax v1.4.0 (Morel et al., 2024) from single-matrix maximum-likelihood gene trees of 219 gene families containing one or more ancient gene duplications predating LECA. The topology shown is the modified species tree used as the baseline topology for downstream reconciliation analyses. Circle size is proportional to ASTRAL-Pro3 quartet support (q1, %), calculated by scoring the fixed modified topology against the same 219 multi-copy gene trees. Major eukaryotic clades are colour-coded and labelled directly on the tree. Terminal labels are shortened to genus names for readability; full taxon names and lineage assignments are provided in Supplementary Table S2. The topology does not imply a preferred root position.

### Reconciliation of pre-LECA gene duplicates rejects most candidate rooting hypotheses on the eukaryotic tree

To evaluate alternative hypotheses for the position of the eukaryotic root, we performed reconciliations of the 219 pre-LECA duplicate gene families or subsets thereof using AleRax under duplication–transfer–loss (DTL) models (Szöllõsi et al. 2013; Morel et al. 2024). Because gene-tree uncertainty can affect reconciliation-based root inference, we used UFBoot gene-tree replicates for the reconciliation analyses. For the final rooting analyses, we excluded the two Hemimastigophora taxa to reduce uncertainty associated with their phylogenetic placement, leaving 129 taxa for the comparison of 18 rooted species-tree topologies. An additional 12 topologies were tested using a reduced dataset of 117 taxa, excluding Metamonada and selected additional taxa to assess the sensitivity of root inference to taxon sampling. For these candidate rooted topologies, we calculated reconciliation log-likelihoods and assessed support using AU tests based on per-gene-family reconciliation log-likelihoods separately for the two datasets (Figure 4; Supplementary Tables S5, S7, and S8). The 15 original rooting hypotheses comprised the Fungi root, Opimoda–Diphoda root, Plantae root, Discoba root, Euglenozoa root, Opisthokonta root, Metamonada root, Unikonta–Bikonta root, Malawimonadida root, Parabasalia root, Opimoda+/Diphoda+ (OpiDip+) hypothesis, Metamonada-OpiDiph hypothesis, and three alternative topologies involving Metamonada + Discoba, Metamonada + SAR, and Metamonada + Discoba + SAR. We included the Metamonada-OpiDiph hypothesis because previous analyses using mitochondrial genes to root eukaryotes have sometimes not included metamonads (Derelle and Lang, 2012; Derelle et al., 2015), or have included only a limited sample of their diversity (Williamson et al., 2025) due to mitochondrial reductive evolution and gene loss in this clade. We therefore reasoned that published support for the Opimoda–Diphoda root from those analyses might, in principle, also be compatible with a still earlier-diverging position for metamonads. The three alternative Metamonada-related topologies were included to explore alternative placements of Metamonada relative to Discoba and SAR in light of recent phylogenomic analyses (Torruella et al., 2025; Williamson et al., 2025).

**Figure 4.**
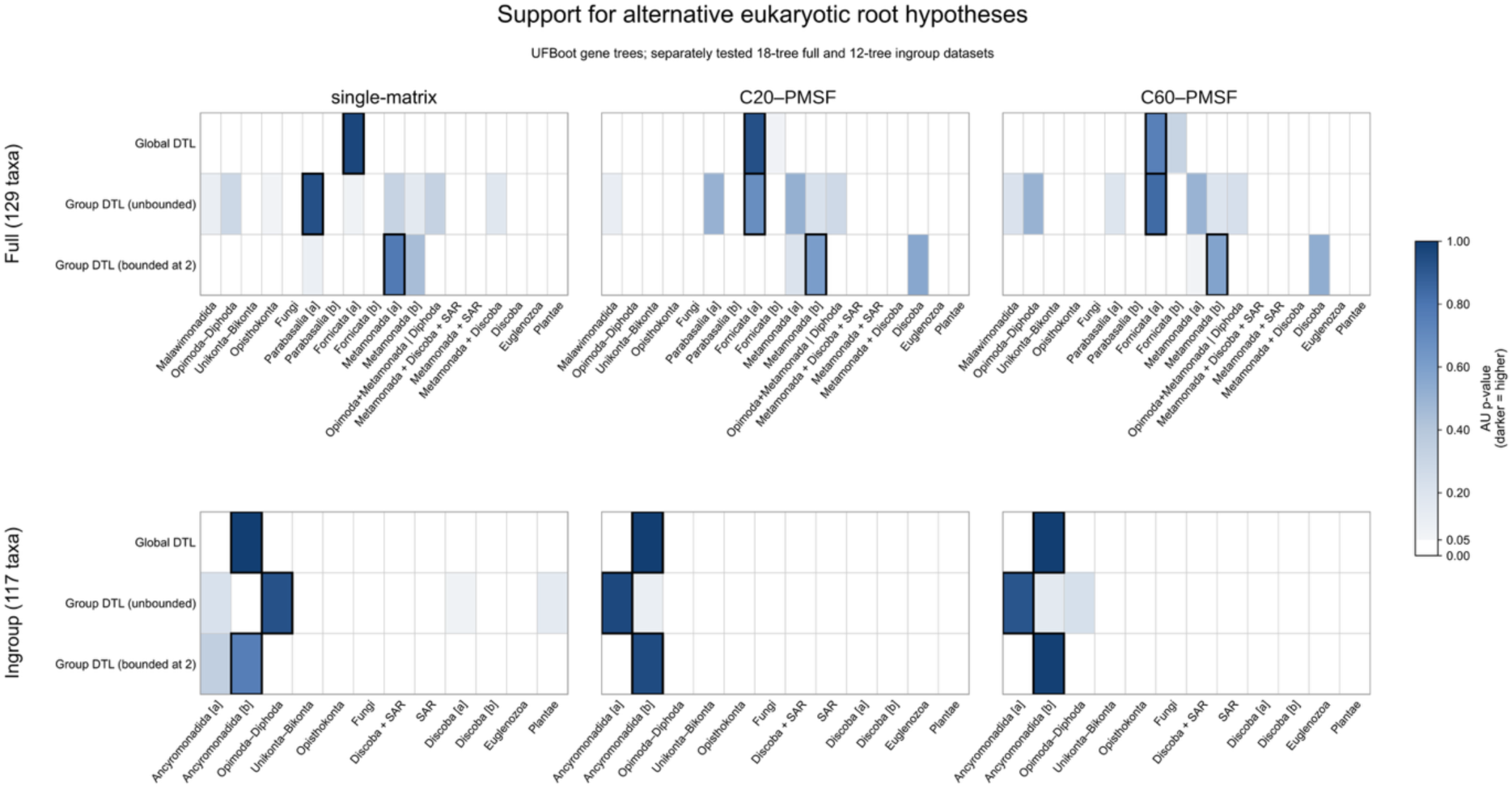
UFBoot-based AU-test support for alternative eukaryotic root hypotheses across gene-tree and reconciliation-model combinations. Heatmaps show approximately unbiased (AU) test p-values for 18 rooted species-tree configurations in the 129-taxon dataset (top) and 12 configurations in the 117-taxon ingroup dataset (bottom), tested separately. Columns represent single-matrix, C20– PMSF and C60–PMSF gene-tree frameworks, and rows represent global DTL, unbounded lineage-specific DTL and lineage-specific DTL with rates bounded at 2. Rooting configurations rejected by the AU test (P < 0.05) are shown in white, whereas unrejected configurations (P ≥ 0.05) are shaded blue, with darker shading indicating higher AU p-values. Black outlines indicate the highest reconciliation log-likelihood within each dataset, model and framework. Labels [a] and [b] distinguish different species-tree backbones sharing the same candidate root. Metamonada [a] also represents the Metamonada-OpiDiph hypothesis, whereas Metamonada [b] represents an alternative Metamonada-rooted topology. **Vertically-evolving gene families, and families with longer alignments, tend to favour metamonad roots**

We first performed a range of analyses using UFBoot gene trees inferred under either site-homogeneous single-matrix or site-heterogeneous substitution models as input to reconciliation analysis, and using both branch-homogeneous and branch-heterogeneous reconciliation models to infer the root position on the species tree. The branch-heterogeneous analyses included lineage-specific DTL models (on 12 named groups). During these analyses, we noticed that parameter estimates sometimes reached the default maximum boundary of 2 in AleRax; we therefore also performed analyses with unbounded rates. Overall, site-heterogeneous substitution models showed improved fit relative to the corresponding single-matrix analyses (Supplementary Table S6), and branch-heterogeneous reconciliation models fit the data better than branch-homogeneous models (Supplementary Table S4), as might be expected given the age of the eukaryotic root and the variation in evolutionary rates and modes across eukaryotes. We therefore focus on results from the site-heterogeneous C20–PMSF and C60–PMSF gene-tree frameworks under branch-heterogeneous DTL models in the following discussion, while presenting the complete set of analyses in the Supplementary Information.

The overall result of these analyses (Figure 4, Supplementary Tables S7 and S8) is that most candidate root hypotheses (Figure 1) were rejected by the approximately unbiased (AU) test at P < 0.05. In the full 129-taxon dataset, the roots within the remaining set of hypotheses not rejected by the AU test with the highest likelihoods included roots on, or within, Metamonada (in particular, Fornicata and Parabasalia), consistent with previous concatenation analyses identifying early-diverging Metamonada lineages (Al Jewari & Baldauf, 2023), with several other root hypotheses also remaining unrejected by the AU test under some model configurations, including a root between Metamonada and Opimoda-Diphoda (Metamonada-OpiDiph), a root between Opimoda-Diphoda with metamonads branching with Discoba (cf. Derelle et al., 2015), or on Discoba (He et al., 2014; Supplementary Tables S7 and S8). In the reduced 117-taxon ingroup dataset, Ancyromonadida-rooted configurations on the reduced candidate species trees had the highest likelihoods in eight of nine model configurations (Figure 4, Supplementary Tables S7 and S8).

Analysis of the individual paralogue family trees showed that a significant contributor of the phylogenetic signal for roots within, or near, Metamonada appears to be that these taxa branch at the base of pre-LECA paralogues more frequently than do other groups (in the site-heterogeneous C60-PMSF trees, median 25%, range 8-33%, compared to a median of ∼0.7%, range 0-7% for other groups), with families in which metamonads branch basal tending to favour roots on or within metamonads (partial Spearman’s ρ = 0.34–0.42 across nine gene-tree and reconciliation-model combinations, all P < 0.001; n = 200 families; Supplementary Table S9). However, the most frequent metamonad groups at the base of paralogues (61% of cases) include sequences from only one sub-lineage (Fornicata, Parabasalia or Preaxyostyla). Thus, to explore the nature of the phylogenetic signal for the identified root region in more detail, we next investigated the properties of families that favoured particular roots.

The fit between model and data is known to be important for accurate phylogenetic inference, especially for difficult problems (Williams et al. 2021, Kapli et al. 2021). In the context of the eukaryotic root, disagreements between analyses favouring metamonad or Discoban roots and those favouring Opimoda-Diphoda roots have often centred on the potential for the fast-or peculiarly-evolving sequences of metamonads to be drawn to the prokaryotic outgroup by long branch attraction (Derelle and Lang 2012; Derelle et al. 2015). Since our analyses on the full dataset also seem to favour metamonad roots (while not rejecting a deep Opimoda-Diphoda split, or indeed a root on Opimoda-Diphoda under certain conditions), we sought to investigate the nature of the support for different roots in more detail. One approach to evaluate conflicting results is to analyse the data using a better-fitting model, as in the site-heterogeneous substitution and branch-heterogeneous reconciliation analyses described above. A complementary approach is to filter the data to identify a subset that is easier to model, and might therefore provide more reliable phylogenetic information given the limitations of current models. This is the motivation that underlies the filtering of alignment sites by rate or composition in standard phylogenetic analyses (Mongiardino Koch, 2021; Philippe et al., 2011; Rodríguez-Ezpeleta et al., 2007; Williams et al., 2021).

We ranked gene families using a range of gene-family metrics, including species coverage, inferred transfer count, copy number per represented genome, composition-failure rate, verticality, alignment length, tip-normalised alignment length, and gene-tree shape (S/I and S/L). Verticality was defined as V/(V+D+T+L), where V represents inferred events of vertical descent and D, T and L represent duplication, transfer and loss events, respectively. The tree-shape metrics S/I and S/L describe the proportion of total internal branch length (I) or total tree length (L) accounted for by the longest internal backbone path (S), respectively (Figure 5A). Gene-family rankings are provided in Supplementary Table S10. Reconciliation-derived metrics were averaged across tested rooting configurations within each gene-tree and DTL-model combination. We then compared rooting support between high- and low-scoring gene-family subsets in the full 129-taxon and reduced 117-taxon datasets (Figure 5; Supplementary Table S11).

**Figure 5.**
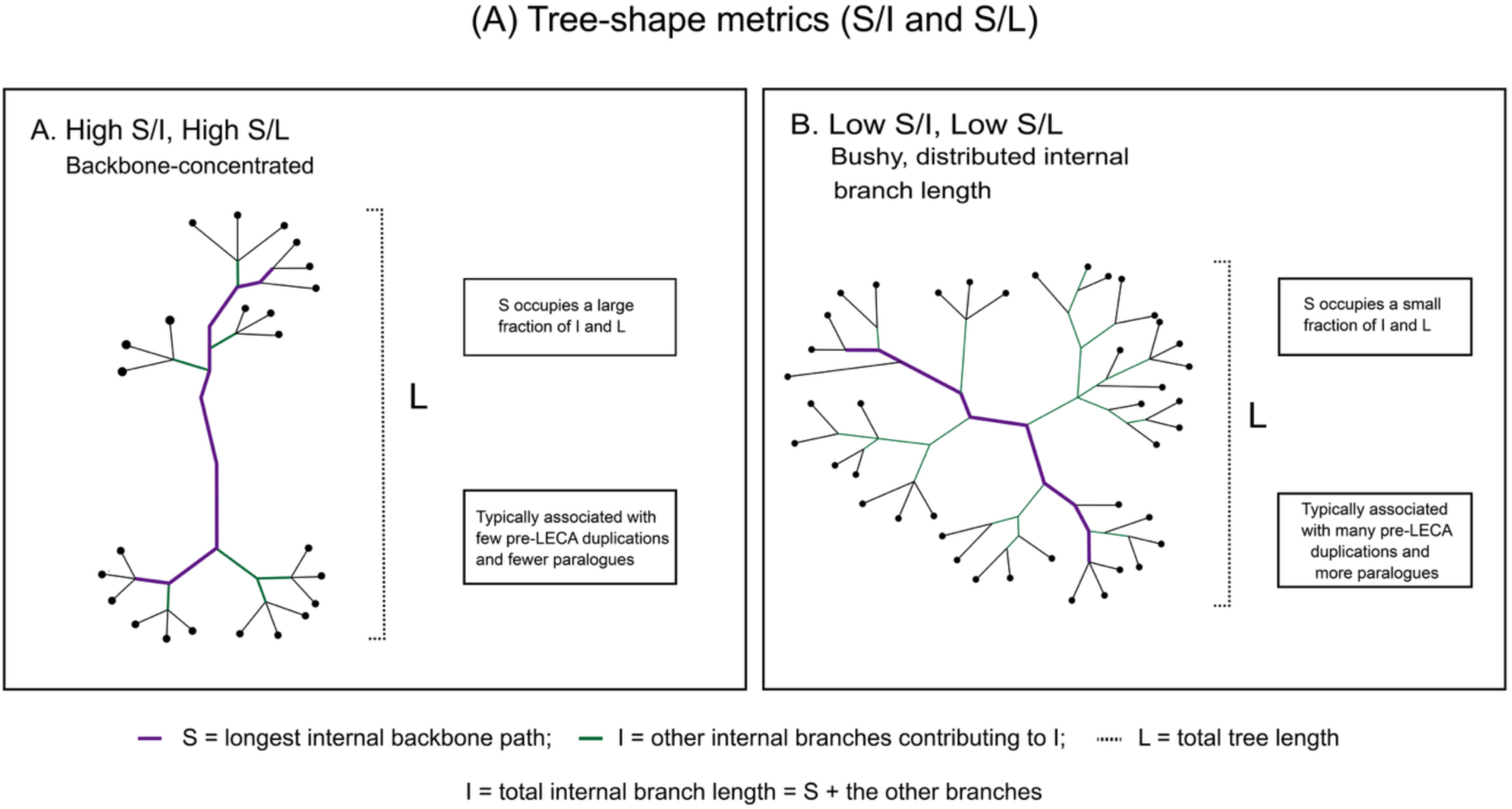
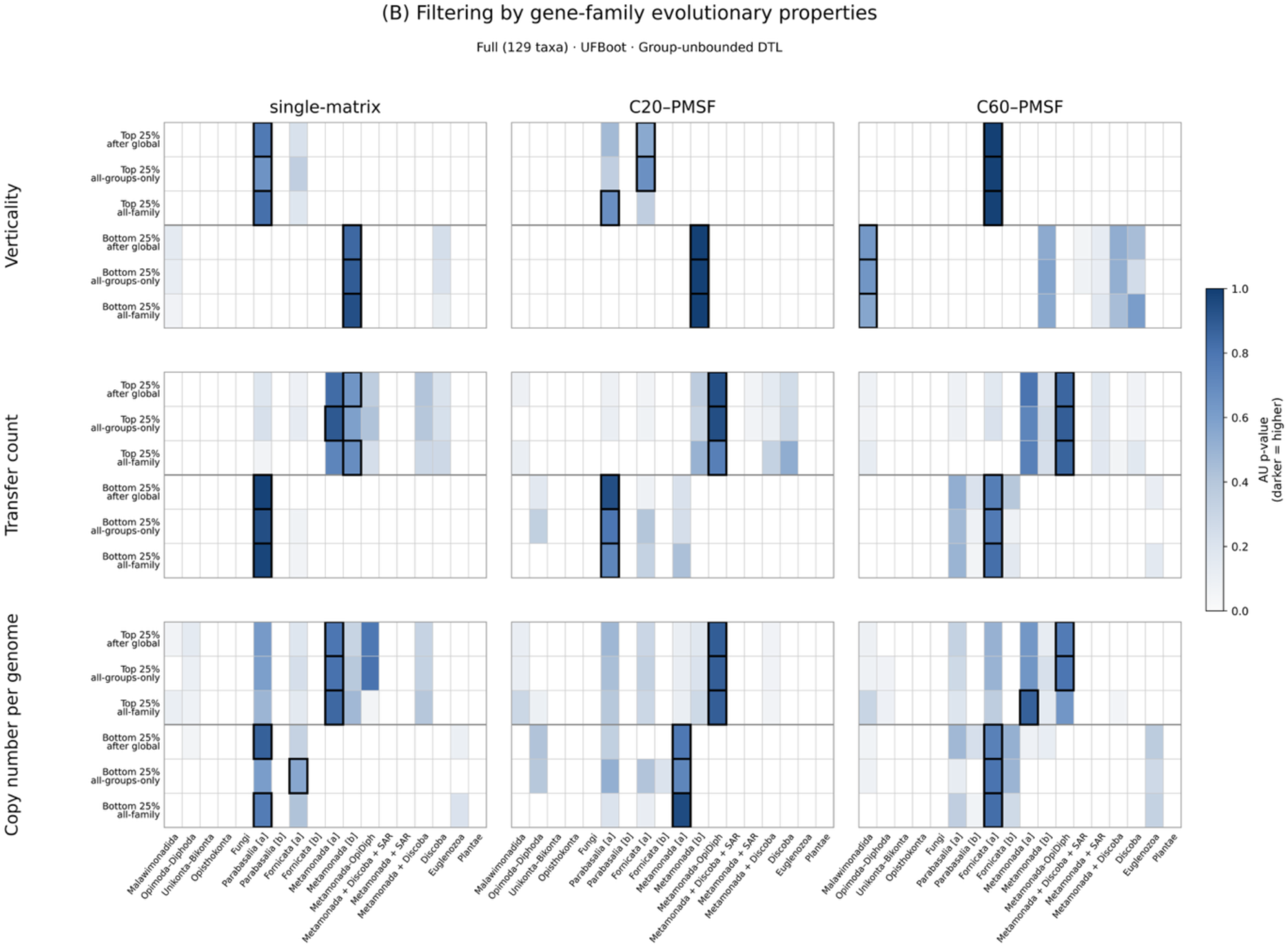
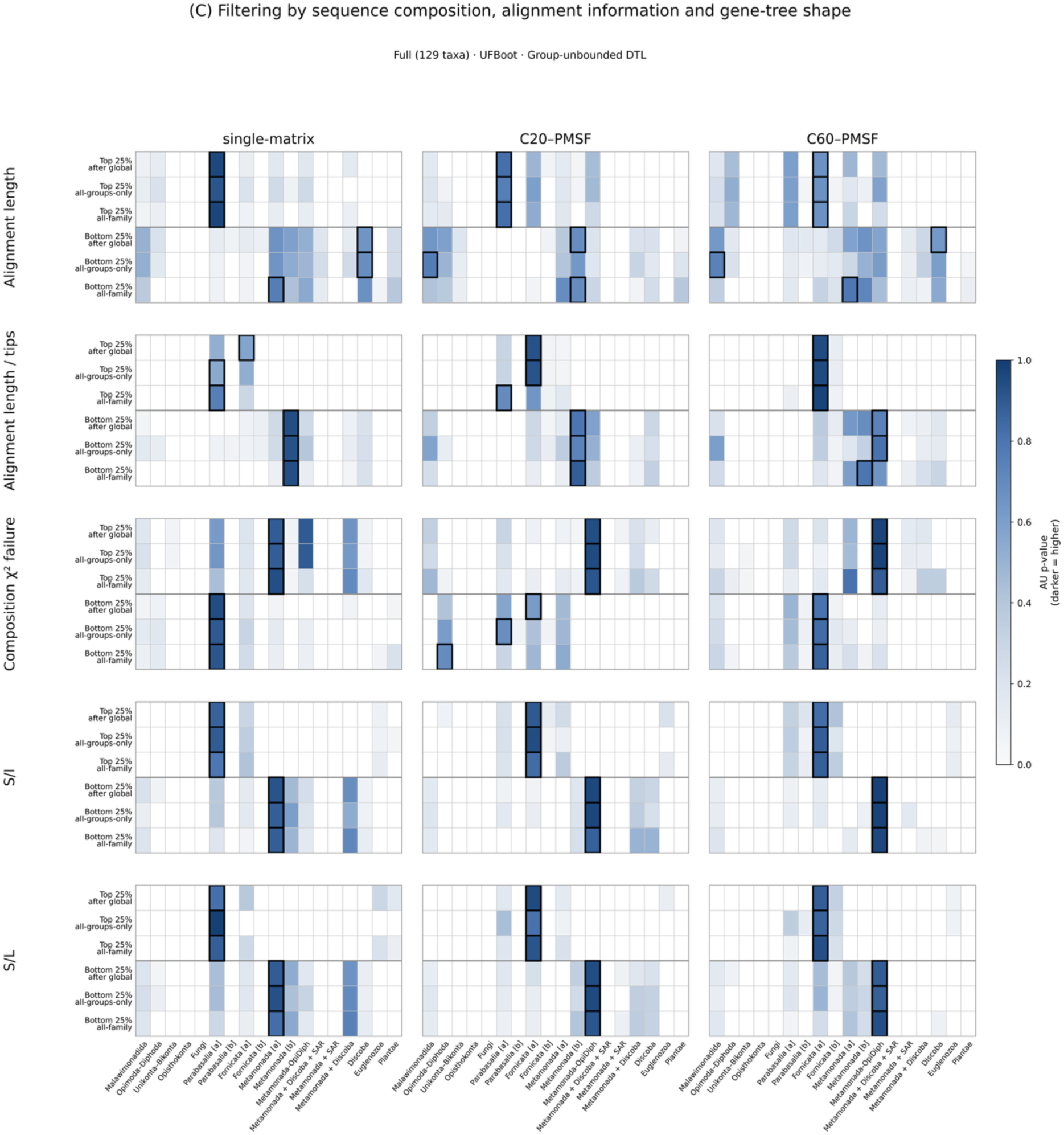
Tree-shape metrics and UFBoot-based sensitivity of eukaryotic root support to gene-family filtering under unbounded lineage-group-specific DTL. **(A)** Schematic illustration of S/I and S/L, where S is the longest internal backbone path, I the total internal branch length, and L the total tree length. Higher values indicate more backbone-concentrated gene trees, whereas lower values indicate bushier trees. Heatmaps show AU support for 18 candidate rooted species trees in the full 129-taxon dataset following filtering by **(B)** gene-family evolutionary properties (verticality, transfer count and copy number per represented genome) and **(C)** sequence- and tree-based metrics (alignment length, tip-normalised alignment length, composition χ² failure rate, S/I and S/L). Columns represent single-matrix, C20–PMSF and C60–PMSF UFBoot gene trees. Rows represent high- and low-quartile subsets under three strategies: all-family ranking, all-groups-only ranking and all-groups filtering after global ranking. Darker shading indicates higher AU support; white indicates AU < 0.05. Black outlines indicate the configuration with the highest summed reconciliation log-likelihood within each filtering subset and gene-tree framework.

In the full dataset, gene families with higher verticality or fewer inferred transfers generally favoured roots within Metamonada, especially within Fornicata and Parabasalia under lineage-specific DTL models. Similarly, families with more alignment columns per tip and higher S/I or S/L values more frequently favoured roots within Fornicata or Parabasalia, although alignment length alone showed a less consistent pattern. Lower-copy-number families also tended to favour roots within Fornicata or Parabasalia, whereas higher-copy-number families more frequently favoured roots on or near Metamonada. Filtering by composition-failure rate produced more model-dependent results (Figure 5B,C; Supplementary Table S11).

We also examined families represented in all major eukaryotic lineages (129 of 219 families in the full dataset and 138 of 219 in the reduced dataset). In the full dataset, the highest-likelihood roots varied among Fornicata, Parabasalia and Metamonada depending on the reconciliation model, whereas the reduced dataset frequently favoured roots on or near Ancyromonadida (Supplementary Table S11).

Overall, these analyses did not suggest that support for roots on or near Metamonada was driven by gene families with less reliable phylogenetic signal. We next explored whether unmodelled features of sequence or gene family evolution might drive the root signal to fast-evolving lineages using simulations.

### Might support for a root on, or near, Metamonads, derive from artifacts in the reconciliation analysis?

Our study was designed to exploit a source of root information that is distinct from concatenation-based outgroup rooting, so as to potentially ameliorate artifacts such as long branch attraction. It is therefore interesting that the credible set of roots supported by reconciliation analyses are reminiscent of recent concatenation-based studies, in finding support for a metamonad or Opimoda-Diphoda root. While agreement between concatenation and reconciliation is in principle encouraging, it seems possible that some of the same biases might affect both kinds of analysis, since - for example - reconciliation is informed by gene tree topologies which are themselves potentially subject to systematic error. While the analyses of empirical data did not tend to support the possibility that metamonad support is an LBA artifact (since it did not decrease with improved model fit, and appeared to be supported by more reliable gene families), we also sought to investigate how reconciliation-based root inference in this setting might be affected by gene tree error and other biases using simulations.

We first investigated the impact of high gene loss rates on reconciliation-based root inference. In our empirical analyses, we noticed that root support is sometimes affected by gene absence, potentially due to a “small genome artifact” described previously (Huang et al. 2025). For example, gene families that lacked representative sequences from a given clade generally had a tendency to prefer a root on that clade, perhaps because absences can be explained not only by gene loss, but also by scenarios in which the absent gene family evolved after that clade diverged from the others in the tree. This pattern was evident, for example, for Fungi, Ancyromonads, Unikonts and Bikonts, although not for Metamonads (Supplementary Note 1; Supplementary Figs. S8–S9). In previous work, this pattern has also added artifactual support to a “DPANN” root for Archaea (Huang et al. 2025). To investigate whether high loss rates might inflate support for metamonad-associated roots, we performed simulations on a fixed species tree rooted on Opimoda-Diphoda, calibrated so that families would have similar occupancy characteristics to the real data (see Methods), in which the metamonad loss rate was set to 1-, 2, 4…16 times the global loss rate; on the real data, metamonad gene loss rates are inferred to be ∼2-2.5 times the average. When inference was performed with the true model, the correct root was obtained with a strong likelihood lead over other hypotheses, as expected (median 1247 LL units). As loss rates in the generating model were increased, the lead for the Opimoda-Diphoda root decreased monotonically, suggesting that unmodelled loss can indeed affect root placement (Figure 6a; Supplementary Table S12). However, even with unmodelled loss rates much higher than the observed values, the simulated Opimoda-Diphoda root came out ahead (e.g., by 151 LL at 16x metamonad loss rates), suggesting that unmodelled loss is unlikely to explain the support for metamonad roots in the empirical data. Interestingly, somewhat lower loss levels (4-fold the average) were sufficient to recover a metamonad root if the true (simulated) root was closer to metamonads, in particular on Discoba (Figure 6b; Supplementary Table S12).

**Fig. 6.**
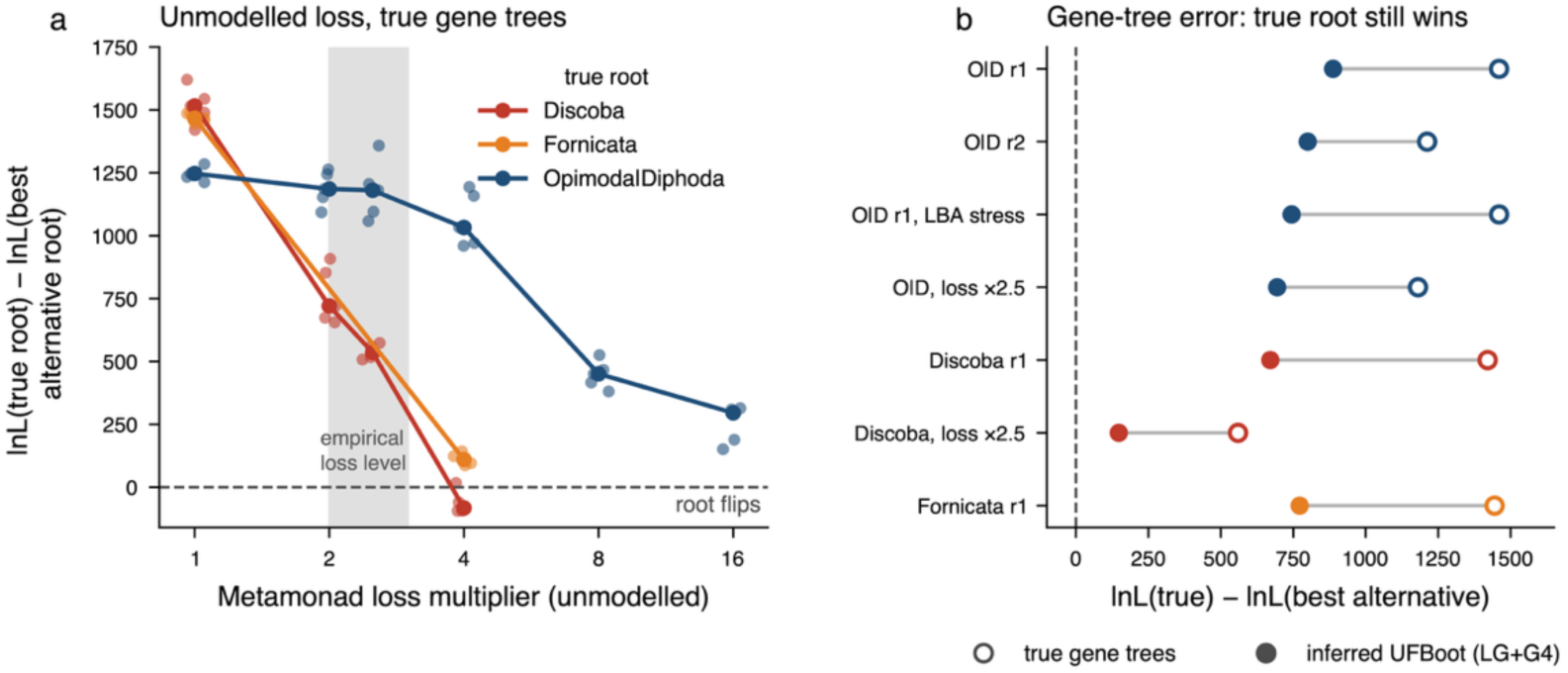
Effects of unmodelled gene loss and gene-tree estimation error on reconciliation-based inference of the eukaryotic root. **(a)** Effect of increasing unmodelled metamonad-specific gene loss on support for the true root under alternative simulated rooting scenarios (Opimoda–Diphoda, Discoba and Fornicata). The y-axis shows the difference in reconciliation log-likelihood between the true root and the best alternative root (ΔlogL), with negative values indicating that an incorrect root is favoured. Points show individual simulation replicates, and lines connect median values. The shaded region indicates the approximate range of metamonad gene-loss rates inferred from the empirical data. **(b)** Effect of gene-tree estimation error on reconciliation-based root inference under representative simulation conditions, including increased gene loss and substitution-model misspecification. Open circles indicate results obtained using the true simulated gene trees, whereas filled circles indicate results using inferred gene trees. Horizontal lines connect results from corresponding simulation conditions. Positive ΔlogL values indicate that the true root is preferred over the best alternative.

The above simulations were performed without gene tree error, but a reasonable hypothesis might be that LBA in individual gene trees draws fast-evolving metamonads to the base of paralogues. To investigate whether LBA resulting from substitution model misspecification might drive support for metamonad roots, we simulated paralogue gene trees under a site-heterogeneous model (LG+C60+G4) and analysed the data under LG+G4. As an additional stress test, we also performed simulations in which a composition shift matching the empirical difference in composition between metamonads and other taxa occurred along the metamonad stem, which might be predicted to increase support for metamonad roots when data were analyzed with the branch homogeneous LG+G4 model. As was the case with the gene loss simulations, deviations from the generating model of increasing magnitude did reduce the likelihood lead of the true rooted tree, but did not result in the (false) recovery of a metamonad root when the true root was on the Opimoda-Diphoda split (Figure 6b; Supplementary Table S12). Finally, we combined unmodelled loss and gene tree error in a single analysis (Figure 6b; Supplementary Table S12); the likelihood lead of the true (simulation) tree again decreased, but not to the point that a metamonad root obtained a higher likelihood.

Overall, these analyses suggest that reconciliation rooting can be impacted by site- and branch-heterogeneity, and by deviations from the reconciliation model. However, at least under the conditions tested, effects substantially greater than those that appear to obtain in the empirical data would be required to seriously mislead the analysis. Nevertheless, real data are more complex, and simulations provide only a partial exploration of the factors that might influence root support in the empirical analysis. A particularly relevant finding from the simulations is that the degree of model misspecification required to shift the maximum likelihood root from its true position in simulations depends on the distance between the true and alternative roots on the unrooted species tree, with much more modest departures sufficient to shift support from, for example, a Discoba root to Metamonada, or vice versa. This might help to explain why credible root sets in reconciliation analyses often include branches that are adjacent or otherwise close in the species tree, as is also the case in the present study (Figure 7).

**Figure 7.**
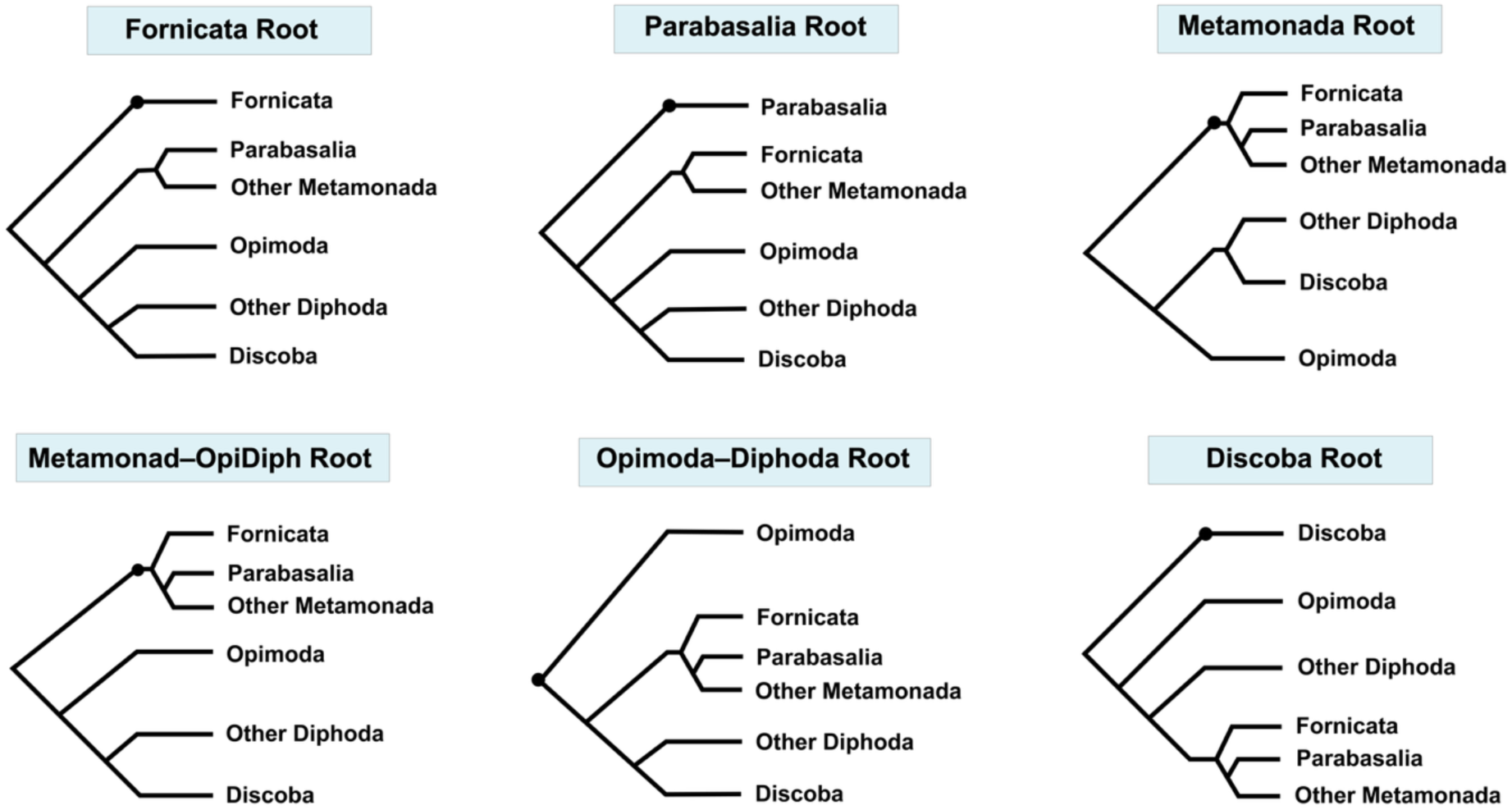
Schematic representation of alternative eukaryotic root hypotheses. Simplified rooted cartoon trees illustrating six rooting hypotheses not rejected in the reconciliation analyses: Fornicata, Parabasalia, Metamonada, Metamonad–OpiDiph, Opimoda–Diphoda and Discoba. Fornicata, Parabasalia and other metamonad lineages are shown separately to illustrate alternative root placements within or near Metamonada. The full-dataset analyses generally favoured roots on or within Metamonada, while alternative hypotheses remained unrejected by the AU test under some gene-tree and reconciliation-model combinations (Figure 4). The cartoons represent simplified branching relationships; some hypotheses were evaluated on different underlying species-tree backbones. Branch lengths are schematic and do not represent divergence times.

### Implications for LECA and eukaryogenesis

Previous attempts to resolve the root of eukaryotic phylogeny have relied on analysis of single-copy genes (Derelle and Lang 2012; He et al. 2014; Derelle et al. 2015; Al Jewari and Baldauf 2023). Our study provides a complementary test of these phylogenetic hypotheses by integrating phylogenomic evidence from 219 gene families that contain gene duplications inferred to predate the last eukaryotic common ancestor (LECA) across 131 representative eukaryotic taxa. The widespread distribution of these ancient duplicate families underscores the extent of genomic innovation that had already occurred before the diversification of extant eukaryotic lineages. In particular, the fixation of many gene duplicates associated with information processing, the cytoskeleton, membrane trafficking, metabolism and mitochondrial processes is consistent with previous reports that elaboration of nuclear information-processing systems, development of the cytoskeleton and endomembrane system, and integration of the mitochondrion into the nascent eukaryotic cell were key aspects of evolution along the eukaryotic stem lineage (Vosseberg et al. 2021; Kay et al. 2025; Pittis and Gabaldón 2016; Vosseberg et al. 2024; Martin et al. 2015; Roger et al. 2017). The information-processing duplicates include genes involved in transcriptional regulation, RNA processing, and DNA replication and repair, while other duplicated families are associated with vesicle trafficking, cytoskeletal organization, and mitochondrial metabolism.

Taken together, these results suggest that current duplication-based evidence is consistent with a root region involving Metamonada, Discoba, a deep split between metamonads and Opimoda-Diphoda, or a root on Opimoda-Diphoda, results that are compatible with a number of concatenation studies that each strongly supported a one or subset of these hypotheses (He et al. 2014; Derelle et al. 2015; Al Jewari and Baldauf 2023; Williamson et al. 2025). While metamonad roots have been argued to derive from long branch attraction in the concatenation context, our results do not appear to lend additional support to this possibility; instead, based on our metrics, it is the more reliable gene families that tend to support metamonad roots in the reconciliation framework, although simulations do identify the possibility that high rates of gene loss might artifactually bolster support for these lineages. This identified root region broadly overlaps the “root zone” encompassing most previously proposed root positions on the unrooted eukaryotic tree summarized by (Lamża, 2026). Related root positions have also been recovered using independent outgroup-based approaches. He et al. (2014), for example, recovered Discoba, and potentially its parent taxon Excavata, as sister to the remainder of the sampled eukaryotic diversity, while other phylogenomic analyses have also recovered rooting configurations involving or adjacent to Metamonada, Discoba, or other excavate-associated lineages (He et al. 2014; Williamson et al. 2025; Al Jewari and Baldauf 2023). As reconciliation-based rooting using pre-LECA gene duplicates is an entirely distinct source of rooting information compared to traditional outgroup-rooting approaches, the agreement between the results of our analyses and some recent outgroup-rooting studies is encouraging. Although the two approaches are subject to different sources of systematic error, their partial convergence on roots involving Metamonada and neighbouring lineages is consistent with the possibility that the root lies in this region. However, the sensitivity of the preferred configuration to reconciliation assumptions and gene-family properties cautions against assigning the root to a single branch. This restricted root region is summarized schematically in Figure 7.

In terms of character evolution, the inference of a complex, aerobic (or facultatively anaerobic), mitochondrion-possessing LECA is largely insensitive to the precise position of the root, because the core cellular features supporting this reconstruction are broadly conserved across major extant eukaryotic groups (Koumandou et al. 2013; Koonin 2010). Reductive genome and mitochondrial evolution, together with specialization for anaerobic and microaerophilic niches, would therefore have occurred during the subsequent evolution of Metamonada lineages. This does not necessarily imply that anaerobic metabolism originated de novo on the metamonad stem; the specialized metabolism of extant metamonads may instead reflect retention and modification of ancestral anaerobic metabolic capacities and subsequent reductive evolution (Müller et al., 2012). The credible rooting hypotheses retained across our analyses are also compatible with the suggestion of a LECA that possessed some excavate-like morphological traits (Williamson et al. 2025; (Lamża, 2026)), including a ventral feeding groove, vane-bearing posterior flagellum, and the associated cytoskeletal architecture. Lamża (2026) similarly reconstructed the ancestral eukaryote as an excavate-like biflagellate bacterivore, most likely adapted to microaerophilic conditions. However, the alternative rooting configurations tested here differ in the arrangement of the earliest-diverging lineages. In particular, the Opimoda–Diphoda configuration used in our tests differs in its internal topology from the Opimoda+–Diphoda+ configuration derived from Williamson et al. (2025), in which Metamonada and Discoba occur as early-diverging lineages on opposite sides of the root. We therefore tested the OpiDip+ configuration separately rather than treating these topological differences as consequences of root placement alone. These alternative rooted topologies therefore imply different initial splits during the eukaryotic radiation, although the rooting results alone do not uniquely determine the metabolic or ecological states of the resulting descendant lineages.

A root on Discoba was not rejected under some analysis conditions, particularly under bounded lineage-specific DTL with C20–PMSF and C60–PMSF gene trees, and our simulations admit the possibility that metamonad root support might result from attraction of a nearby root into the Metamonada due to high loss rates or gene tree error. A discoban root was previously proposed by He et al. (2014). Discoba is one of the three lineages (together with Malawimonadida and Metamonada) previously suggested to be part of an Excavata group united by their shared excavate characters (Simpson 2003). Under a Discoba root, LECA remains compatible with a complex, aerobic and mitochondrion-bearing reconstruction, as above. Discoba includes the jakobids, which possess some of the most gene-rich mitochondrial genomes known (e.g. 66 mitochondrially-encoded genes in *Andalucia godoyi*) (Gray et al. 2020). These gene-rich mitochondrial genomes retain unusually large complements of genes from the ancestral mitochondrial genome and therefore provide an informative window into early mitochondrial genome evolution. Regardless of the precise root position, mitochondrial acquisition and extensive integration of the endosymbiont into the host cell must have preceded LECA.

While our analyses discriminate among many previously proposed positions for the eukaryotic root, they do not robustly resolve several alternative configurations involving Metamonada and its constituent lineages, Discoba and Opimoda–Diphoda. While the number of pre-LECA duplicates we identified and analyzed is comparable to, and in some cases greater than, the number of genes used in concatenation-based, outgroup-rooting approaches, a natural next step to boost power might be to bring more data to bear by analysing more gene families for a representative sample of eukaryotic genomes, and by obtaining more genomic and transcriptomic data from undersampled metamonad lineages. However, our analyses also highlight an important challenge for such efforts. Even among the widely conserved pre-LECA duplicate families analysed here, root support is sensitive to gene-family properties, including inferred transfer count and verticality. . Expanding the dataset to include less conserved families will increase the amount of potentially informative signal, but may also exacerbate the difficulty of distinguishing genuine deep phylogenetic history from patterns generated by differential loss and incomplete representation across lineages. Further progress will likely depend not only on broader family sampling, but also on improved reconciliation models that more adequately capture lineage-specific patterns of gene loss, absence, and retention. The partial convergence between duplication-based reconciliation and traditional outgroup-rooting approaches is encouraging, suggesting that these complementary sources of evidence may be narrowing the search for the eukaryotic root.

## Materials & Methods

### Taxon sampling

To maximize phylogenetic coverage and ensure balanced representation of the major eukaryotic lineages, we assembled a dataset of 131 representative species spanning the major diversity of extant eukaryotes, including Amorphea (Opisthokonta, Amoebozoa), Discoba (Jakobida, Euglenozoa, Heterolobosea), Archaeplastida (Glaucophyta, Rhodophyta, Viridiplantae), SAR (Stramenopila, Alveolata, Rhizaria), Cryptista, Haptista and Metamonada. Taxon sampling followed the most recent large-scale phylogenomic frameworks (Kay et al., 2025; Torruella et al., 2025; Williamson et al., 2025), ensuring both completeness and comparability across clades. The complete list of sampled taxa, dataset identifiers and lineage assignments, together with their data sources, is provided in Supplementary Table S2.

The proteomic data were obtained from public repositories including NCBI (O’Leary et al., 2016) and the EukProt v3 database (Richter et al., 2020). For each lineage, we selected the highest-quality and most up-to-date datasets available, excluding assemblies that were redundant or of low quality. The final dataset of 131 eukaryotic taxa was used for the initial gene-family analyses and gene-tree inference. The principal AleRax reconciliation and rooting analyses were performed using a 129-taxon dataset, excluding two hemimastigophoran taxa (*Hemimastix kukwesjijk* and *Spironema* sp.). An additional 117-taxon dataset was generated by further excluding nine metamonad taxa, two malawimonad taxa and *Pygsuia biforma* for taxon-sampling sensitivity analyses.

### Identification of pre-LECA duplicate families

Our analysis began with the 19,630 cross-domain gene family trees generated by (Kay et al., 2025) using the Domain Origins (DO) pipeline (Tria et al., 2021), which integrates bacterial, archaeal and eukaryotic homologues into unified gene-family phylogenies. These trees served as the initial dataset from which candidate pre-LECA duplicate families were identified.

To detect ancient duplication events, we systematically screened each unrooted gene tree for topological splits separating two eukaryotic paralogue clades, each containing representatives from multiple major eukaryotic supergroups. Such patterns indicate that the corresponding paralogues must already have existed before the diversification of extant eukaryotes, and are therefore consistent with duplications predating the last eukaryotic common ancestor (LECA) (Tria et al., 2021; Vosseberg & Snel, 2017). We therefore use “pre-LECA duplicate family” here as an operational term for families in which both paralogue clades were already present by LECA, rather than as an assignment of the duplication event to a specific rooted branch.

Applying these criteria yielded 219 candidate pre-LECA duplicate families (Supplementary Table S1). These curated gene families formed the basis for all subsequent phylogenomic and reconciliation analyses.

### Gene tree inference

For each gene family, protein sequences were first aligned with MAFFT v7.480 (Katoh & Standley, 2013), and poorly aligned regions were removed using BMGE (Criscuolo & Gribaldo, 2010). Maximum-likelihood gene trees were then inferred with IQ-TREE 2 (Minh et al., 2020) under multiple substitution-model frameworks. For the primary analyses, we inferred single-matrix gene trees using ModelFinder Plus (-m MFP) to identify the best-fitting site-homogeneous empirical substitution model for each family. To assess the sensitivity of downstream reconciliation analyses to substitution-model choice, we also analysed gene trees under a C20–PMSF site-heterogeneous profile-mixture framework, implemented by extending the MFP-selected best-fitting single-matrix model with C20+F+G and re-optimizing the corresponding single-matrix tree topologies under this framework, and under the LG+C60 model implemented with the PMSF approximation. Branch support was assessed using ultrafast bootstrap (UFBoot), with 1,000 bootstrap replicate trees generated per gene family. These UFBoot replicate trees were used as the input for all subsequent AleRax reconciliation, rooting and sensitivity analyses, thereby accounting for gene-tree uncertainty.These procedures yielded three UFBoot gene-tree sets for the 219 retained pre-LECA duplicate families, which were used in subsequent reconciliation and sensitivity analyses. All inferred gene trees used in this study, including the single-matrix, C20–PMSF and C60–PMSF tree sets, are provided in Supplementary Data 2, with separate files or directories for each substitution-model framework.

### Species tree inference

To infer the species-tree topology of eukaryotes, we used AleRax v1.4.0 (Morel et al., 2024) on the 219 high-confidence pre-LECA duplicate gene trees inferred under the single-matrix framework. AleRax estimates the species tree by jointly evaluating concordance and conflict among unrooted gene trees under a duplication–transfer–loss (DTL) framework. Species-tree searches were performed using the HYBRID mode (--species-tree-search HYBRID), starting from a random initial tree (-s random); MiniNJ-derived starting trees (-s MiniNJ) produced identical topologies. We also repeated the species-tree search using the corresponding C20– PMSF and C60–PMSF maximum-likelihood gene trees. For the final topology used in downstream analyses, Metamonada and Archaeplastida were treated as monophyletic, and the positions of the apusomonads *Thecamonas* and *Manchomonas* were modified to group them with the breviate *Pygsuia*, consistent with previous phylogenomic evidence. Because this manual modification altered the AleRax-inferred topology, the original AleRax branch-support values were not transferred to the modified tree. Support for the final topology was instead evaluated using ASTRAL-Pro3 quartet support (q1) against the same set of 219 gene trees.

Although AleRax returns a rooted species tree under its branch-homogeneous DTL model, this rooting was not used in our analyses. Because our objective was to evaluate alternative rooting hypotheses without conditioning the analyses on the root inferred during species-tree estimation, we used the AleRax output solely as an unrooted species-tree topology, which served as the baseline framework for downstream reconciliation-based root inference under both branch-homogeneous and branch-heterogeneous DTL models. Additional species-tree configurations were also evaluated for rooting hypotheses involving alternative relationships among major eukaryotic lineages. The final modified species-tree topology, with ASTRAL-Pro3 quartet support (q1), is shown in Figure 3, whereas the original AleRax-inferred topology with its local posterior probability (LPP) support values is shown in Supplementary Fig. S2. The corresponding tree files, including the AleRax-inferred species trees based on the single-matrix, C20–PMSF and C60–PMSF maximum-likelihood gene-tree sets, and the final modified tree used for downstream reconciliation analyses, are available in Supplementary Data 1.

### Gene tree-species tree reconciliation and root hypothesis testing

To infer the root of the eukaryotic species tree, we performed gene tree–species tree reconciliation using AleRax v1.4.0 (Morel et al., 2024) on the 219 pre-LECA duplicate gene families. All reconciliation-based rooting analyses were performed using UFBoot gene trees inferred under the single-matrix, C20–PMSF and C60–PMSF substitution-model frameworks. This framework explains the incongruence between gene trees and the species tree in terms of gene duplication (D), transfer (T), and loss (L) events, and integrates over all possible DTL event histories to evaluate the likelihood of different rooting hypotheses. Because different reconciliation models may favor different rooting hypotheses, we explored a range of DTL models—a global model, in which all species-tree branches share a common set of duplication, transfer and loss rates, and two lineage-specific models, in which predefined eukaryotic lineage groups have separate D, T and L rates. The lineage-specific analyses included an unbounded model and a bounded model (box2), in which the group-specific rates were constrained by an upper bound of 2. Within this framework, we evaluated statistical support for 15 alternative eukaryotic rooting hypotheses, including the Fungi root (Cerón-Romero et al., 2022), the Opimoda–Diphoda hypothesis (Derelle et al., 2015; Derelle & Lang, 2012; Williamson et al., 2025), the Plantae root (Rogozin et al., 2009), the Discoba root (He et al., 2014), the Euglenozoa root (Cavalier-Smith, 2010), the Opisthokonta root (Cerón-Romero et al., 2022; Stechmann & Cavalier-Smith, 2002), the Metamonada root (Torruella et al., 2025) , and the Unikonta–Bikonta hypothesis (Stechmann & Cavalier-Smith, 2002; Derelle & Lang, 2012), as well as the Parabasalia root(Al Jewari & Baldauf, 2023) and the Opimoda+/Diphoda+ (OpiDip+) hypothesis(Williamson et al., 2025). We also tested the ‘Metamonada-OpiDiph’ hypothesis, in which Metamonada lies outside the Opimoda–Diphoda division. This hypothesis has not been explicitly proposed in the literature but was included as an additional configuration motivated by uncertainty in the placement of Metamonada relative to the Opimoda–Diphoda division. We further tested three alternative rooting hypotheses involving Metamonada + Discoba, Metamonada + SAR, and Metamonada + Discoba + SAR. These additional configurations were included to explore alternative placements of Metamonada relative to other major eukaryotic lineages, in light of recent phylogenomic analyses(Torruella et al., 2025; Williamson et al., 2025). In addition, we tested a scheme to place the root in Malawimonadida. This root position has been explicitly proposed in the literature(Cavalier-Smith, 2022), and malawimonads have also been recovered as a deeply branching group in several phylogenetic studies(He et al., 2014; Torruella et al., 2025). For the analyses, these rooting hypotheses were represented by 18 rooted species-tree configurations in the principal 129-taxon dataset, including alternative placements evaluated on different species-tree backbone topologies. We also analysed a reduced 117-taxon dataset, for which 12 rooted configurations were evaluated. For each candidate rooting configuration, we calculated the reconciliation log-likelihood for each gene family under each gene-tree framework and DTL model. We then compared the summed reconciliation log-likelihoods across configurations using an AU test (Approximately Unbiased test; Shimodaira, 2002) implemented in the CONSEL package (Shimodaira & Hasegawa, 2001). The AU test was applied to the per-family reconciliation log-likelihoods, using gene families as the resampling units, to assess whether alternative rooting configurations could be statistically rejected; configurations with AU ≥ 0.05 were considered unrejected.

### Sensitivity analyses

To evaluate the robustness of alternative rooting hypotheses and to identify potential sources of systematic bias, we performed sensitivity analyses targeting substitution models, reconciliation model assumptions, and gene-family composition. All sensitivity analyses were performed using UFBoot gene trees and the corresponding AleRax reconciliation likelihoods.

Substitution model comparisons Gene trees were analysed under three substitution-model frameworks: (i) the best-fitting site-homogeneous models selected by ModelFinder Plus in IQ-TREE 2 (Kalyaanamoorthy et al., 2017), (ii) a C20 site-heterogeneous profile-mixture framework (Lartillot & Philippe, 2004), implemented by extending the MFP-selected best-fitting single-matrix model with C20+F+G and re-optimizing the corresponding single-matrix tree topologies under this framework, and (iii) the LG+C60 empirical profile-mixture model implemented using the PMSF two-step procedure (Quang et al., 2008; Wang et al., 2018). These alternative UFBoot gene-tree sets were used in downstream reconciliation analyses to assess whether substitutional heterogeneity influenced likelihood support for competing rooting hypotheses.

Reconciliation model

To test whether root inferences were sensitive to assumptions about gene-family loss dynamics, we recalculated species-tree likelihoods for each candidate rooting hypothesis under three duplication–transfer–loss (DTL) reconciliation frameworks applied to the same gene trees: (i) a global model with a single shared set of D, T and L rates across the species tree; (ii) an unbounded lineage-specific model; and (iii) a bounded lineage-specific model (box2). In the lineage-specific models, species were assigned to 12 predefined eukaryotic lineage groups—Plantae, SAR, Discoba, Metamonada, Holomycota, Holozoa, Ancyromonadida, Malawimonadida, CRuMs, Amoebozoa, Apusomonadida and Pygsuia biforma—with a separate set of D, T and L parameters estimated for each group. The bounded model additionally constrained group-specific rate estimates using a bound of 2.This allowed us to assess the effect of lineage-specific DTL-rate modelling and rate constraints on support for alternative rooting hypotheses.

Gene-family filtering and subsetting To assess whether support for alternative rooting hypotheses was influenced by gene-family properties, we constructed multiple filtered datasets based on species representation, inferred transfer count, copy number per represented genome, composition-failure rate, alignment length, tip-normalized alignment length, verticality, and tree shape. Verticality was quantified as V/(V+D+T+L), where V denotes inferred events of vertical descent and D, T and L denote inferred duplication, transfer and loss events, respectively. Because verticality depends on the assumed species-tree root, it was calculated separately under each of the 18 rooted configurations in the 129-taxon dataset (or 12 configurations in the reduced 117-taxon dataset), and the resulting root-specific values were averaged for each gene family within each gene-tree framework. Tree shape was quantified using S/I and S/L backbone concentration metrics, where S is the length of the longest internal backbone path, I is the total internal branch length, and L is the total tree length. These metrics were calculated after pruning the UFBoot gene trees to the corresponding taxon dataset, with values averaged across 1,000 bootstrap replicates per gene family.

Inferred transfer count was calculated from AleRax reconciliation outputs by summing inferred transfer events across species-tree branches for each gene family and rooting hypothesis. For UFBoot analyses, transfer counts were averaged across sampled reconciliations before being averaged across the 15 tested rooting hypotheses. Copy number per represented genome was calculated as the number of terminal tips in the gene tree divided by the number of genomes represented in that family. This metric was used as a tip-based proxy for relative gene-family copy-number expansion rather than as a reconciliation-inferred duplication count. Composition-failure rate was calculated as the proportion of sequences in each gene family failing the amino-acid composition χ² test implemented in IQ-TREE. Composition-failure rates were calculated separately for the 129- and 117-taxon datasets using the corresponding taxon-pruned alignments.

Full lineage coverage was defined as the presence of at least one representative from each major sampled eukaryotic lineage. This criterion was satisfied by 129 of the 219 families in the 129-taxon dataset and 138 families in the reduced 117-taxon dataset. To evaluate the effect of broad taxonomic coverage, we used three complementary filtering strategies. First, in the all-family ranking strategy, all retained gene families were ranked by each metric, and top- and bottom-quartile subsets were selected. Second, in the all-groups-only ranking strategy, we first restricted the dataset to families satisfying the full-lineage-coverage criterion (“all-groups” datasets), and then ranked these families separately by each metric before defining top- and bottom-quartile subsets. Third, in the all-groups filtering after global ranking strategy, we first ranked all retained families by each metric and selected the top and bottom quartiles, and then applied the full-lineage-coverage filter. Because this third procedure removed different numbers of families from the upper and lower quartiles, the resulting subsets were not always the same size. High and low quartiles were defined separately for the corresponding taxon datasets and ranking strategies. The resulting metric values, ranks, and quartile assignments for all gene families are provided in Supplementary Table S10.

To examine the contribution of metamonad representation more directly, we also partitioned gene families according to whether they included metamonad homologues (“metamonad-present”) or lacked them (“metamonad-absent”). This allowed us to test whether root support was disproportionately influenced by the inclusion or exclusion of metamonad-bearing families.

Together, these procedures yielded subsets defined by full lineage coverage, metamonad presence or absence, high or low inferred transfer count, high or low copy number per represented genome, high or low composition-failure rate, high or low S/I and S/L backbone concentration, high or low verticality, high or low alignment length, and high or low tip-normalised alignment length. Reconciliation analyses and likelihood-based statistical tests (AU, KH and SH) were repeated across these subsets to evaluate the robustness of competing rooting hypotheses under alternative data partitions. Gene-family ranking metrics used for filtering analyses are provided in Supplementary Table S10, and the corresponding AU-test results for the eight metric-based filtering analyses are presented in Supplementary Table S11 (Figure 5).

### Simulations

We asked whether the metamonad-type roots favoured by the empirical analyses could arise as an artefact when the true root lies elsewhere, focusing on a balanced root between Opimoda and Diphoda. We also explored simulations with a true root closer to metamonads (on the Discoba stem) and a root on the Fornicata stem (inside metamonads), to test recovery when the true root is at or near metamonads. All simulations used the 129-taxon backbone (Hemimastigophora excluded), rooted at each of the three positions.

Gene families were simulated with a custom forward simulator that reproduces the undated DTL model implemented in AleRax (Morel et al.): on every species branch, including the root branch and terminal branches, each gene copy undergoes speciation, duplication, transfer or loss with probabilities proportional to 1:D:T:L. Transfer recipients are drawn uniformly from branches that are neither the donor nor its direct ancestors (AleRax default), and transfers are therefore not constrained in time. Rates were set to the global-model estimates from the empirical data (D = 0.71, L = 0.89, T = 0.095 per branch), with gamma-distributed family-level rate multipliers. To mimic the selection of pre-LECA duplicates, families were retained only if they carried a duplication on the root branch with both paralogue subtrees present in ≥45 species, and at least three genes. With these settings simulated families matched the empirical families in size, species occupancy and copy-number distribution. Each dataset comprised 219 families, as in the empirical analysis. The simulator reproduced AleRax’s extinction probabilities exactly, and AleRax recovered the generating rates and the true root on unperturbed data. The custom UndatedDTL simulator and associated simulation and analysis scripts are provided in Supplementary Data 3.

To test unmodelled loss, the loss rate was multiplied by 2, 2.5, 4, 8 or 16 on every metamonad branch (including metamonad stem branches), while inference assumed a single global loss rate. The empirical metamonad loss multiplier, estimated by matching the occupancy of Fornicata, Parabasalia and Preaxostyla in the 219 families, was about 2–3. Further conditions were (i) loss ×4 on metamonads and on the other lineages that support unusual roots (Malawimonadida, Ancyromonadida, Apusomonadida, Pygsuia); (ii) strong among-family rate heterogeneity (gamma multipliers, shared across D, L and T, plus a variant with independent multipliers per rate (2 replicates)); and (iii) lineage-specific missing genes matched to empirical occupancy, fitted with and without AleRax’s fraction-missing correction. Five replicate datasets were simulated per condition.

For selected datasets we simulated protein alignments on the true gene trees and inferred gene trees from them. Gene-tree branch lengths were derived from species-branch lengths estimated from the 219 empirical single-matrix ML gene trees (metamonad branches are a median 2.1× longer than comparable branches; Fornicata 2.1–3.7×), with lognormal family rates and per-branch noise. Alignments were simulated with AliSim (IQ-TREE 3.1.4) under LG+C60+G4 at empirical alignment lengths (gap-free), and gene trees were inferred with IQ-TREE under LG+G4 with 1,000 ultrafast bootstrap replicates. This deliberate under-parameterisation (site-homogeneous inference on site-heterogeneous data) favours long-branch attraction. We also ran a combined condition, metamonad loss ×2.5 plus gene-tree error, with one dataset for each of the Opimoda-Diphoda and Discoba-rooted trees. In an additional long-branch-attraction stress test, metamonad branches were given the empirical metamonad amino-acid composition (each C60 profile tilted multiplicatively towards the metamonad/non-metamonad frequency ratio) and heterotachy (half the sites with metamonad branches ×4, half ×0.25).

Each dataset was analysed with AleRax under the global (one D, T, L) undated DTL model, with uniform origination as in the empirical analysis. For every candidate root the species tree was fixed (--species-tree-search SKIP) and rates re-optimised. A root search (REROOT) from the true root was run as a check for better rootings outside the panel. We used AleRax’s default four fixed-point iterations; twenty iterations changed root log-likelihood differences by <0.3% (although they recover D and L more accurately).

For simulated and empirical gene trees we computed the fraction of metamonad genes that branch first within their pre-LECA paralogue (same paralogue-split rooting and criterion as for the empirical trees), and its per-family correlation with support for metamonad roots.

## Data availability

The sequence alignments, gene-tree samples, species-tree configurations, and custom analysis and simulation code supporting this study are available in the Figshare repository at https://doi.org/10.6084/m9.figshare.35076463.

## Acknowledgments

We thank every researcher and reviewer who participated in this work. This work was supported by the China Scholarship Council, the Gordon and Betty Moore Foundation (grant GBMF9741 to TAW, AS, GJSz, DP and PCJD), and the John Templeton Foundation (grant 63451 to TAW, AS, GJSz, DP and PCJD); the opinions expressed in this publication are those of the authors and do not necessarily reflect the views of the John Templeton Foundation. We used Claude (Opus 5.5) to implement the UndatedDTL simulator.

## Author Contributions

CL, CK, TAW and OS performed analyses. TAW, PCJD, AS, DP and GJSz conceived analyses. All authors interpreted results and contributed to drafting and revision of the manuscript.

## References

Al Jewari, C., & Baldauf, S. L. (2023). An excavate root for the eukaryote tree of life. Science Advances, 9(17), eade4973.

Brown, J. R., & Doolittle, W. F. (1995). Root of the universal tree of life based on ancient aminoacyl-tRNA synthetase gene duplications. Proceedings of the National Academy of Sciences of the United States of America, 92(7), 2441–2445.

Cavalier-Smith, T. (2010). Kingdoms Protozoa and Chromista and the eozoan root of the eukaryotic tree. Biology Letters, 6(3), 342–345.

Cavalier-Smith, T. (2022). Ciliary transition zone evolution and the root of the eukaryote tree: implications for opisthokont origin and classification of kingdoms Protozoa, Plantae, and Fungi. Protoplasma, 259(3), 487–593.

Cerón-Romero, M. A., Fonseca, M. M., de Oliveira Martins, L., Posada, D., & Katz, L. A. (2022). Phylogenomic Analyses of 2,786 Genes in 158 Lineages Support a Root of the Eukaryotic Tree of Life between Opisthokonts and All Other Lineages. Genome Biology and Evolution, 14(8). 10.1093/gbe/evac119

Derelle, R., & Lang, B. F. (2012). Rooting the eukaryotic tree with mitochondrial and bacterial proteins. Molecular Biology and Evolution, 29(4), 1277–1289.

Derelle, R., Torruella, G., Klimeš, V., Brinkmann, H., Kim, E., Vlček, Č., Lang, B. F., & Eliáš, M. (2015). Bacterial proteins pinpoint a single eukaryotic root. Proceedings of the National Academy of Sciences of the United States of America, 112(7), E693–E699.

Gogarten, J. P., Kibak, H., Dittrich, P., Taiz, L., Bowman, E. J., Bowman, B. J., Manolson, M. F., Poole, R. J., Date, T., Oshima, T., Konishi, J., Denda, K., & Yoshida, M. (1989). Evolution of the vacuolar H+-ATPase: implications for the origin of eukaryotes. Proceedings of the National Academy of Sciences of the United States of America, 86(17), 6661–6665.

He, D., Fiz-Palacios, O., Fu, C.-J., Fehling, J., Tsai, C.-C., & Baldauf, S. L. (2014). An alternative root for the eukaryote tree of life. Current Biology, 24(4), 465–470.

Iwabe, N., Kuma, K., Hasegawa, M., Osawa, S., & Miyata, T. (1989). Evolutionary relationship of archaebacteria, eubacteria, and eukaryotes inferred from phylogenetic trees of duplicated genes. Proceedings of the National Academy of Sciences of the United States of America, 86(23), 9355–9359.

Katz, L. A. (2012). Origin and diversification of eukaryotes. Annual Review of Microbiology, 66(1), 411–427.

Kay, C. J., Spang, A., Szöllősi, G. J., Pisani, D., Williams, T. A., & Donoghue, P. C. J. (2025). Dated gene duplications elucidate the evolutionary assembly of eukaryotes. Nature. 10.1038/s41586-025-09808-z

Lamża, Ł. (2026). Deep-branching eukaryotes and early events in protist evolution. Biological Reviews of the Cambridge Philosophical Society, 101(2), 735–750.

Minh, B. Q., Schmidt, H. A., Chernomor, O., Schrempf, D., Woodhams, M. D., von Haeseler, A., & Lanfear, R. (2020). IQ-TREE 2: New Models and Efficient Methods for Phylogenetic Inference in the Genomic Era. Molecular Biology and Evolution, 37(5), 1530–1534.

Mongiardino Koch, N. (2021). Phylogenomic Subsampling and the Search for Phylogenetically Reliable Loci. Molecular Biology and Evolution, 38(9), 4025–4038.

Moody, E. R. R., Álvarez-Carretero, S., Mahendrarajah, T. A., Clark, J. W., Betts, H. C., Dombrowski, N., Szánthó, L. L., Boyle, R. A., Daines, S., Chen, X., Lane, N., Yang, Z., Shields, G. A., Szöllősi, G. J., Spang, A., Pisani, D., Williams, T. A., Lenton, T. M., & Donoghue, P. C. J. (2024). The nature of the last universal common ancestor and its impact on the early Earth system. Nature Ecology & Evolution, 8(9), 1654–1666.

Morel, B., Williams, T. A., Stamatakis, A., & Szöllősi, G. J. (2024). AleRax: a tool for gene and species tree co-estimation and reconciliation under a probabilistic model of gene duplication, transfer, and loss. Bioinformatics (Oxford, England), 40(4), btae162.

Müller, M., Mentel, M., van Hellemond, J. J., Henze, K., Woehle, C., Gould, S. B., Yu, R.-Y., van der Giezen, M., Tielens, A. G. M., & Martin, W. F. (2012). Biochemistry and evolution of anaerobic energy metabolism in eukaryotes. Microbiology and Molecular Biology Reviews : MMBR, 76(2), 444–495.

O’Leary, N. A., Wright, M. W., Brister, J. R., Ciufo, S., Haddad, D., McVeigh, R., Rajput, B., Robbertse, B., Smith-White, B., Ako-Adjei, D., Astashyn, A., Badretdin, A., Bao, Y., Blinkova, O., Brover, V., Chetvernin, V., Choi, J., Cox, E., Ermolaeva, O., … Pruitt, K. D. (2016). Reference sequence (RefSeq) database at NCBI: current status, taxonomic expansion, and functional annotation. Nucleic Acids Research, 44(D1), D733–D745.

Philippe, H., Brinkmann, H., Lavrov, D. V., Littlewood, D. T. J., Manuel, M., Wörheide, G., & Baurain, D. (2011). Resolving difficult phylogenetic questions: why more sequences are not enough. PLoS Biology, 9(3), e1000602.

Philippe, H., & Roure, B. (2011). Difficult phylogenetic questions: more data, maybe; better methods, certainly. BMC Biology, 9(1), 91.

Philippe, H., Zhou, Y., Brinkmann, H., Rodrigue, N., & Delsuc, F. (2005). Heterotachy and long-branch attraction in phylogenetics. BMC Evolutionary Biology, 5(1), 50.

Richter, D. J., Berney, C., Strassert, J. F. H., Poh, Y.-P., Herman, E. K., Muñoz-Gómez, S. A., Wideman, J. G., Burki, F., & de Vargas, C. (2020). EukProt: A database of genome-scale predicted proteins across the diversity of eukaryotes. In bioRxiv (p. 2020.06.30.180687). bioRxiv. 10.1101/2020.06.30.180687

Rodríguez-Ezpeleta, N., Brinkmann, H., Roure, B., Lartillot, N., Lang, B. F., & Philippe, H. (2007). Detecting and overcoming systematic errors in genome-scale phylogenies. Systematic Biology, 56(3), 389–399.

Rogozin, I. B., Basu, M. K., Csürös, M., & Koonin, E. V. (2009). Analysis of rare genomic changes does not support the unikont-bikont phylogeny and suggests cyanobacterial symbiosis as the point of primary radiation of eukaryotes. Genome Biology and Evolution, 1, 99–113.

Rokas, A., & Carroll, S. B. (2005). More genes or more taxa? The relative contribution of gene number and taxon number to phylogenetic accuracy. Molecular Biology and Evolution, 22(5), 1337–1344.

Shen, X.-X., Hittinger, C. T., & Rokas, A. (2017). Contentious relationships in phylogenomic studies can be driven by a handful of genes. Nature Ecology & Evolution, 1(5), 126.

Shimodaira, H., & Hasegawa, M. (2001). CONSEL: for assessing the confidence of phylogenetic tree selection. *Bioinformatics (Oxford*, England), 17(12), 1246–1247.

Stechmann, A., & Cavalier-Smith, T. (2002). Rooting the eukaryote tree by using a derived gene fusion. *Science (New York*, N.Y.), 297(5578), 89–91.

Torruella, G., Galindo, L. J., Moreira, D., & López-García, P. (2025). Phylogenomics of neglected flagellated protists supports a revised eukaryotic tree of life. Current Biology : CB, 35(1), 198–207.e4.

Tria, F. D. K., Brueckner, J., Skejo, J., Xavier, J. C., Kapust, N., Knopp, M., Wimmer, J. L. E., Nagies, F. S. P., Zimorski, V., Gould, S. B., Garg, S. G., & Martin, W. F. (2021). Gene duplications trace mitochondria to the onset of eukaryote complexity. Genome Biology and Evolution, 13(5). 10.1093/gbe/evab055

Vosseberg, J., & Snel, B. (2017). Domestication of self-splicing introns during eukaryogenesis: the rise of the complex spliceosomal machinery. Biology Direct, 12(1), 30.

Williamson, K., Eme, L., Baños, H., McCarthy, C. G. P., Susko, E., Kamikawa, R., Orr, R. J. S., Muñoz-Gómez, S. A., Minh, B. Q., Simpson, A. G. B., & Roger, A. J. (2025). A robustly rooted tree of eukaryotes reveals their excavate ancestry. Nature, 640(8060), 974–981.

Williams, T. A., Cox, C. J., Foster, P. G., Szöllősi, G. J., & Embley, T. M. (2020). Phylogenomics provides robust support for a two-domains tree of life. Nature Ecology & Evolution, 4(1), 138–147.

Williams, T. A., Schrempf, D., Szöllősi, G. J., Cox, C. J., Foster, P. G., & Embley, T. M. (2021). Inferring the deep past from molecular data. Genome Biology and Evolution. 10.1093/gbe/evab067

Zhaxybayeva, O., & Gogarten, J. P. (2004). Cladogenesis, coalescence and the evolution of the three domains of life. Trends in Genetics : TIG, 20(4), 182–187.

