## Supplementary Figures 1-9 for "Rooting the tree of eukaryotes using ancient gene duplications"

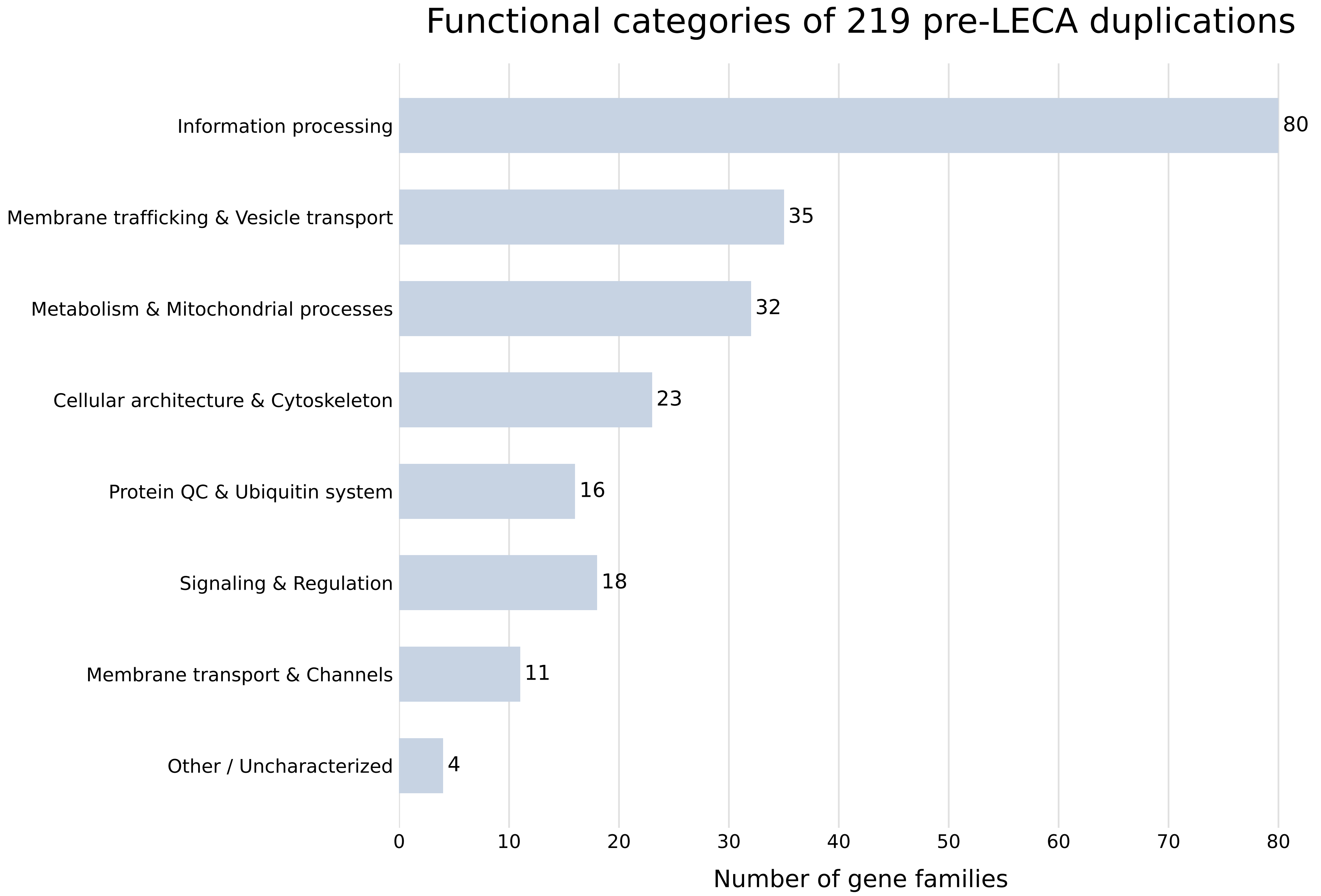
Functional categories of 219 pre-LECA duplications
Information processing
Membrane trafficking & Vesicle transport
Metabolism & Mitochondrial processes
Cellular architecture & Cytoskeleton
Protein QC & Ubiquitin system
Signaling & Regulation
Membrane transport & Channels
Other / Uncharacterized
80
35
32
23
16
18
11
4
0
10
20
30
40
50
60
70
80
Number of gene families

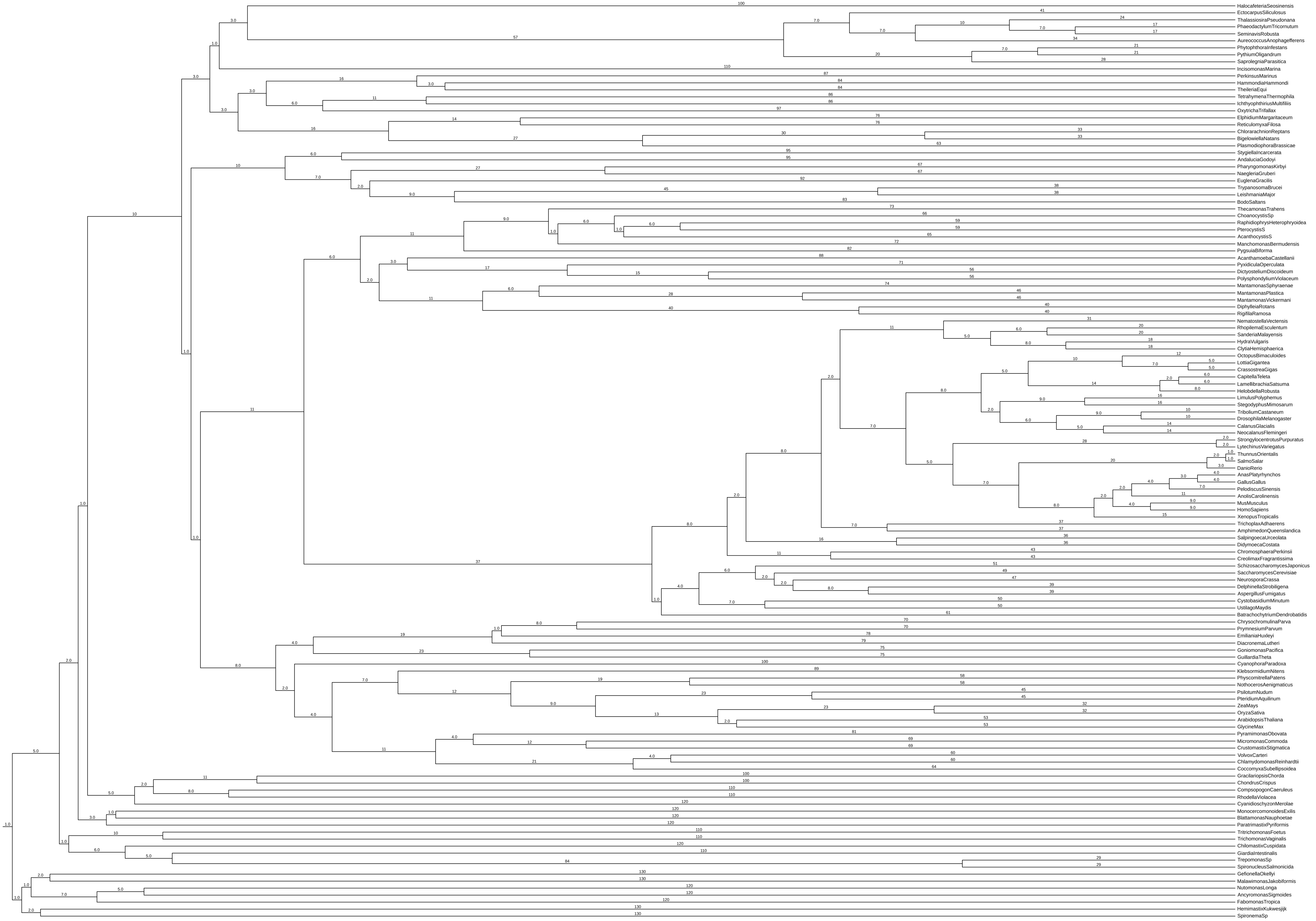

HalocafeteriaSeosinensis
EctocarpusSiliculosus
ThalassiosiraPseudonana
PhaeodactylumTricornutum
SeminavisRobusta
AureococcusAnophagefferens
PhytophthoraInfestans
PythiumOligandrum
SaprolegniaParasitica
IncisomonasMarina
PerkinsusMarinus
HammondiaHammondi
TheileriaEqui
TetrahymenaThermophila
IchthyophthiriusMultifiliis
OxytrichaTrifallax
ElphidiumMargaritaceum
ReticulomyxaFilosa
ChlorarachnionReptans
BigelowiellaNatans
PlasmodiophoraBrassicae
StygiellaIncarcerata
AndaluciaGodoyi
PharyngomonasKirbyi
NaegleriaGruberi
EuglenaGracilis
TrypanosomaBrucei
LeishmaniaMajor
BodoSaltans
ThecamonasTrahens
ChoanocystisSp
RaphidiophrysHeterophryoidea
PterocystisS
AcanthocystisS
ManchomonasBermudensis
PygsuiaBiforma
AcanthamoebaCastellanii
PyxidiculaOperculata
DictyosteliumDiscoideum
PolysphondyliumViolaceum
MantamonasSphyraenae
MantamonasPlastica
MantamonasVickermani
DiphylleiaRotans
RigifilaRamosa
NematostellaVectensis
RhopilemaEsculentum
SanderiaMalayensis
HydraVulgaris
ClytiaHemisphaerica
OctopusBimaculoides
LottiaGigantea
CrassostreaGigas
CapitellaTeleta
LamellibrachiaSatsuma
HelobdellaRobusta
LimulusPolyphemus
StegodyphusMimosarum
TriboliumCastaneum
DrosophilaMelanogaster
CalanusGlacialis
NeocalanusFlemingeri
StrongylocentrotusPurpuratus
LytechinusVariegatus
ThunnusOrientalis
SalmoSalar
DanioRerio
AnasPlatyrhynchos
GallusGallus
PelodiscusSinensis
AnolisCarolinensis
MusMusculus
HomoSapiens
XenopusTropicalis
TrichoplaxAdhaerens
AmphimedonQueenslandica
SalpingoecaUrceolata
DidymoecaCostata
ChromosphaeraPerkinsii
CreolimaxFragrantissima
SchizosaccharomycesJaponicus
SaccharomycesCerevisiae
NeurosporaCrassa
DelphinellaStrobiligena
AspergillusFumigatus
CystobasidiumMinutum
UstilagoMaydis
BatrachochytriumDendrobatidis
ChrysochromulinaParva
PrymnesiumParvum
EmilianiaHuxleyi
DiacronemaLutheri
GoniomonasPacifica
GuillardiaTheta
CyanophoraParadoxa
KlebsormidiumNitens
PhyscomitrellaPatens
NothocerosAenigmaticus
PsilotumNudum
PteridiumAquilinum
ZeaMays
OryzaSativa
ArabidopsisThaliana
GlycineMax
PyramimonasObovata
MicromonasCommoda
CrustomastixStigmatica
VolvoxCarteri
ChlamydomonasReinhardtii
CoccomyxaSubellipsoidea
GracilariopsisChorda
ChondrusCrispus
CompsopogonCaeruleus
RhodellaViolacea
CyanidioschyzonMerolae
MonocercomonoidesExilis
BlattamonasNauphoetae
ParatrimastixPyriformis
TritrichomonasFoetus
TrichomonasVaginalis
ChilomastixCuspidata
GiardiaIntestinalis
TrepomonasSp
SpironucleusSalmonicida
GefionellaOkellyi
MalawimonasJakobiformis
NutomonasLonga
AncyromonasSigmoides
FabomonasTropica
HemimastixKukwesjijk
SpironemaSp

**Supplementary Figure S3**

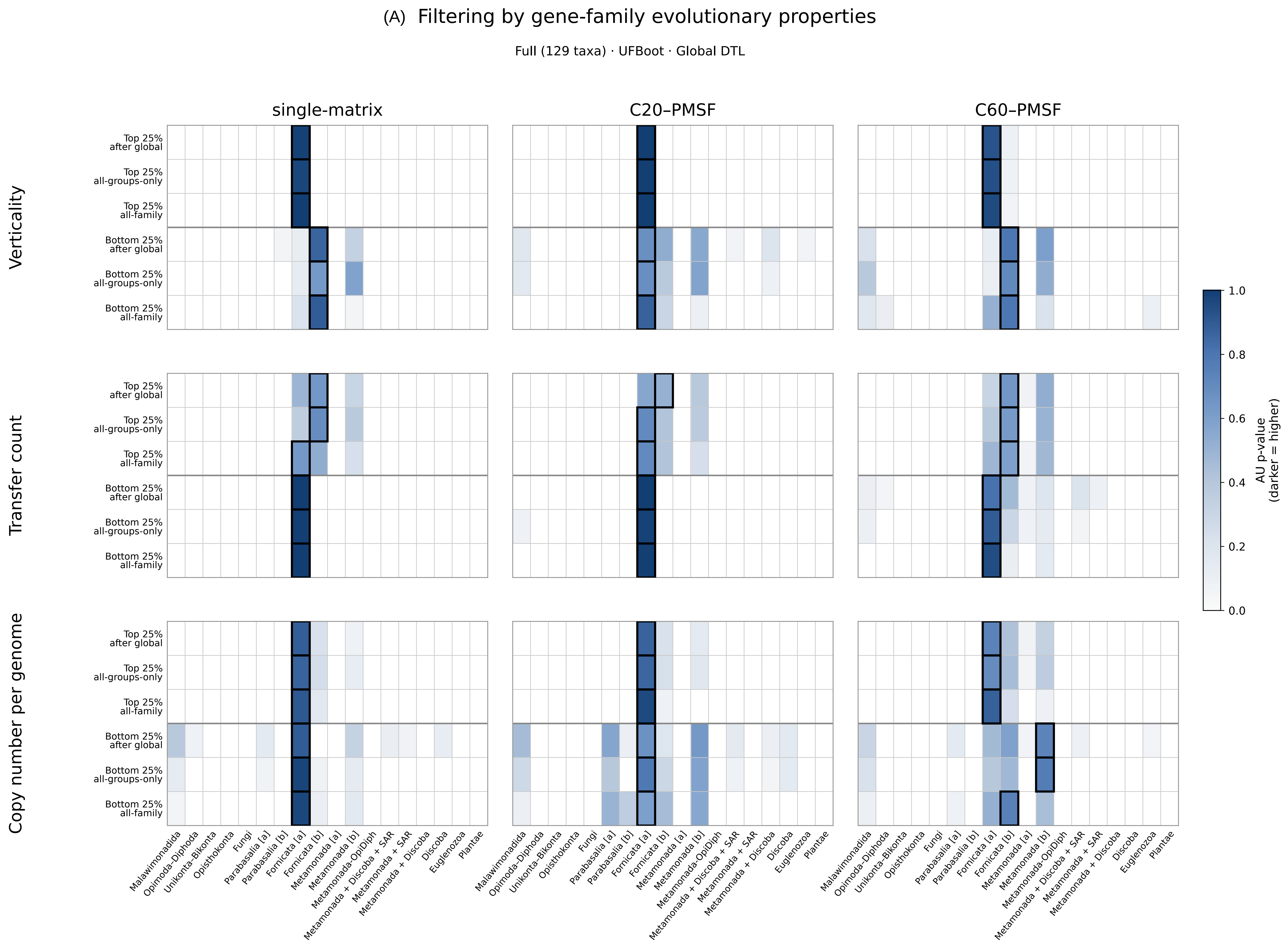


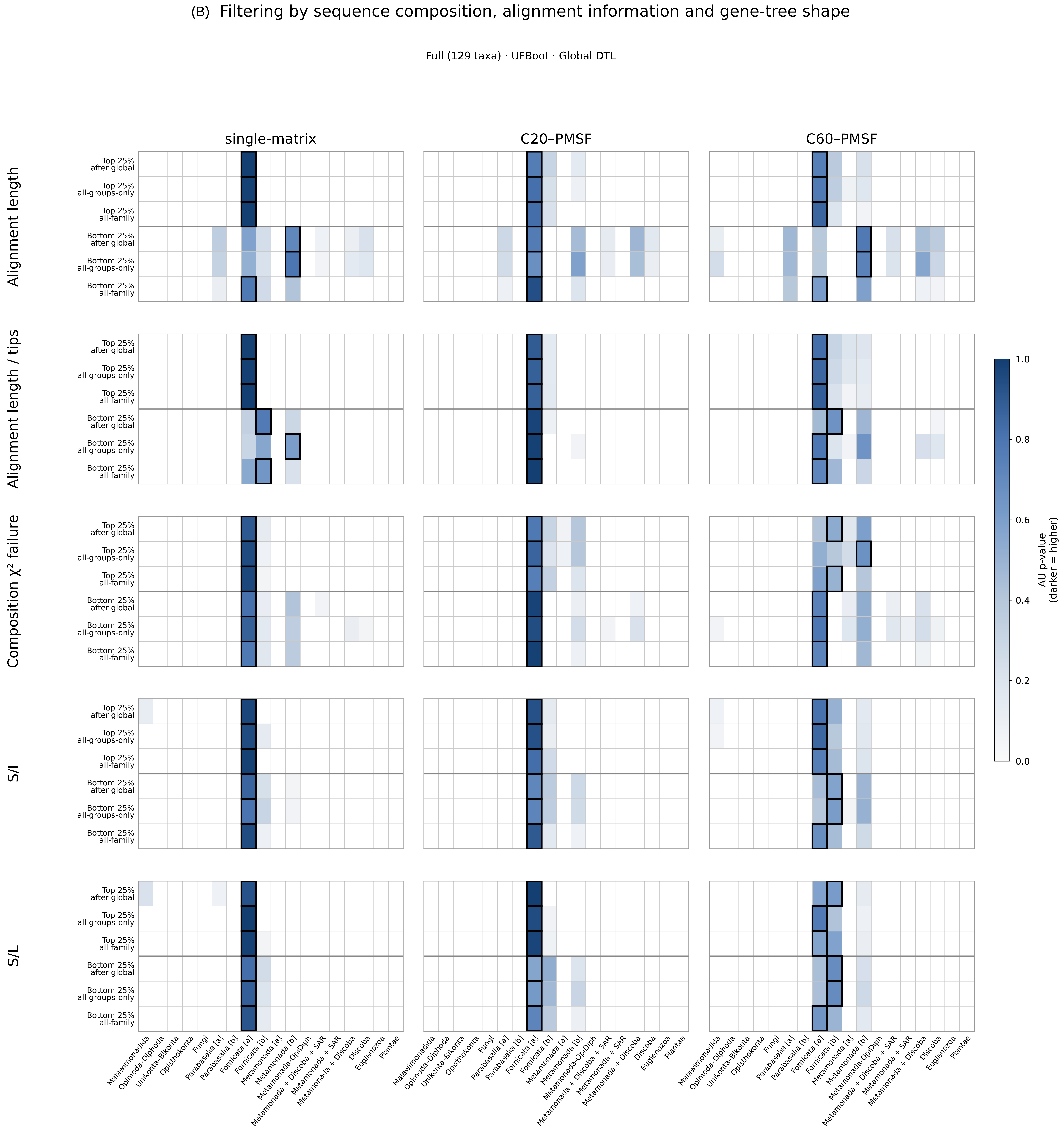


**Supplementary Figure S3.** Sensitivity of eukaryotic root support to gene-family filtering in the Full 129-taxon dataset under global DTL. (A) Filtering by gene-family evolutionary properties: verticality, inferred transfer count and copy number per represented genome. (B) Filtering by sequence- and tree-based properties: alignment length, tip-normalised alignment length, composition chi-squared failure rate, S/I and S/L. Heatmaps show AU support for alternative rooted species trees under three UFBoot gene-tree frameworks (single-matrix, C20–PMSF and C60–PMSF). Within each metric, the upper and lower three rows show high- and low-quartile subsets, respectively, selected by all-groups filtering after global ranking, all-groups-only ranking, and all-family ranking. Darker blue indicates higher AU support; white indicates AU < 0.05. Black outlines mark the rooted tree with the highest summed reconciliation log-likelihood for each subset and gene-tree framework. The full dataset compares 18 rooted species trees, and the reduced dataset compares 12.

**Supplementary Figure S4**

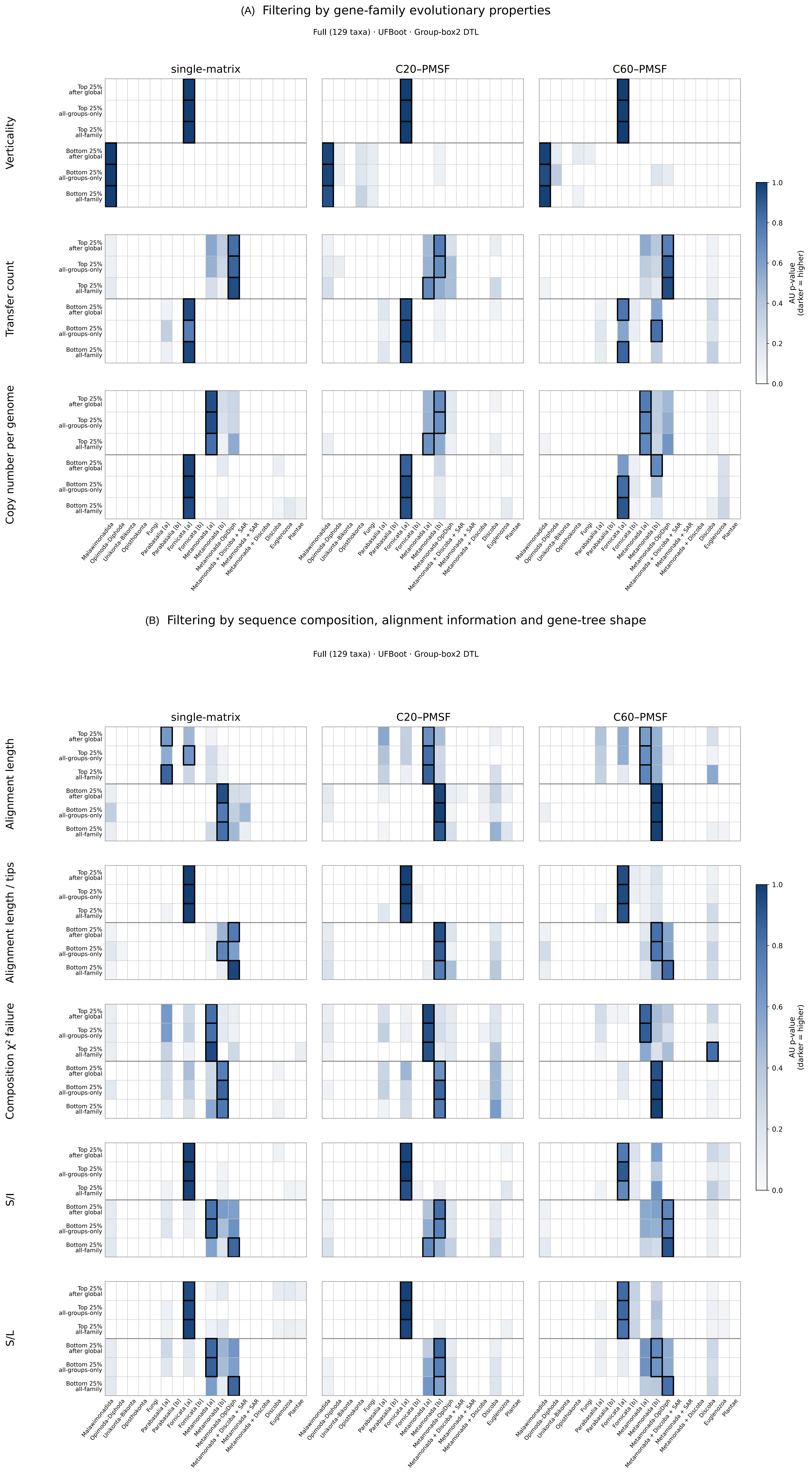


**Supplementary Figure S4.** Sensitivity of eukaryotic root support to gene-family filtering in the Full 129-taxon dataset under bounded lineage-specific DTL (box2). (A) Filtering by gene-family evolutionary properties: verticality, inferred transfer count and copy number per represented genome. (B) Filtering by sequence- and tree-based properties: alignment length, tip-normalised alignment length, composition chi-squared failure rate, S/I and S/L. Heatmaps show AU support for alternative rooted species trees under three UFBoot gene-tree frameworks (single-matrix, C20–PMSF and C60–PMSF). Within each metric, the upper and lower three rows show high- and low-quartile subsets, respectively, selected by all-groups filtering after global ranking, all-groups-only ranking, and all-family ranking. Darker blue indicates higher AU support; white indicates AU < 0.05. Black outlines mark the rooted tree with the highest summed reconciliation log-likelihood for each subset and gene-tree framework. The full dataset compares 18 rooted species trees, and the reduced dataset compares 12.

**Supplementary Figure S5**

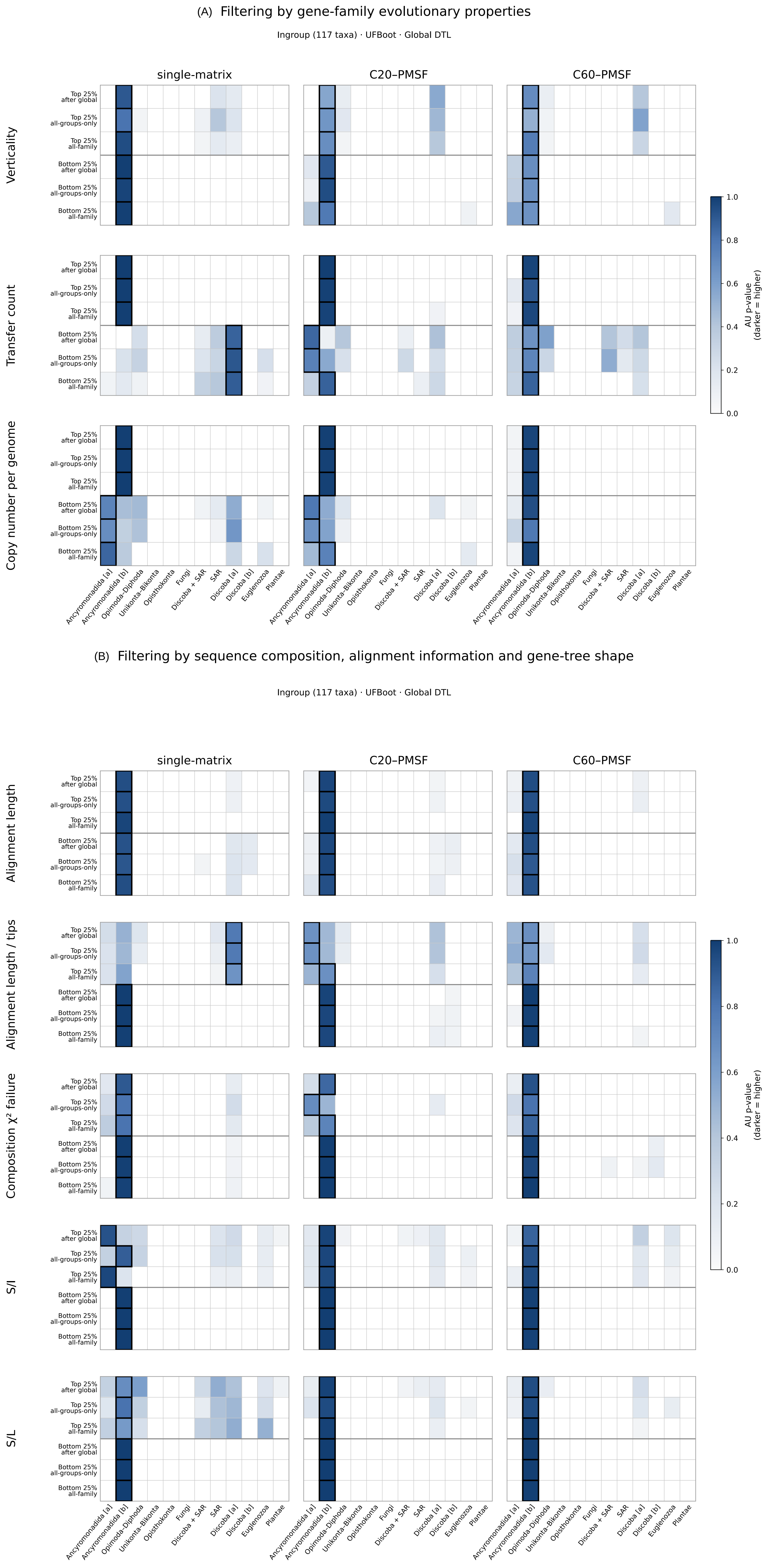


**Supplementary Figure S5.** Sensitivity of eukaryotic root support to gene-family filtering in the Reduced 117-taxon dataset under global DTL. (A) Filtering by gene-family evolutionary properties: verticality, inferred transfer count and copy number per represented genome. (B) Filtering by sequence- and tree-based properties: alignment length, tip-normalised alignment length, composition chi-squared failure rate, S/I and S/L. Heatmaps show AU support for alternative rooted species trees under three UFBoot gene-tree frameworks (single-matrix, C20–PMSF and C60–PMSF). Within each metric, the upper and lower three rows show high- and low-quartile subsets, respectively, selected by all-groups filtering after global ranking, all-groups-only ranking, and all-family ranking. Darker blue indicates higher AU support; white indicates AU $< 0.05$. Black outlines mark the rooted tree with the highest summed reconciliation log-likelihood for each subset and gene-tree framework. The full dataset compares 18 rooted species trees, and the reduced dataset compares 12.

### Supplementary Figure S6

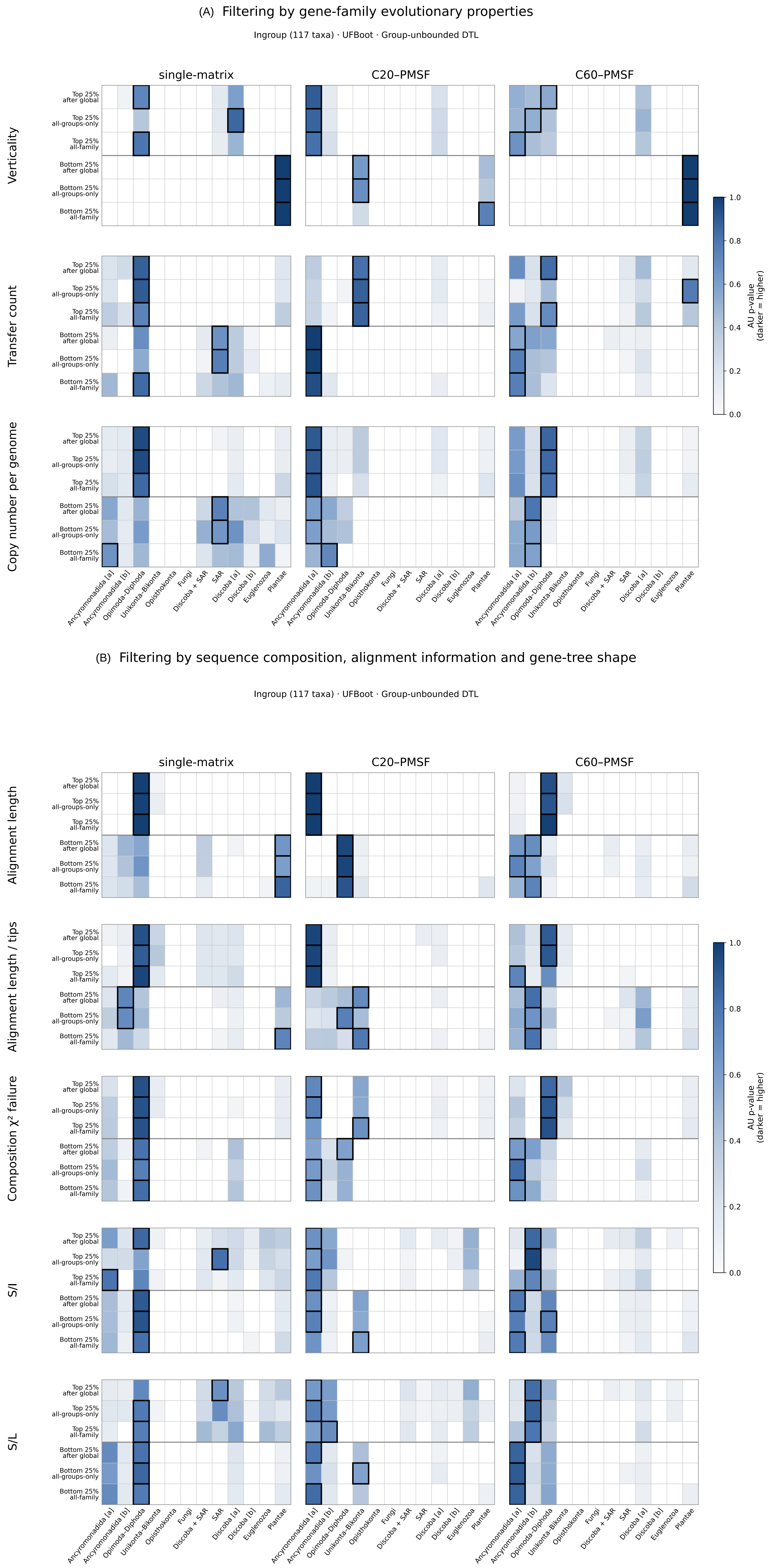


**Supplementary Figure S6.** Sensitivity of eukaryotic root support to gene-family filtering in the Reduced 117-taxon dataset under unbounded lineage-specific DTL. (A) Filtering by gene-family evolutionary properties: verticality, inferred transfer count and copy number per represented genome. (B) Filtering by sequence- and tree-based properties: alignment length, tip-normalised alignment length, composition chi-squared failure rate, S/I and S/L. Heatmaps show AU support for alternative rooted species trees under three UFBoot gene-tree frameworks (single-matrix, C20–PMSF and C60–PMSF). Within each metric, the upper and lower three rows show high- and low-quartile subsets, respectively, selected by all-groups filtering after global ranking, all-groups-only ranking, and all-family ranking. Darker blue indicates higher AU support; white indicates AU < 0.05. Black outlines mark the rooted tree with the highest summed reconciliation log-likelihood for each subset and gene-tree framework. The full dataset compares 18 rooted species trees, and the reduced dataset compares 12.

### Supplementary Figure S7

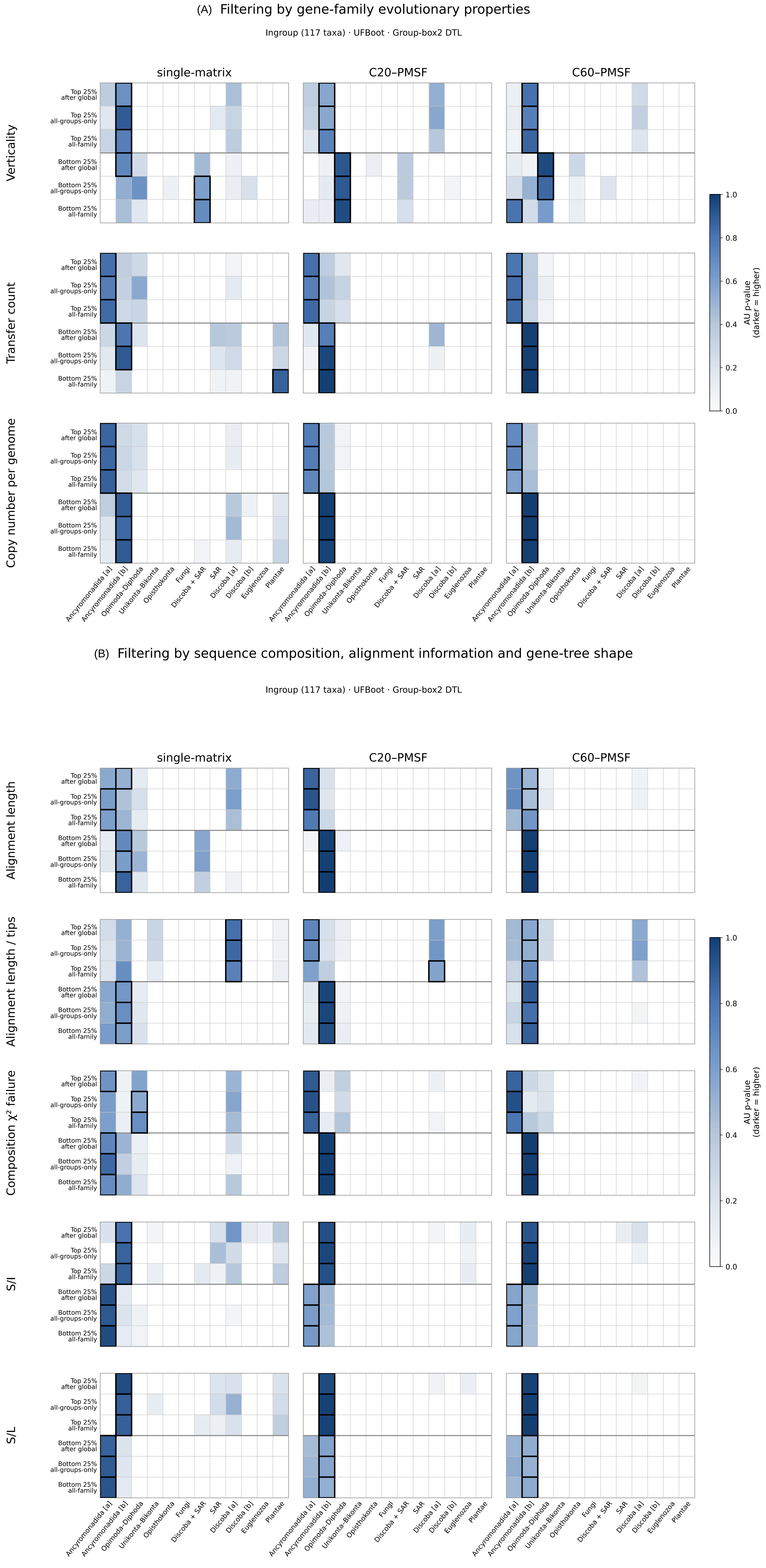


**Supplementary Figure S7.** Sensitivity of eukaryotic root support to gene-family filtering in the Reduced 117-taxon dataset under bounded lineage-specific DTL (box2). (A) Filtering by gene-family evolutionary properties: verticality, inferred transfer count and copy number per represented genome. (B) Filtering by sequence- and tree-based properties: alignment length, tip-normalised alignment length, composition chi-squared failure rate, S/I and S/L. Heatmaps show AU support for alternative rooted species trees under three UFBoot gene-tree frameworks (single-matrix, C20–PMSF and C60–PMSF). Within each metric, the upper and lower three rows show high- and low-quartile subsets, respectively, selected by all-groups filtering after global ranking, all-groups-only ranking, and all-family ranking. Darker blue indicates higher AU support; white indicates AU < 0.05. Black outlines mark the rooted tree with the highest summed reconciliation log-likelihood for each subset and gene-tree framework. The full dataset compares 18 rooted species trees, and the reduced dataset compares 12.

**(A) Gene-family representation and relative root support — Full (129 taxa)**

C20/PMSF UFBoot | Global DTL

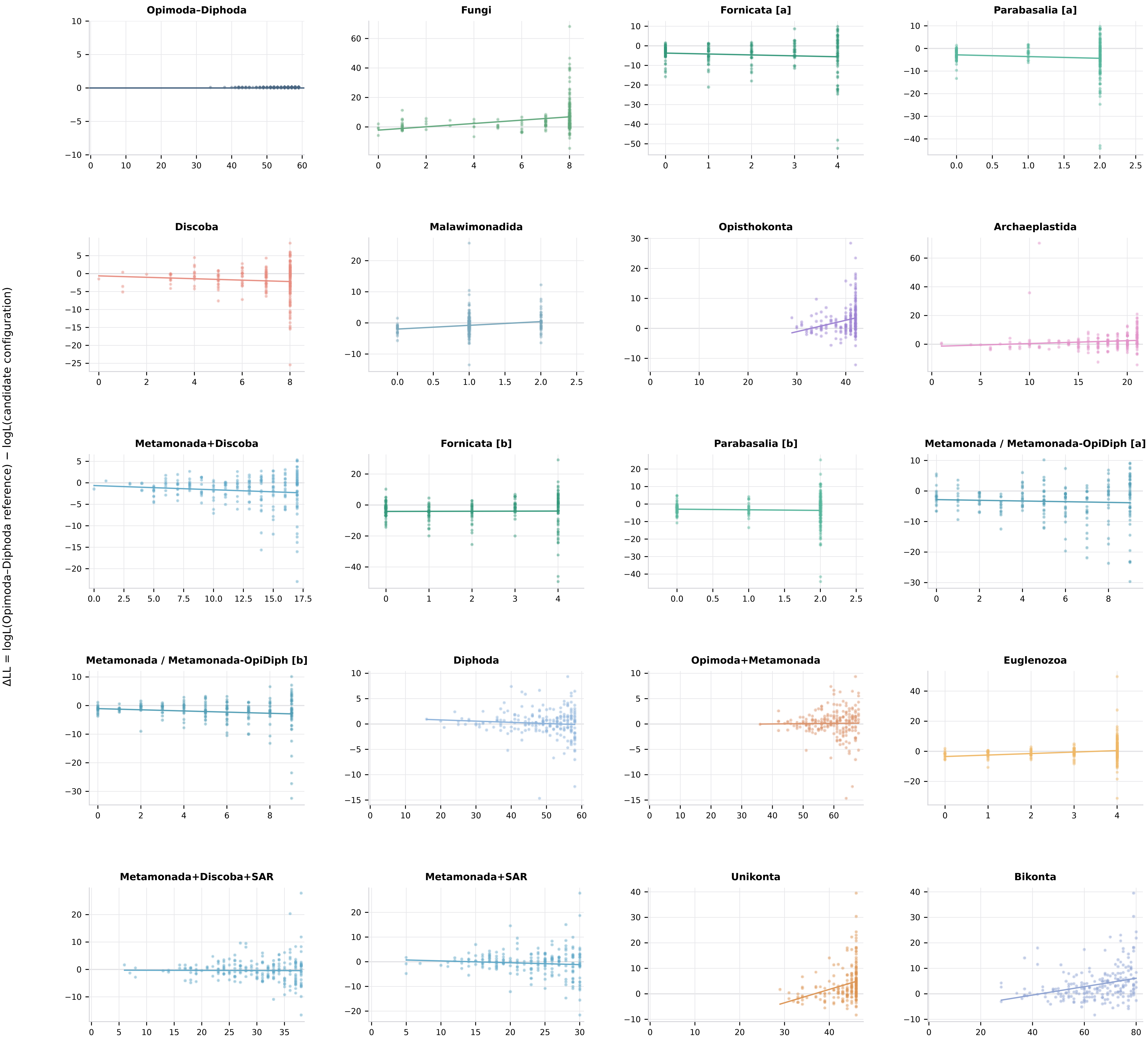

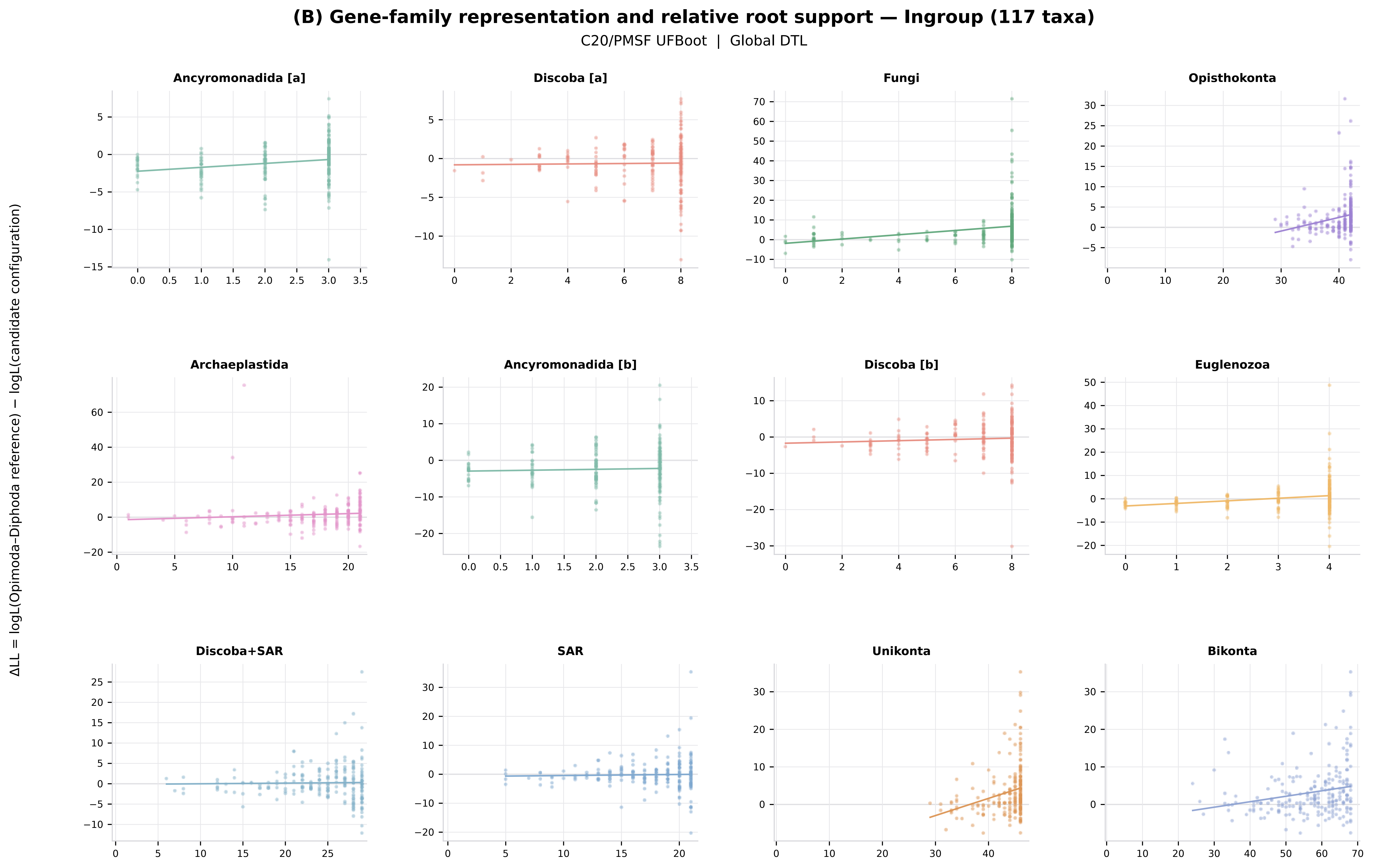

(B) Gene-family representation and relative root support — Ingroup (117 taxa)
C20/PMSF UFBoot | Global DTL
ΔLL = logL(Opimoda–Diphoda reference) − logL(candidate configuration)
Ancyromonadida [a]
Discoba [a]
Fungi
Opisthokonta
Archaeplastida
Ancyromonadida [b]
Discoba [b]
Euglenozoa
Discoba+SAR
SAR
Unikonta
Bikonta

**(A) Gene-family representation and relative root support — Full (129 taxa)**

single-matrix (MFP) UFBoot | Global DTL

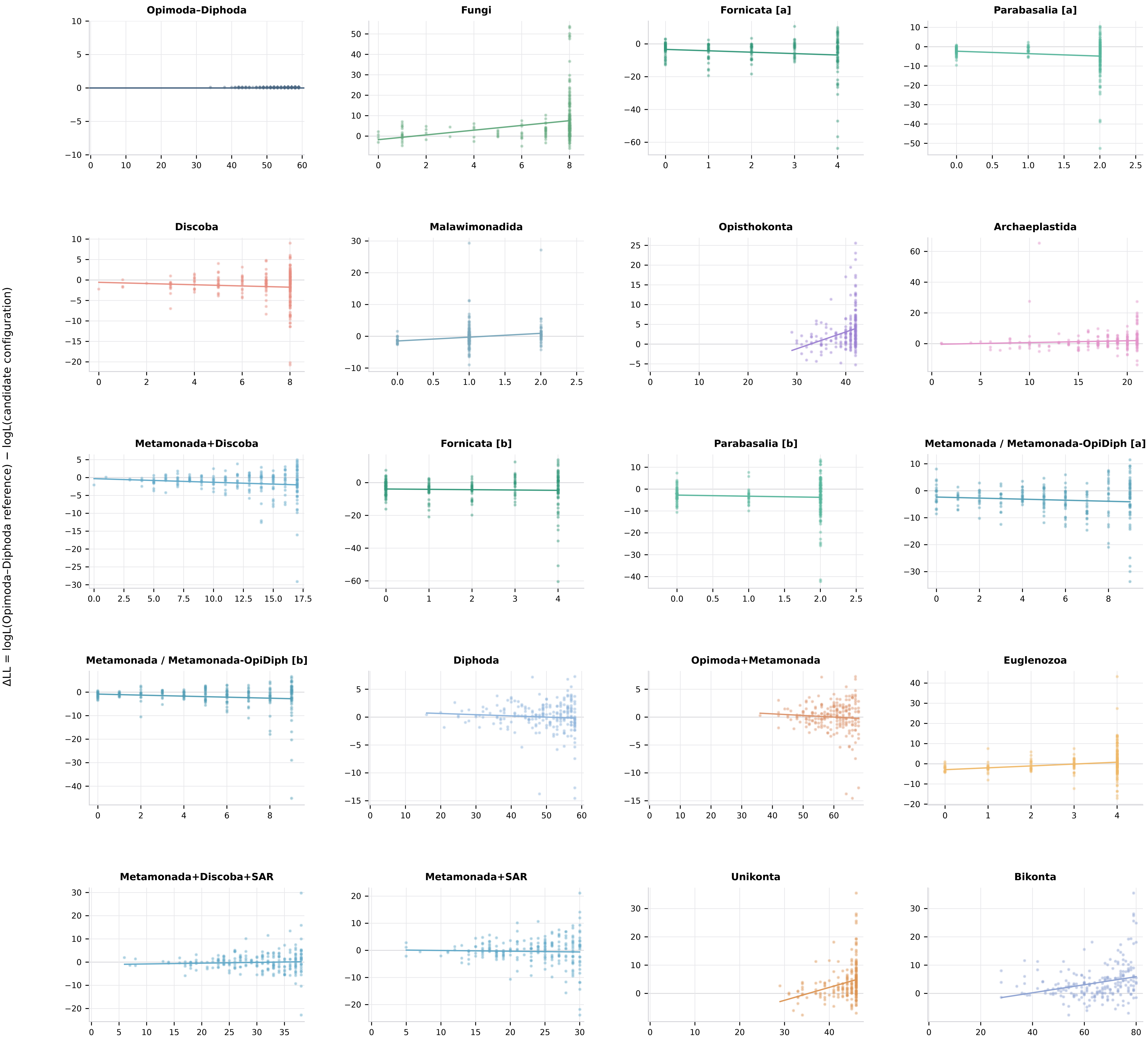

**(A) Gene-family representation and relative root support — Full (129 taxa)**

single-matrix (MFP) UFBoot | Group-specific DTL (rates bounded at 2)

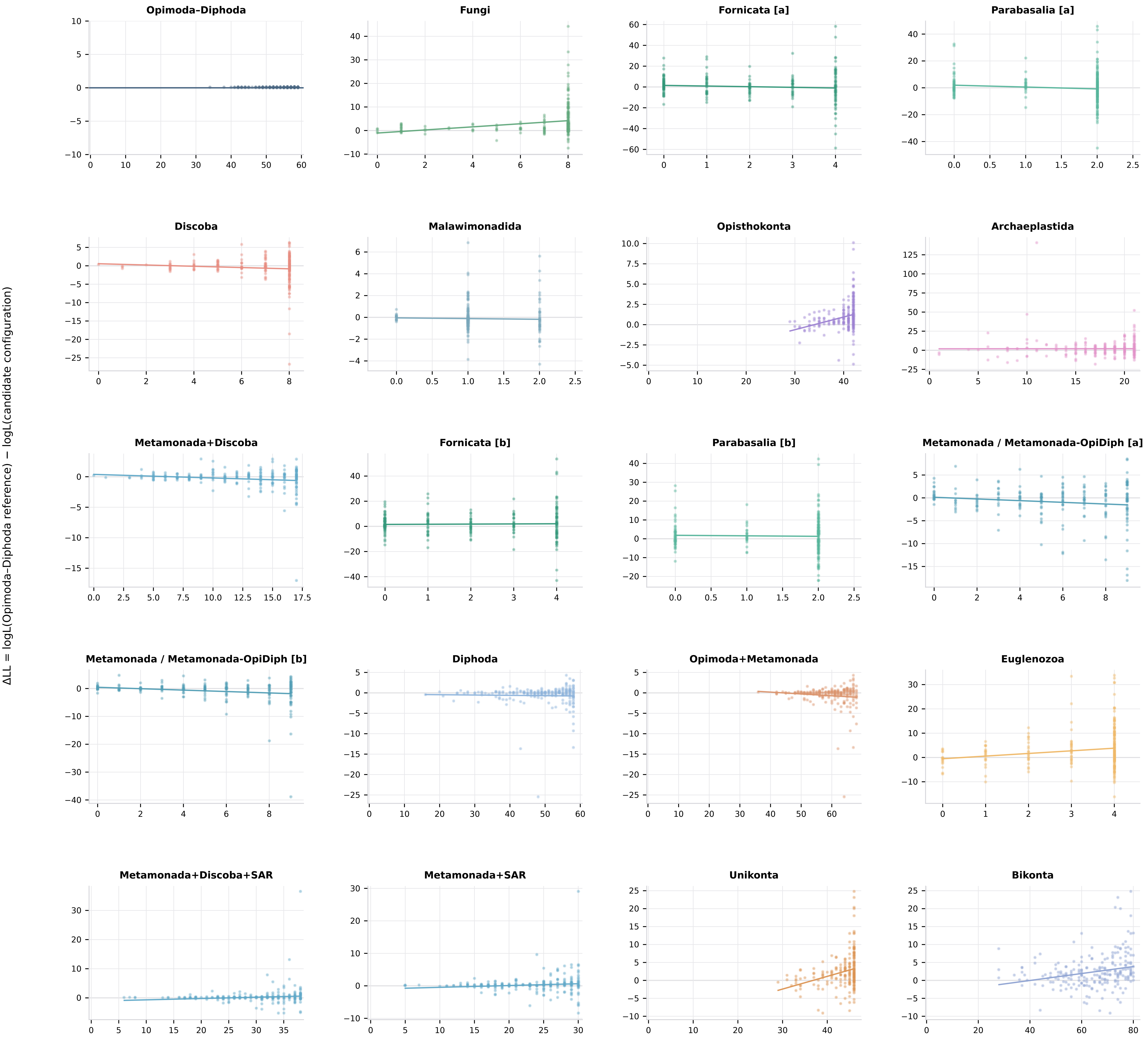

**(A) Gene-family representation and relative root support — Full (129 taxa)**

single-matrix (MFP) UFBoot | Group-specific DTL (unbounded)

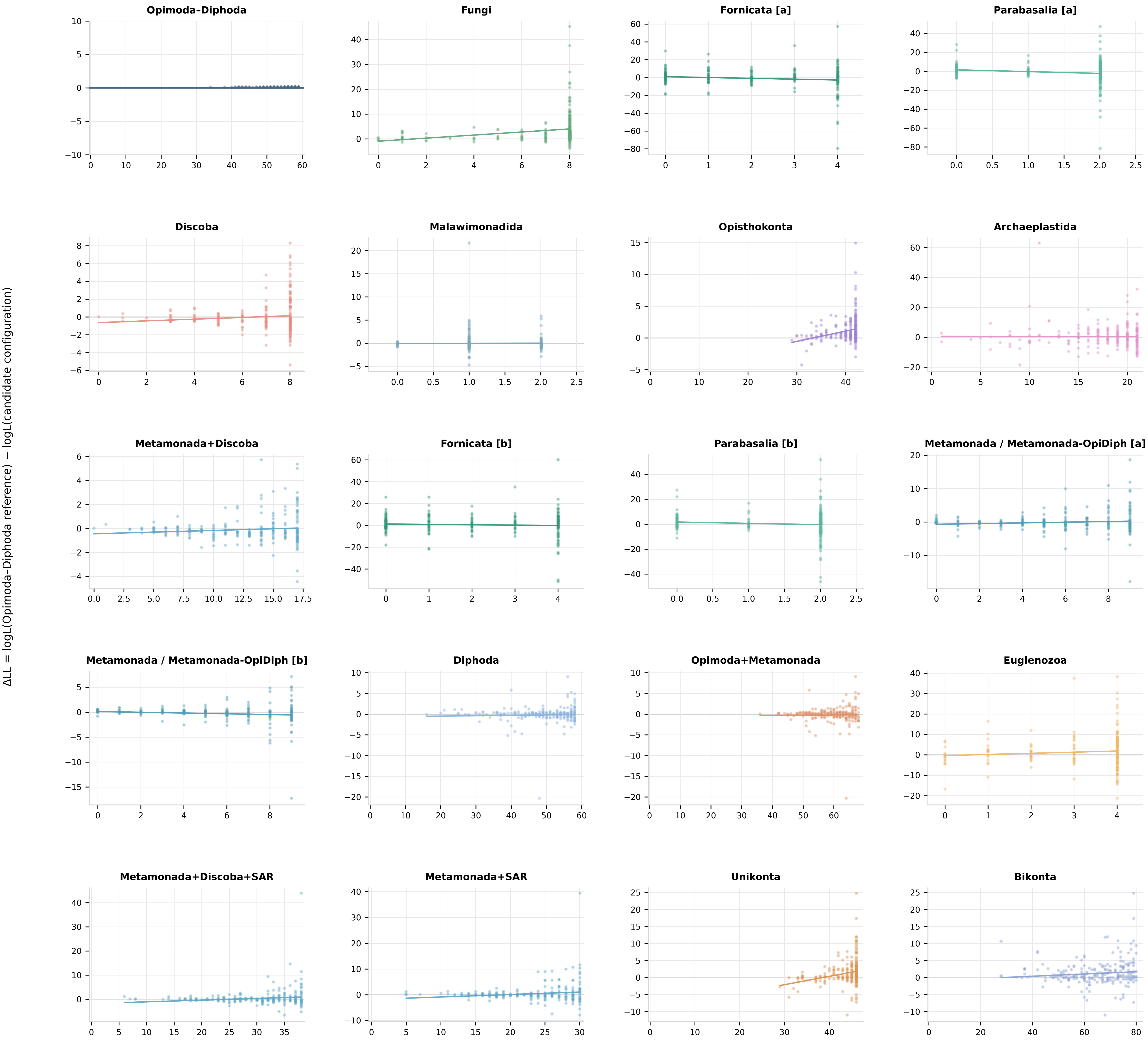

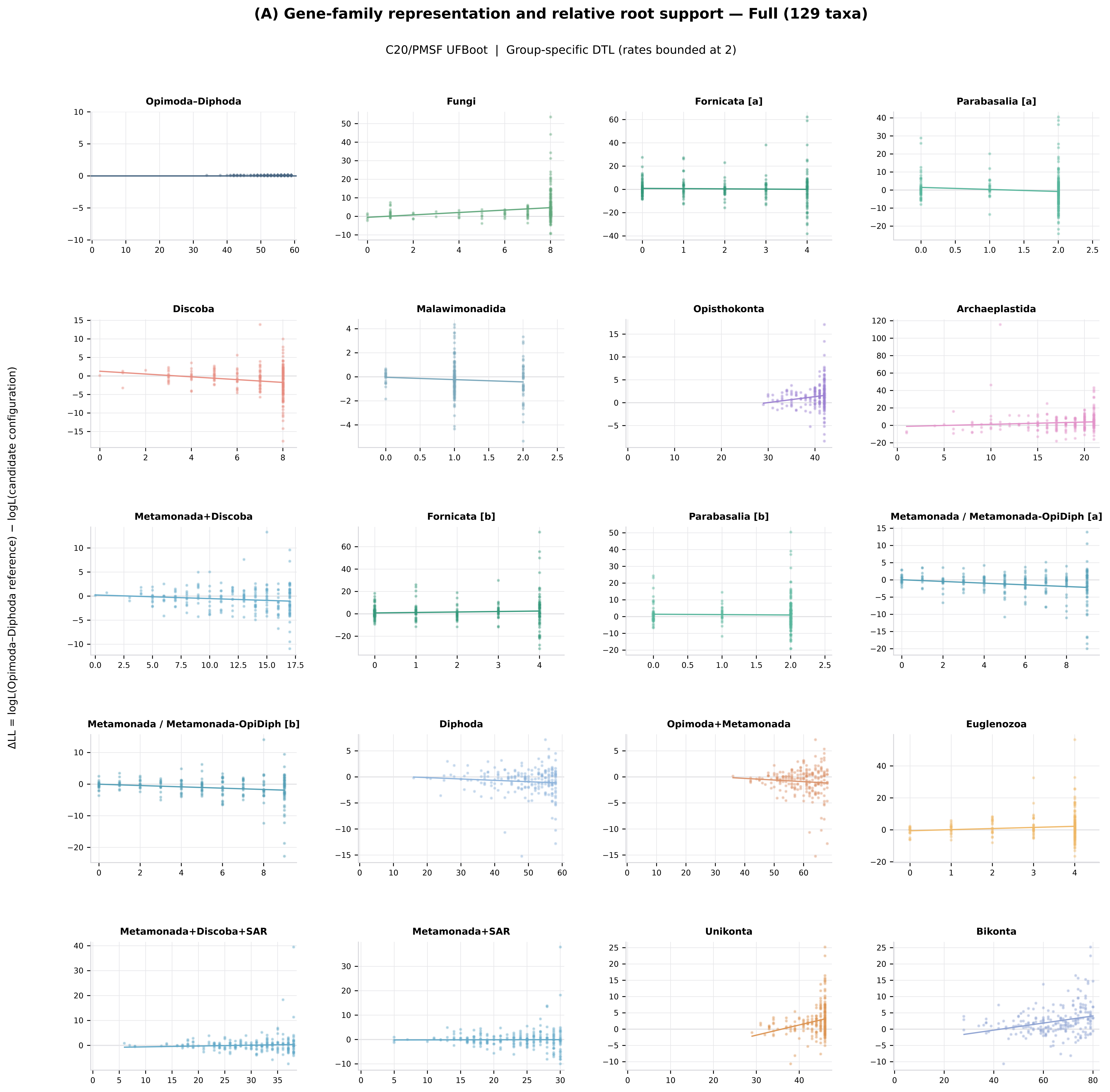

(A) Gene-family representation and relative root support — Full (129 taxa)
C20/PMSF UFBoot | Group-specific DTL (rates bounded at 2)
ΔLL = logL(Opimoda–Diphoda reference) − logL(candidate configuration)
Opimoda–Diphoda
Fungi
Fornicata [a]
Parabasalia [a]
Discoba
Malawimonadida
Opisthokonta
Archaeplastida
Metamonada+Discoba
Fornicata [b]
Parabasalia [b]
Metamonada / Metamonada-OpiDiph [a]
Metamonada / Metamonada-OpiDiph [b]
Diphoda
Opimoda+Metamonada
Euglenozoa
Metamonada+Discoba+SAR
Metamonada+SAR
Unikonta
Bikonta

**(A) Gene-family representation and relative root support — Full (129 taxa)**

C20/PMSF UFBoot | Group-specific DTL (unbounded)

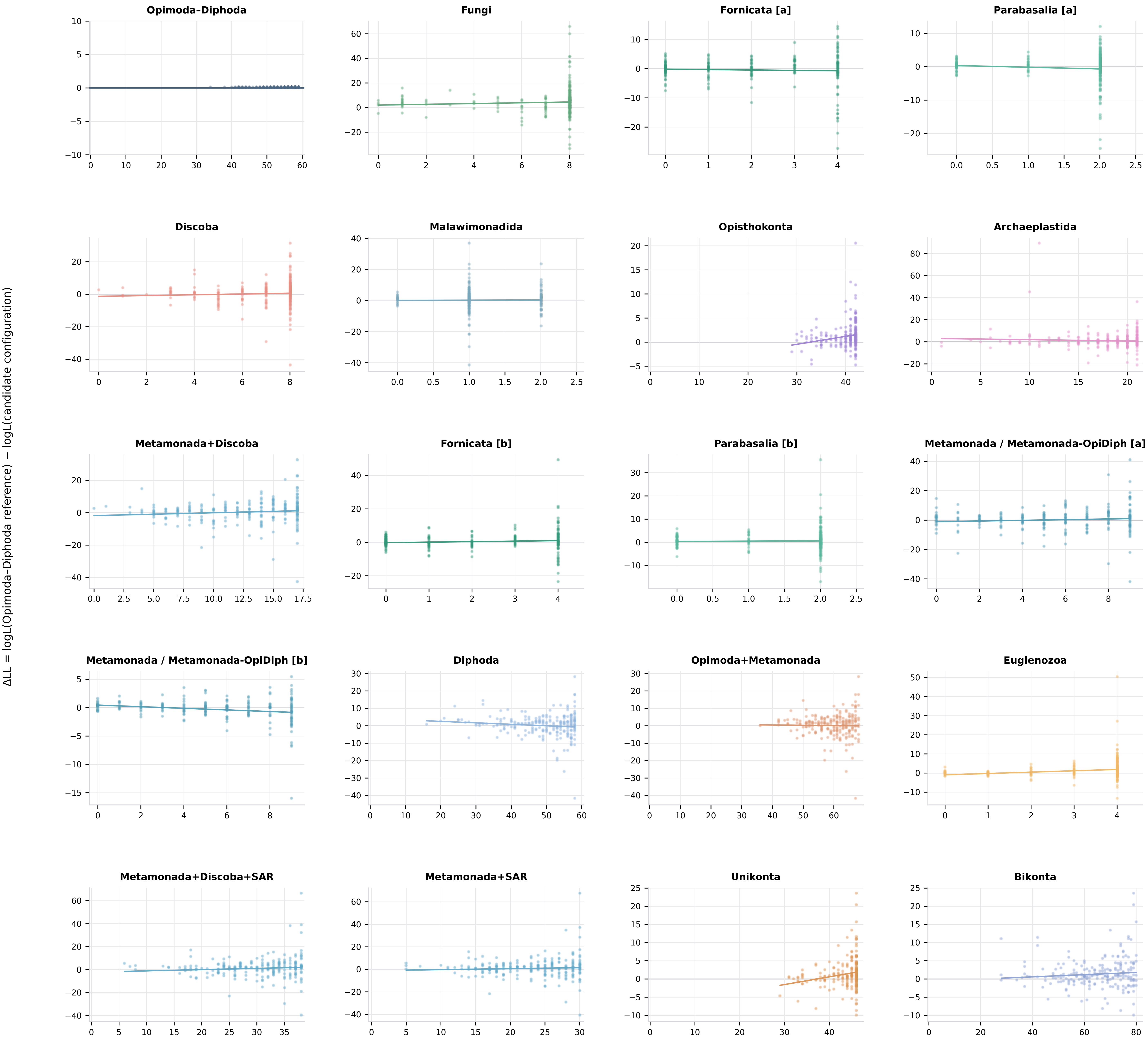

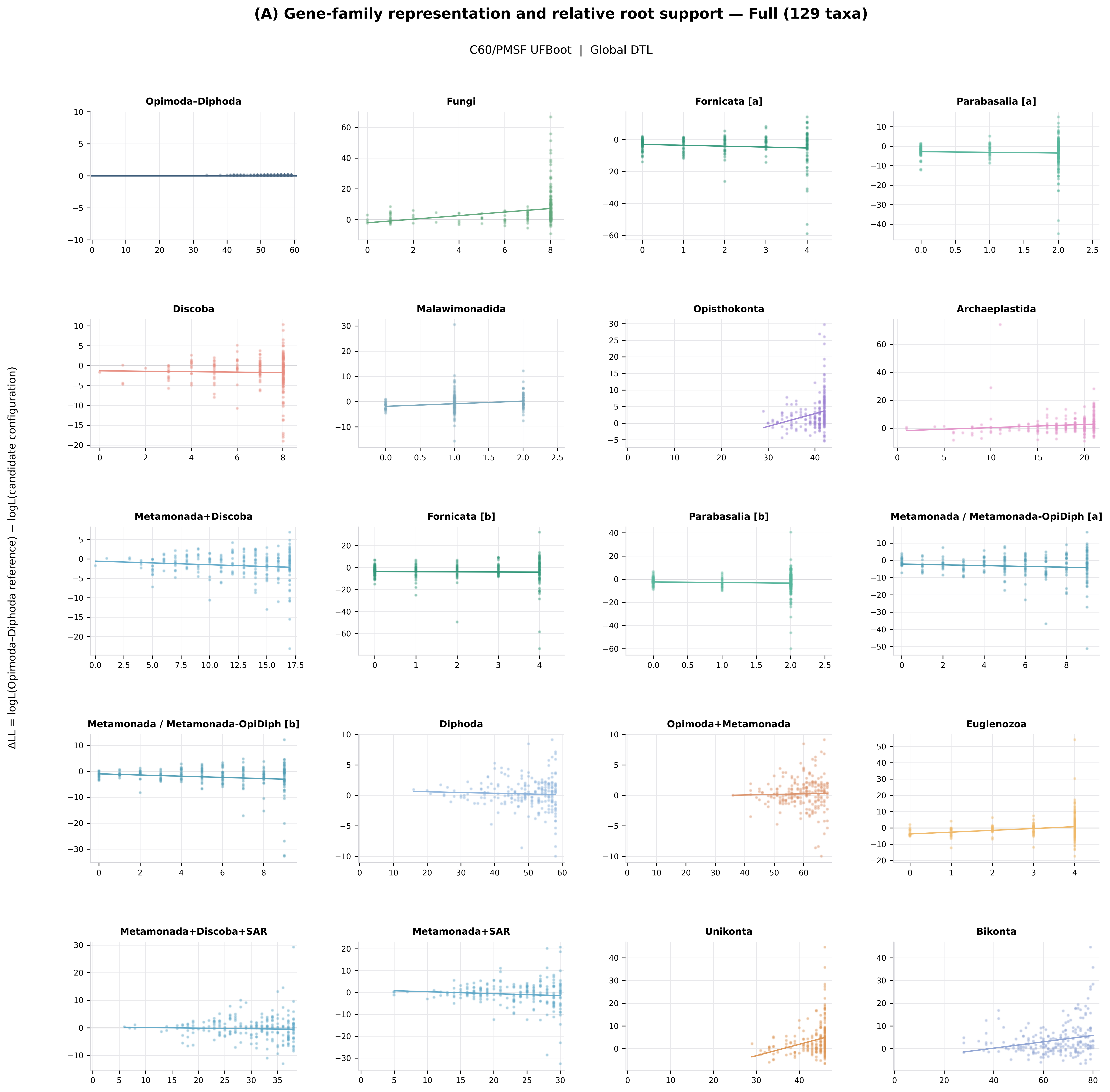

(A) Gene-family representation and relative root support — Full (129 taxa)
C60/PMSF UFBoot | Global DTL
ΔLL = logL(Opimoda–Diphoda reference) − logL(candidate configuration)
Opimoda–Diphoda
Fungi
Fornicata [a]
Parabasalia [a]
Discoba
Malawimonadida
Opisthokonta
Archaeplastida
Metamonada+Discoba
Fornicata [b]
Parabasalia [b]
Metamonada / Metamonada-OpiDiph [a]
Metamonada / Metamonada-OpiDiph [b]
Diphoda
Opimoda+Metamonada
Euglenozoa
Metamonada+Discoba+SAR
Metamonada+SAR
Unikonta
Bikonta

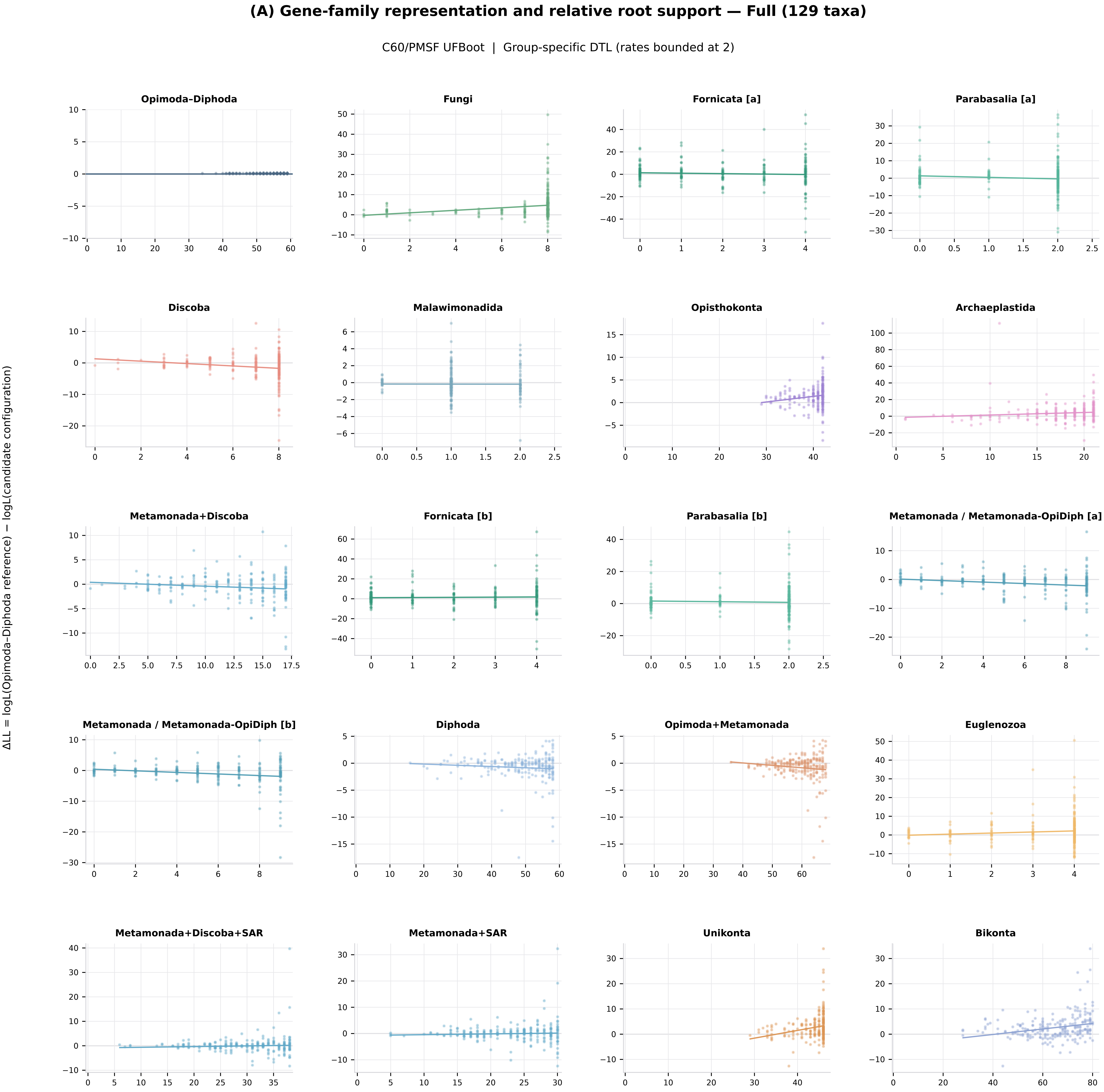
(A) Gene-family representation and relative root support — Full (129 taxa)
C60/PMSF UFBoot | Group-specific DTL (rates bounded at 2)
ΔLL = logL(Opimoda–Diphoda reference) − logL(candidate configuration)
Opimoda–Diphoda
Fungi
Fornicata [a]
Parabasalia [a]
Discoba
Malawimonadida
Opisthokonta
Archaeplastida
Metamonada+Discoba
Fornicata [b]
Parabasalia [b]
Metamonada / Metamonada-OpiDiph [a]
Metamonada / Metamonada-OpiDiph [b]
Diphoda
Opimoda+Metamonada
Euglenozoa
Metamonada+Discoba+SAR
Metamonada+SAR
Unikonta
Bikonta

### (A) Gene-family representation and relative root support — Full (129 taxa)

C60/PMSF UFBoot | Group-specific DTL (unbounded)

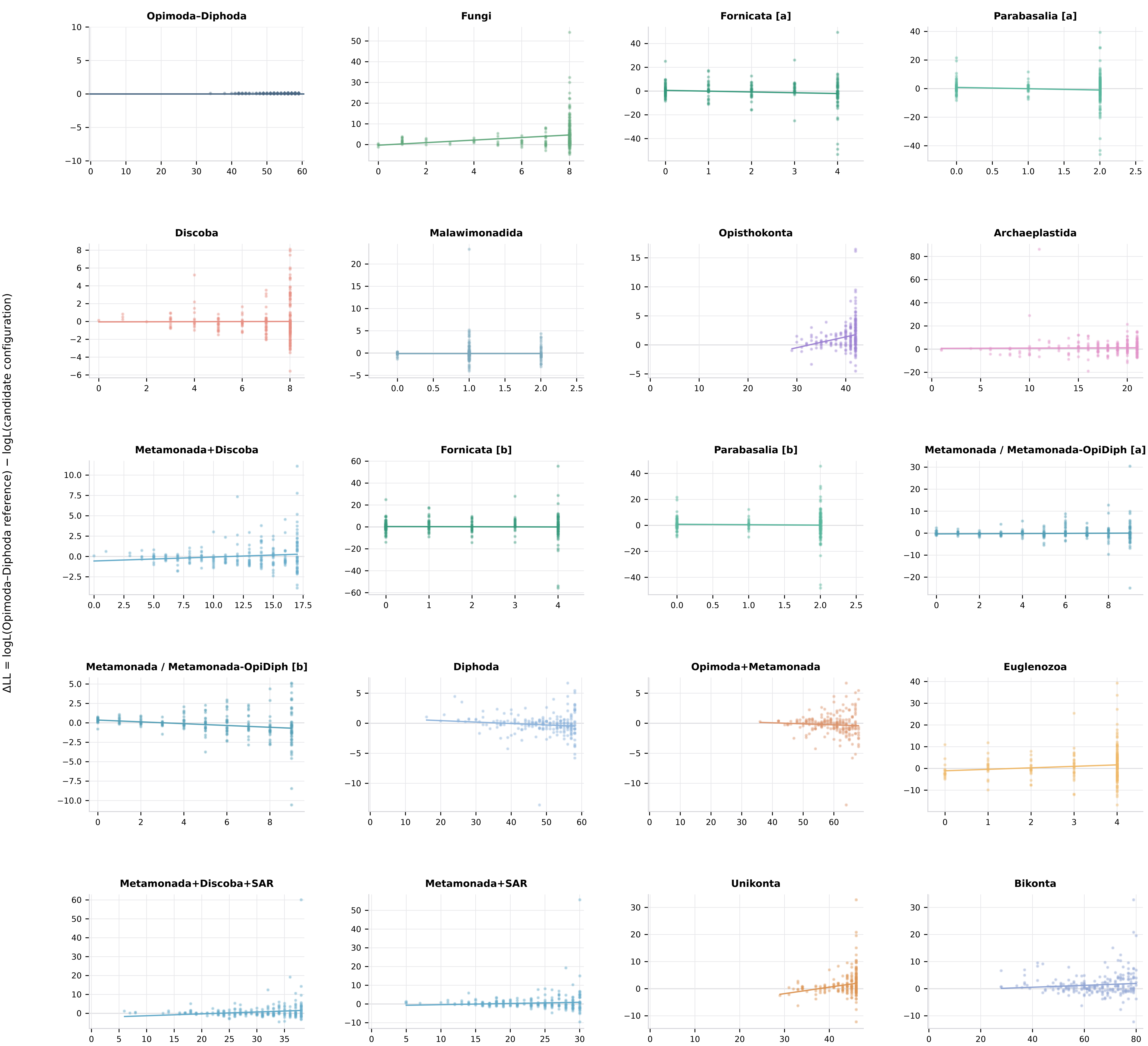

**(B) Gene-family representation and relative root support — Ingroup (117 taxa)**

single-matrix (MFP) UFBoot | Global DTL

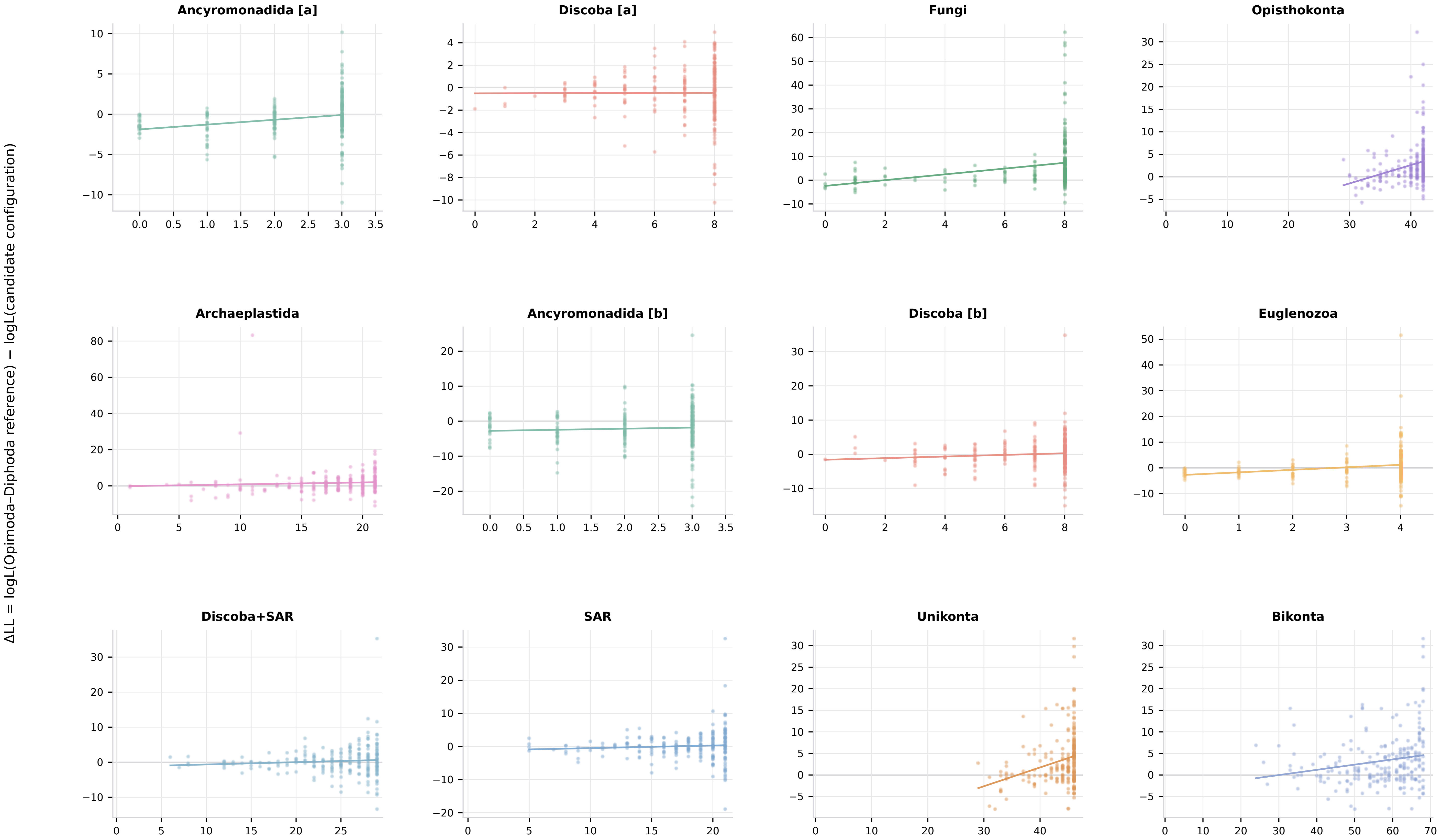

**(B) Gene-family representation and relative root support — Ingroup (117 taxa)**

single-matrix (MFP) UFBoot | Group-specific DTL (rates bounded at 2)

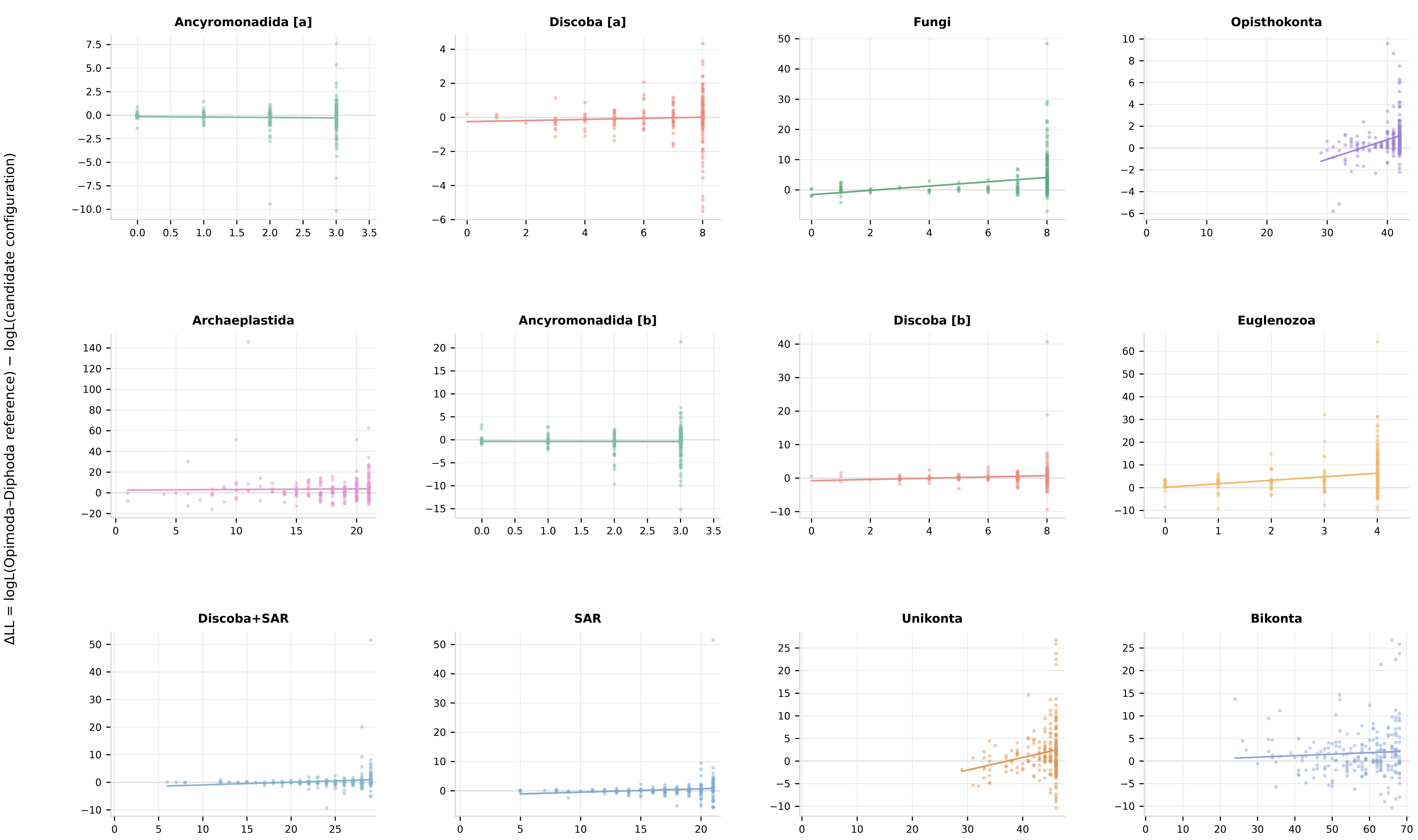

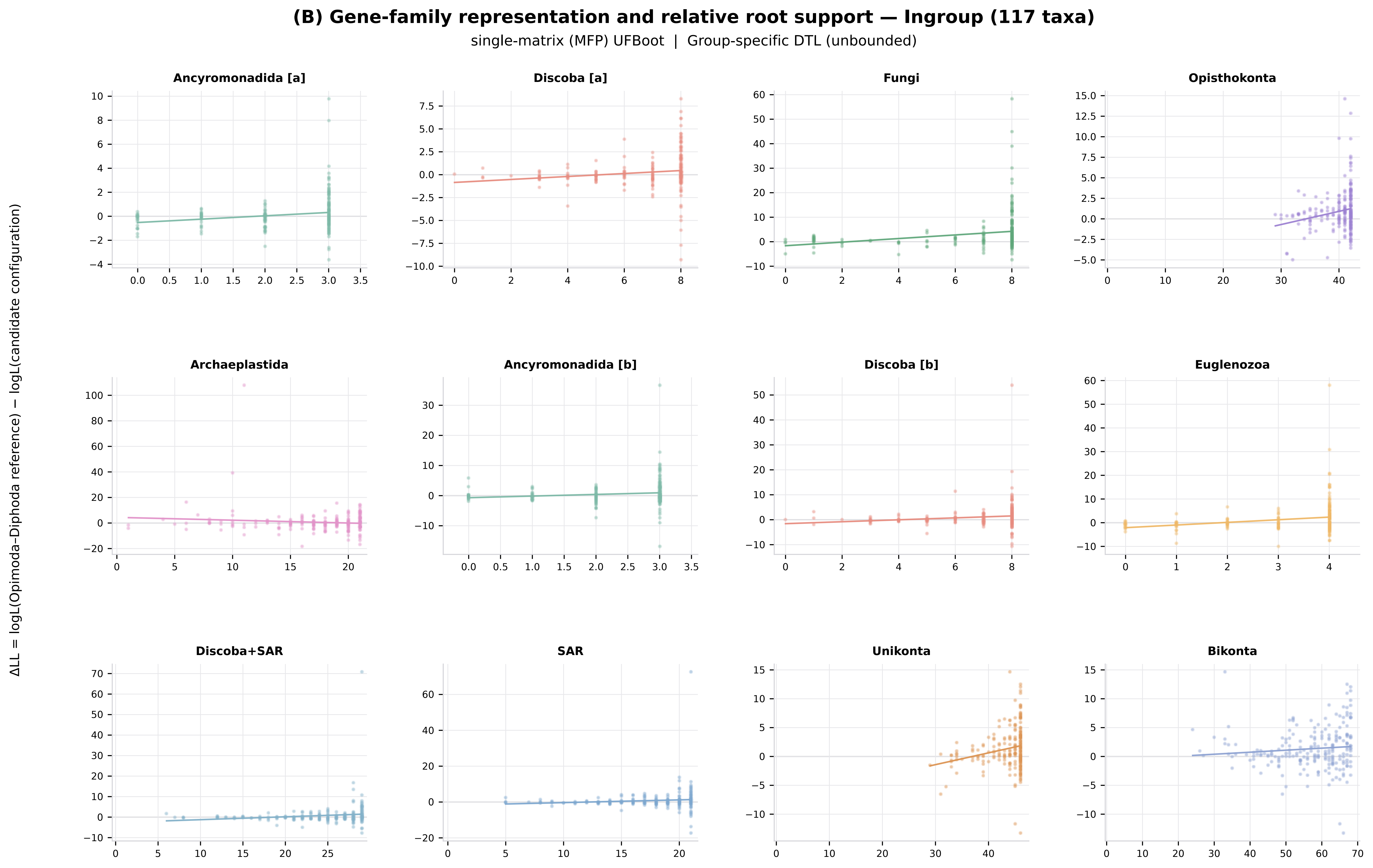

(B) Gene-family representation and relative root support — Ingroup (117 taxa)
single-matrix (MFP) UFBoot | Group-specific DTL (unbounded)
ΔLL = logL(Opimoda–Diphoda reference) − logL(candidate configuration)
Ancyromonadida [a]
Discoba [a]
Fungi
Opisthokonta
Archaeplastida
Ancyromonadida [b]
Discoba [b]
Euglenozoa
Discoba+SAR
SAR
Unikonta
Bikonta

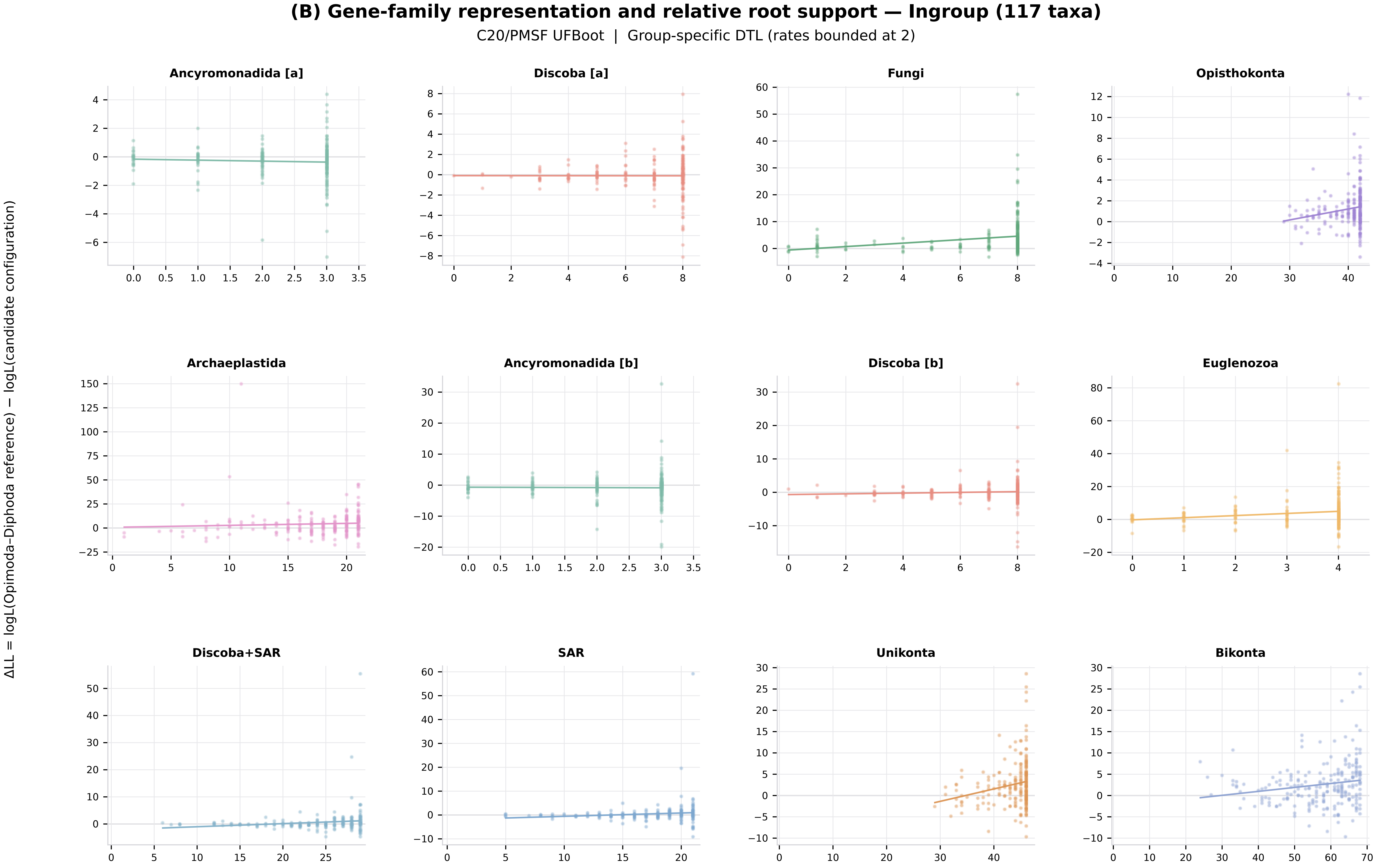
(B) Gene-family representation and relative root support — Ingroup (117 taxa)
C20/PMSF UFBoot | Group-specific DTL (rates bounded at 2)
ΔLL = logL(Opimoda–Diphoda reference) − logL(candidate configuration)
Ancyromonadida [a]
Discoba [a]
Fungi
Opisthokonta
Archaeplastida
Ancyromonadida [b]
Discoba [b]
Euglenozoa
Discoba+SAR
SAR
Unikonta
Bikonta

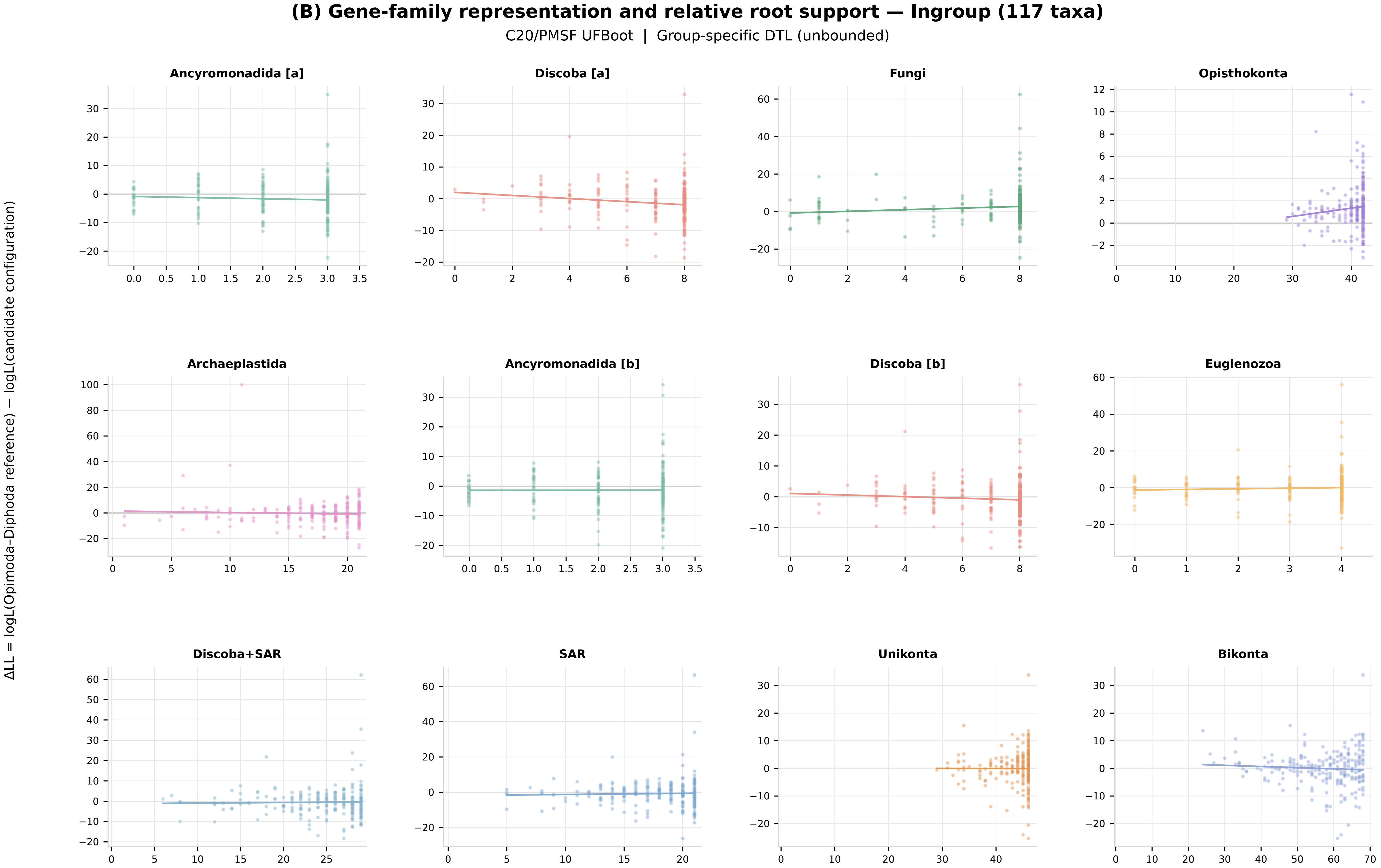
(B) Gene-family representation and relative root support — Ingroup (117 taxa)
C20/PMSF UFBoot | Group-specific DTL (unbounded)
ΔLL = logL(Opimoda–Diphoda reference) − logL(candidate configuration)
Ancyromonadida [a]
Discoba [a]
Fungi
Opisthokonta
Archaeplastida
Ancyromonadida [b]
Discoba [b]
Euglenozoa
Discoba+SAR
SAR
Unikonta
Bikonta

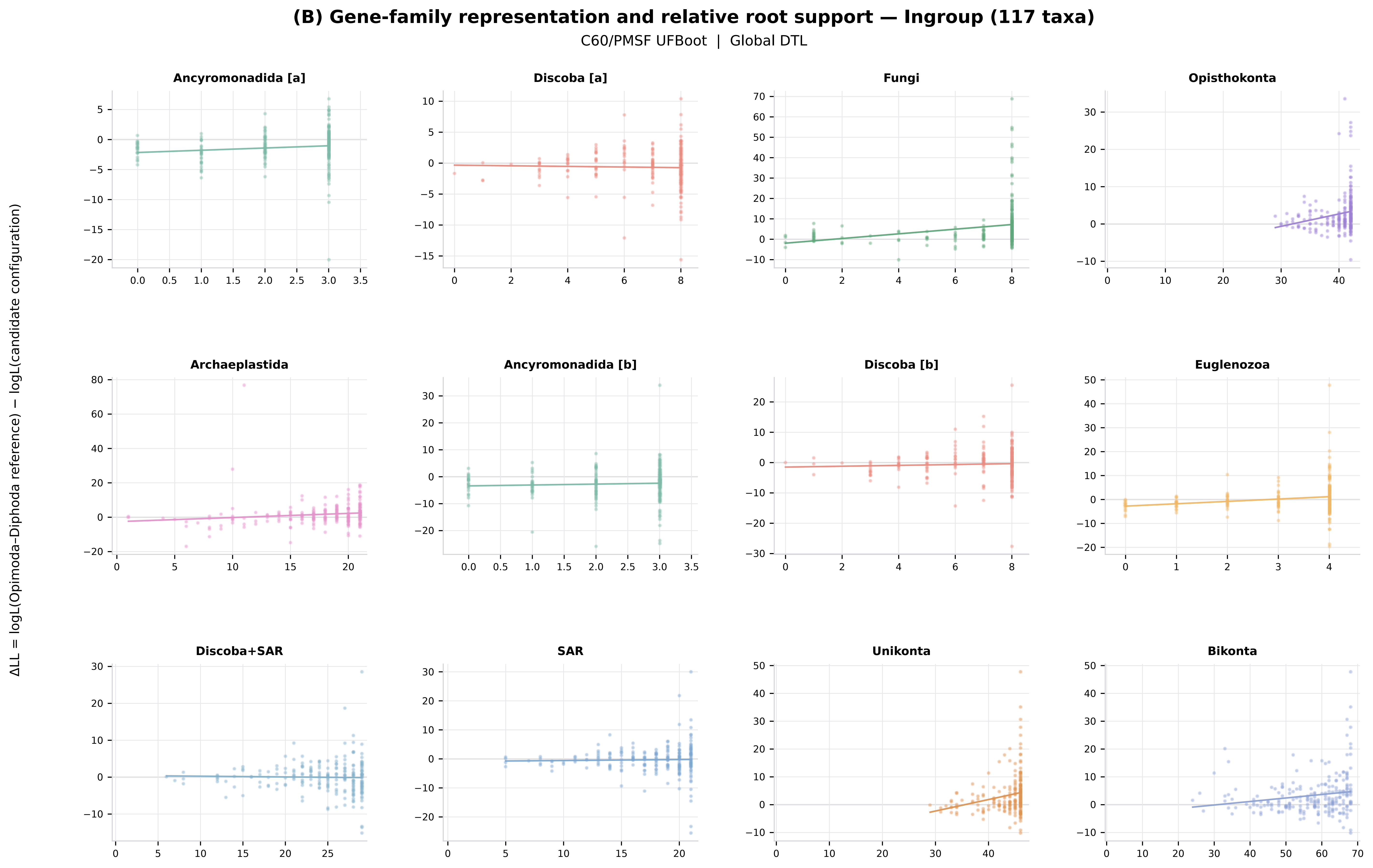

(B) Gene-family representation and relative root support — Ingroup (117 taxa)
C60/PMSF UFBoot | Global DTL
ΔLL = logL(Opimoda–Diphoda reference) − logL(candidate configuration)
Ancyromonadida [a]
Discoba [a]
Fungi
Opisthokonta
Archaeplastida
Ancyromonadida [b]
Discoba [b]
Euglenozoa
Discoba+SAR
SAR
Unikonta
Bikonta

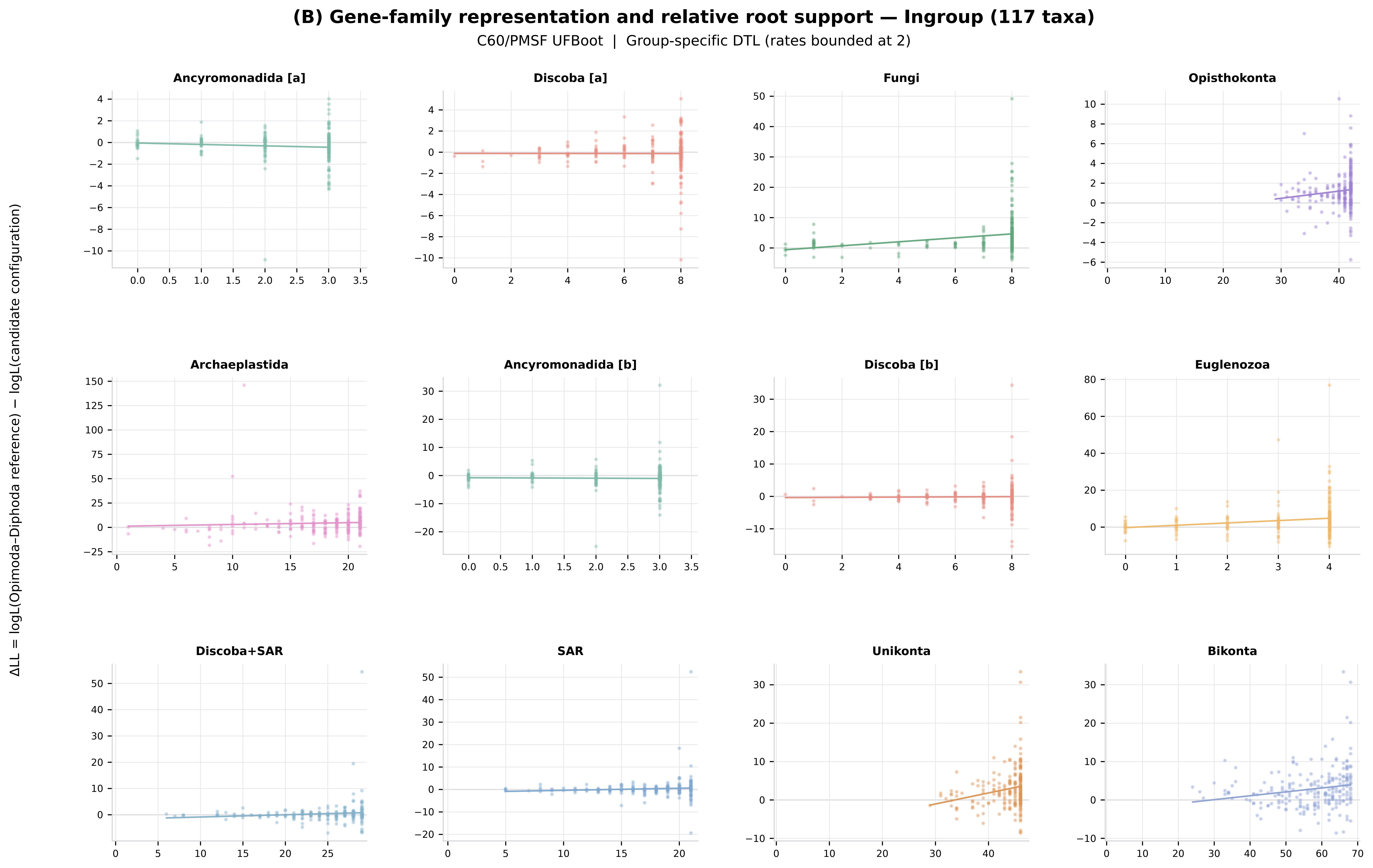
(B) Gene-family representation and relative root support — Ingroup (117 taxa)
C60/PMSF UFBoot | Group-specific DTL (rates bounded at 2)
ΔLL = logL(Opimoda–Diphoda reference) − logL(candidate configuration)
Ancyromonadida [a]
Discoba [a]
Fungi
Opisthokonta
Archaeplastida
Ancyromonadida [b]
Discoba [b]
Euglenozoa
Discoba+SAR
SAR
Unikonta
Bikonta

**(B) Gene-family representation and relative root support — Ingroup (117 taxa)**

C60/PMSF UFBoot | Group-specific DTL (unbounded)

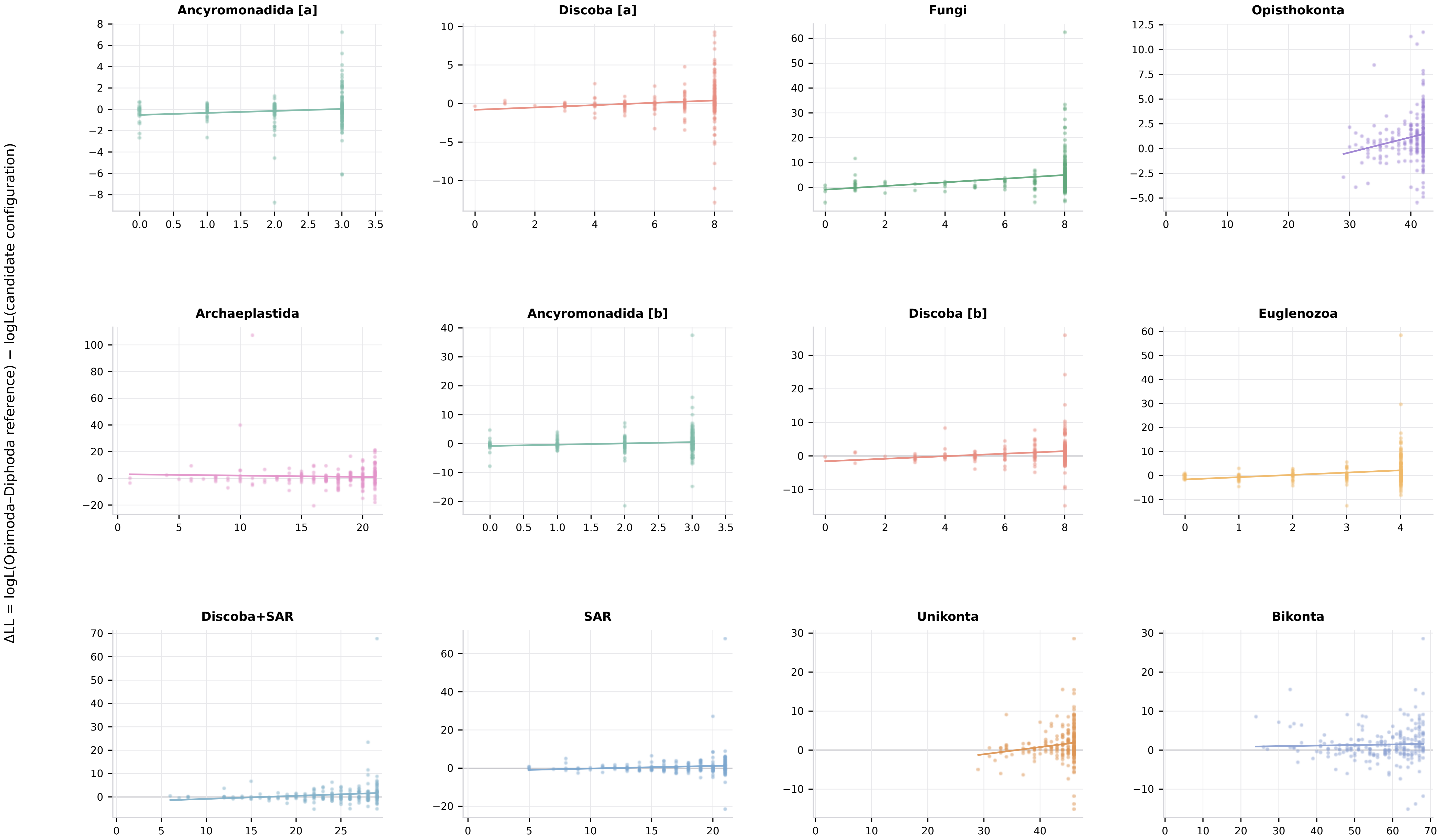
