## Supplementary Note 1 for "Rooting the tree of eukaryotes using ancient gene duplications"

Supplementary Note 1. Sensitivity analyses indicate that root inference is influenced by gene-family absence

To better understand support for alternative rooting hypotheses, we examined gene families according to their relative support for each alternative rooting hypothesis compared with an Opimoda–Diphoda root, by calculating the difference in log-likelihood (ΔLL) between the Opimoda–Diphoda root and the alternative hypothesis for each family using the UFBoot-based reconciliation analyses (Supplementary Table S5). Under this definition, lower or more negative ΔLL values indicate greater relative support for alternative rooting hypotheses over Opimoda–Diphoda. We examined these relationships in both the full dataset (129 taxa) and the reduced ingroup dataset (117 taxa), with the C20–PMSF UFBoot analyses under the branch-homogeneous DTL model shown in Supplementary Fig. S8a,b. This analysis revealed a broad, though not universal, trend: gene families that were absent or had low representation in a lineage tended to favour a root on that lineage. For example, gene families with no or limited representation in Fungi tended to favour a fungal root over the Opimoda–Diphoda root (Supplementary Figs. S8–S16).

This pattern is consistent with expectations under reconciliation models when loss rates are underestimated: when a gene family is absent from a lineage, that absence can be interpreted as evidence that the lineage diverged before the family originated. If gene loss is underestimated, such absences may therefore artefactually strengthen support for placing gene-poor lineages near the root. This effect was observed for several non-metamonad candidate roots, but it was not uniform across all rooting hypotheses. Metamonads exhibit more gene-family absences than most other major sampled eukaryotic clades. In particular, they lack homologues of 19 of the 219 pre-LECA duplicate families, compared with none in Opisthokonta and only one each in Discoba and Viridiplantae (Supplementary Table S1). However, metamonad-related roots did not show a simple absence-driven pattern. The Metamonada and Metamonada-OpiDiph hypotheses, although biologically distinct, were represented by the same rooted topology in the original tree set. We therefore examined two Metamonada-rooted configurations with different species-tree backbones in the revised analyses (Supplementary Fig. S8a). In the UFBoot-based analyses, the relationships between metamonad representation and ΔLL were weak or negative in several comparisons, but varied among rooting configurations and reconciliation models. Thus, support for metamonad-related roots cannot be explained solely by complete metamonad gene-family absence.

Since we would expect underestimation of lineage-specific loss rates to be more severe with branch-homogeneous models, which assume a common loss rate for all lineages, we asked whether this representation-associated effect is also seen in the branch-heterogeneous DTL analyses, in which branches were assigned to 12 predefined major eukaryotic lineage groups, with a separate set of D, T and L rates estimated for each group, as described in the Methods. Comparison of the relationship between ΔLL and lineage representation in gene families revealed that these relationships varied among reconciliation models, with some correlations becoming weaker or changing direction under branch-heterogeneous models (Supplementary Figs. S10–S13 and S15–S16).

This indicates that lineage-specific loss-rate modelling alone is insufficient to remove all representation-associated effects on relative root support. However, the metamonad-related signal was more complex than expected under a simple “small-genome attraction” model. In site-heterogeneous UFBoot analyses, especially those inferred using C20–PMSF or C60–PMSF substitution models, trends in support for metamonad-related roots varied among rooting configurations and reconciliation models, with some relationships being weak rather than reflecting a consistent genome-wide effect of metamonad gene-family absence. Overall, these analyses indicate that uneven gene-family presence and absence is associated with variation in root support, but that support for metamonad-related roots may reflect a combination of genuine phylogenetic signal, lineage representation, and model fit. Comparisons involving different species-tree backbones may additionally reflect differences in underlying topology as well as root placement.
